# *Arsenophonus* ghosts in bed bug genomes: multiple origins and 50 million years of persistence

**DOI:** 10.64898/2026.07.17.739188

**Authors:** Shruti Gupta, Jana Martinů, Ondřej Balvín, Václav Hypša

## Abstract

Although numerous studies have documented horizontal gene transfer (HGT) from bacteria into insect genomes, they have been heavily biased toward the endosymbiont *Wolbachia*. In contrast, comparatively few studies have examined HGT from other symbionts and insect-associated bacteria. Moreover, research on these systems has primarily focused on transferred genes that retain function in the insect host, contributing to metabolism, symbiosis, detoxification, or host adaptation. Consequently, relatively few empirical systems have been available to investigate the persistence, degradation, diversification, and vertical inheritance of apparently non-functional HGT-derived sequences over macroevolutionary timescales. The widespread occurrence of putatively *Arsenophonus*-derived sequences across Cimicidae therefore provides a rare opportunity to extend these investigations beyond the *Wolbachia* model and to examine the long-term evolutionary fate of non-functional HGT-derived DNA across tens of millions of years of host diversification. We show that *Arsenophonus*-derived HGTs have multiple independent origins across Cimicidae but that one major HGT lineage has persisted through diversification of the subfamily Cimicinae for at least 50 million years. Phylogenetic and compositional analyses indicate that an ancestral *Arsenophonus*-derived genomic region has undergone progressive fragmentation, leaving numerous dispersed, apparently non-functional remnants while preserving a clear evolutionary signature. These results extend the study of bacterial HGT beyond the *Wolbachia* model and demonstrate that non-functional symbiont-derived DNA can persist over macroevolutionary timescales.

## Introduction

Horizontal gene transfer (HGT) from bacteria to insects is widespread and can involve the transfer of individual genes, operons, or large chromosomal fragments into host genomes (Husnik and McCutcheon 2018; Li et al. 2022). In some cases, transferred genes become integrated into host physiology and acquire important biological functions, contributing to novel adaptations. Consequently, most studies of bacterial HGT in insects have focused on functional transfers and their role in host evolution. Early studies of *Wolbachia*-derived insertions demonstrated both the widespread occurrence of bacterial-to-host gene transfer and the complex evolutionary dynamics of transferred DNA within host genomes (Kondo et al. 2002). Although *Wolbachia* is the best-studied source of bacterial-to-host transfers in insects, it is not the only symbiont known to contribute genetic material to host genomes. Functional HGTs have also been documented from nutritional symbionts such as *Carsonella, Tremblaya*, and *Moranella*, whose genes have become integrated into host metabolic pathways in psyllids and mealybugs (Sloan et al. 2014, Husnik et al. 2013). These examples show that HGT from symbiotic bacteria is a widespread feature of insect evolution.

However, not all transferred DNA acquires a functional role. Most bacterial sequences integrated into insect genomes appear to be non-functional and may persist long after the original symbiont has disappeared. Such “genomic fossils” provide evidence of past host–symbiont associations even in species that no longer harbour the donor bacterium. A recent example is provided by the woodwasp *Sirex noctilio*, whose genome contains remnants of *Wolbachia*-derived genes despite the apparent absence of *Wolbachia* infection in extant populations (Queffelec et al. 2022). Compared with functional HGTs, the evolutionary fate of such non-functional transferred sequences remains poorly understood. In particular, little is known about their long-term persistence, diversification, and distribution across radiating host lineages.

In *Cimex lectularius*, only a few studies have addressed HGT, showing that transposable elements constitute a substantial fraction of the genome and that numerous sequences of bacterial origin are present (Benoit et al. 2016; Rosenfeld et al. 2016; Miles et al. 2024). Notably, many of these sequences were assigned to the bacterial genus *Arsenophonus*, despite the apparent absence of *Arsenophonus* among currently detected symbionts. During our previous survey of cimicid-associated symbionts, we similarly detected occasional *Arsenophonus*-like sequences in metagenomic datasets, although no *Arsenophonus* genomes were recovered from any of the 13 investigated cimicid species (Hypsa et al. 2025). The origin of these sequences is particularly intriguing given the biology of *Arsenophonus*, one of the most widespread groups of insect-associated bacteria. Members of the genus possess highly dynamic genomes characterized by abundant mobile elements, extensive recombination, and frequent lateral gene transfer (Mouton et al. 2012, Darby et al. 2010, Frost et al. 2020), making them plausible donors of genetic material to host genomes.

The widespread occurrence of putatively *Arsenophonus*-derived sequences across Cimicidae raises two alternative hypotheses regarding their evolutionary origin. One possibility is that the common ancestor of extant cimicids harboured an *Arsenophonus* symbiont that transferred genetic material into the host genome. Following the loss of the symbiont, the transferred sequences were inherited throughout cimicid diversification and persisted in descendant lineages. If these sequences are largely non-functional, however, their persistence over as much as ∼115 million years of cimicid evolution (the estimated crown age of Cimicidae; Roth et al., 2019) would require the retention of recognizable bacterial signatures over an exceptionally long evolutionary timescale. Under neutral evolution, horizontally acquired sequences are expected to accumulate mutations and progressively diverge from their ancestral state (Lawrence and Ochman 1997, Gogarten and Townsend 2005).

Alternatively, the observed sequences may reflect repeated transfers from distinct *Arsenophonus* lineages that independently associated with cimicid hosts throughout their evolutionary history. Yet this scenario raises a different difficulty: recurrent transfers would imply repeated acquisition of multiple *Arsenophonus* symbionts despite the apparent absence of stable *Arsenophonus* infections in extant cimicids. This contrasts with the evolutionary histories of other cimicid symbionts, including *Wolbachia, Sodalis*, and *Symbiopectobacterium*, which show evidence of long-term persistence within multiple cimicid lineages (Hypsa et al. 2025).

In this study, we investigate the evolutionary origin of *Arsenophonus*-derived sequences in Cimicidae and evaluate alternative scenarios explaining their distribution across the family. In doing so, we address a broader question concerning the long-term evolutionary fate of ancient HGT-derived sequences. The widespread occurrence of putatively *Arsenophonus*-derived DNA across Cimicidae provides a rare opportunity to examine whether horizontally transferred bacterial sequences can remain detectable over tens of millions of years of host evolution, potentially preserving evidence of historical host–symbiont associations long after the donor symbionts themselves have disappeared.

## Results and discussion

### Candidate HGTs incorporated in reference genome

Our initial analysis focused on the *Cimex lectularius* Harlain isolate (hereafter *Cl*_Harlain), which provides a high-quality chromosome-level genome and therefore minimizes the risk of assembly artefacts.

Screening with 30,603 *Arsenophonus* BLAST queries identified 5,697 candidate homologous regions in the *Cl*_Harlain genome. Reciprocal back-BLAST analysis confirmed an *Arsenophonus* origin for 473 sequences, corresponding to 169 unique loci after collapsing overlapping duplicate hits (Supplementary data S1). To demonstrate that the identified HGT loci are integrated into the bed bug genome, pairwise alignments were visualized by the approach illustrated in Figure 1. Each alignment comprised two 5-kb fragments. One fragment was extracted from the *Cl_*Harlain genome, with the HGT locus centred in the sequence. The second fragment was extracted from the *Arsenophonus* genome containing the closest homolog of the HGT locus, which was likewise centred within the 5-kb fragment. The majority of these alignments demonstrated an abrupt transition from a well-aligned central HGT region to non-homologous flanking sequence (Figure 1, Supplementary figure S1).

**Figure 1.**
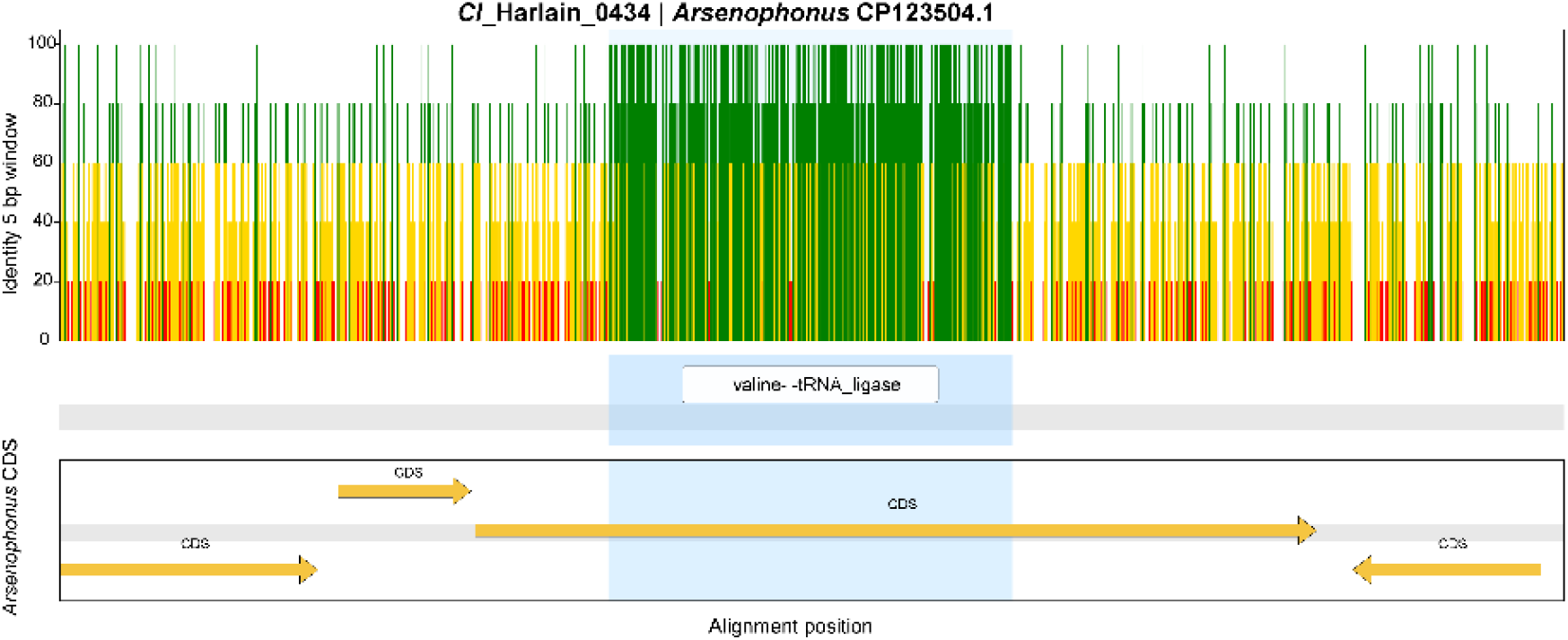
Visualization of an Arsenophonus-derived HGT integrated into the Cimex lectularius Harlain (Cl_Harlain) genome. Cl_Harlain_0434 | Arsenophonus CP123504.1 indicates the HGT candidate identifier and the accession of its closest Arsenophonus homolog. The upper panel shows sequence similarity between two 5-kb genomic regions calculated in 5-bp sliding windows. The central region of high similarity (green island) corresponds to the horizontally transferred fragment, whereas the flanking sequences show no detectable homology, indicating integration of the HGT into the bed bug genome. The complete set of 169 alignments is shown in **Supplementary Figure S1**.

The distribution of *Arsenophonus*-like fragments in the *Cl*_Harlain genome is significantly non-random, as evidenced by short-range clustering that is much stronger than expected under random placement. Specifically, 19 loci occur in multi-locus clusters within 10 kb compared with a random expectation of 1.04, 21 loci cluster within 50 kb compared with 5.17 expected, and 27 loci cluster within 100 kb compared with 10.23 expected. This pattern may reflect either host genome effects, whereby certain genomic regions are more permissive to HGT retention or detection, or the fragmentation of larger ancestral *Arsenophonus*-derived insertions through deletion and genomic rearrangement. Given the phylogenetic evidence described below, we consider the latter explanation more likely for at least part of the *Cl_*Harlain signal.

The complete *Cl*_Harlain-associated HGT set shows a consistent GC content centered at 38–39%, with a few short, low-GC fragments extending the lower tail. GC content is strongly correlated with matched *Arsenophonus* sequences (Pearson’s *r* = 0.87; Supplementary figure S2), indicating retention of the bacterial compositional signature, although *Cl*_Harlain fragments are on average ∼1.6 percentage points GC-poorer. This difference is only weakly related to fragment length (*r* = ™0.13), but the most extreme low-GC outliers are short fragments. Excluding loci <150 bp substantially narrows the distribution and eliminates potential multimodality, suggesting that it is driven by a few short, compositionally unstable fragments rather than distinct GC classes. These results are consistent with acquisition of a larger *Arsenophonus*-derived region that subsequently fragmented over evolutionary time.

Gene-content analysis showed that the 169-locus candidate HGT set is functionally heterogeneous rather than dominated by a single gene family, mobile element, or functional module. Although several annotations recurred, no single annotation predominated. The most common annotations were *hypothetical protein, gorA, ubiD*, and *hslV*, whereas all others were represented by only one or two loci. The observed recurrence is consistent with fragmentation of ancestral *Arsenophonus*-derived material, repeated assignment of short conserved fragments to the same bacterial genes, or paralogy within the donor genome. Functional categories included central, amino acid, and lipid metabolism; cell envelope and division; transport and membrane proteins; protein folding and proteolysis; DNA replication and repair; transcription and translation; cofactor and vitamin metabolism; and hypothetical or uncharacterized genes. Only a small fraction of fragments were annotated as mobile- or recombination-related genes, although some loci showed mobile-element signals in the broader NCBI genomic context. Overall, the *Cl*_Harlain-associated fragments are best interpreted as a dispersed, degraded collection of ordinary bacterial gene fragments rather than a mobile-element-dominated insertion or a coherent functional cassette. This pattern is consistent with fragmentation of ancestral *Arsenophonus*-derived material. The gene content of this candidate HGT set is summarized in Supplementary table S1.

### Functional comparison between the reference and HGT candidates

Because phylogenetic analyses indicated that at least a substantial subset of the *Cl*_Harlain-associated HGT loci originated from a single evolutionary event, we compared their functional composition with that of 30 representative *Arsenophonus* genomes (see Methods). The aim was to determine whether the frequencies of functional gene categories retained by HGT resembled the functional profiles of facultative or obligate *Arsenophonus* symbionts. The PCoA analysis revealed clear structure among the taxa, with PCoA1 and PCoA2 accounting for 67.63% and 8.96% of the total variation, respectively, indicating that most functional variation was captured along a single dominant axis. The genomes formed two major functional clusters. Functional Cloud 2 was enriched for obligate and co-obligate symbionts, whereas Functional Cloud 1 contained a greater proportion of facultative lineages (Figure 2, Supplementary table S2), although several notable exceptions indicate that the relationship between functional profile and the current type of symbiosis is not one-to-one.

**Figure 2.**
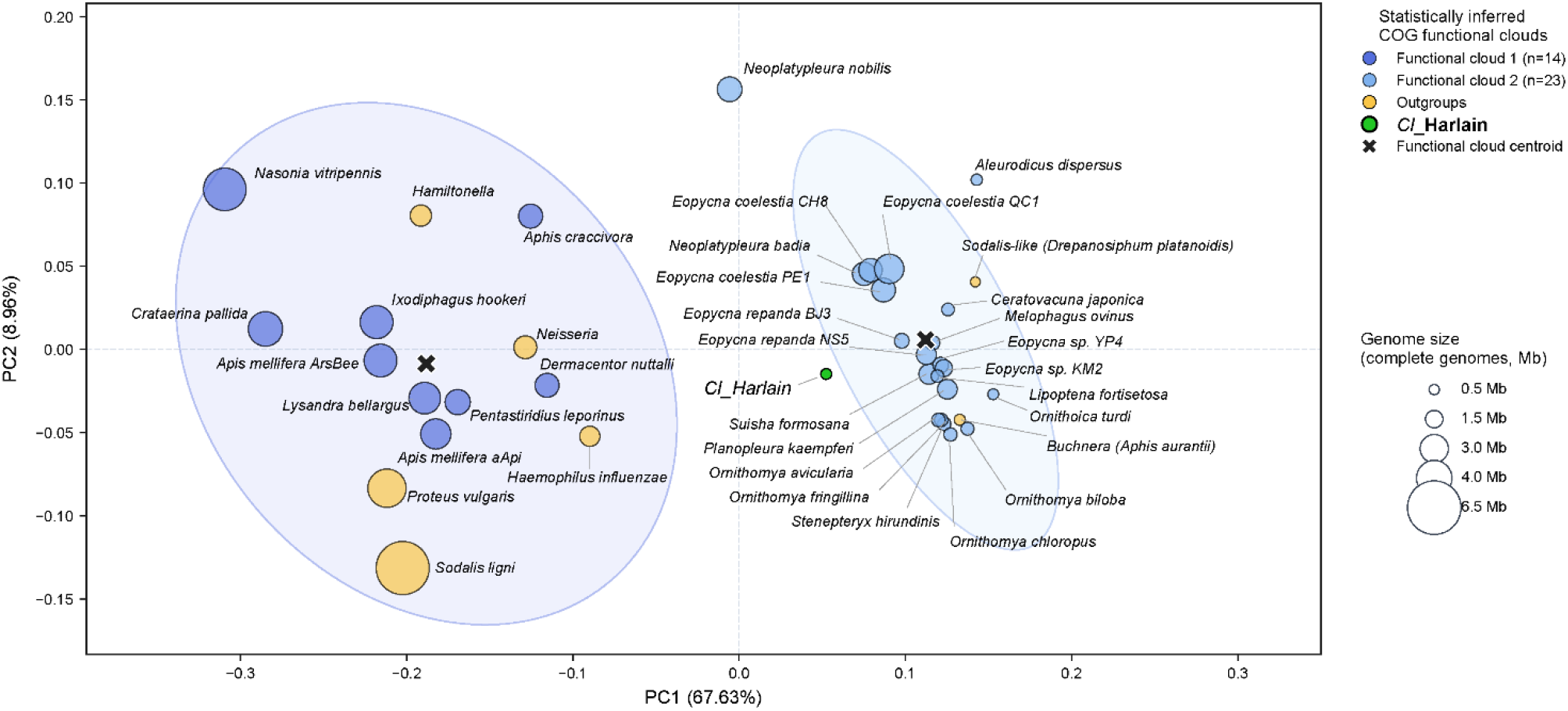
Functional comparison of the Cl_Harlain HGT repertoire and representative bacterial genomes. Principal coordinates analysis (PCoA) shows that the functional composition of the Cl_Harlain HGT repertoire is most similar to that of Arsenophonus genomes occupying Functional Cloud 2, which is enriched in obligate symbionts. This pattern is supported by pairwise Bray–Curtis nearest-neighbour analysis: the closest functional neighbour of the Cl_Harlain HGT repertoire was the Arsenophonus symbiont of Suisha formosana (Bray–Curtis distance = 0.1143), and all ten nearest neighbours belonged to Functional Cloud 2. Dark and light blue symbols represent Arsenophonus genomes, with labels indicating their host species. Yellow symbols represent the outgroup genomes. Details of all genomes are provided in Supplementary table S2.

Along PC1, the HGT gene set occupied functional space closer to Functional Cloud 2 than to Cloud 1 (centroid distance 0.0635 versus 0.2397). Because the analysis was based on normalized COG proportions, this relationship reflects similarity in functional composition rather than genome size. PC1 primarily separated translation-, ribosome-, and biogenesis-associated functions (COG J; ρ = 0.9527), characteristic of Cloud 2, from genes of unknown function (COG S; ρ = ™0.9341), characteristic of the opposite end of the ordination. Consistent with this pattern, the *Cl*_Harlain HGT repertoire showed an overall COG composition resembling Cloud 2, including similar representation of coenzyme metabolism (COG H), cell wall/membrane/envelope biogenesis (COG M), and post-translational modification, protein turnover, and chaperones (COG O). These results indicate that the transferred repertoire resembles the multivariate functional profile of Cloud 2 rather than being enriched for a single functional category.

PC2 accounted for only 8.96% of the total variation and primarily reflected a contrast between replication, recombination and repair (COG L; ρ = 0.7991) and amino acid transport and metabolism (COG E; ρ = ™0.8015). *Cl*_Harlain occupied a central position within Cloud 2 along this axis, indicating little deviation from the characteristic functional profile of the cluster. Together, the PCoA placement, centroid distances, nearest-neighbour Bray–Curtis analysis, and COG-category associations indicate that the *Cl*_Harlain HGT repertoire represents a biologically structured subset of *Arsenophonus*-derived genes. Although these analyses do not identify the direct donor lineage, they show that the retained HGT repertoire most closely resembles the functional composition of the *Arsenophonus* genomes occupying Functional Cloud 2, which predominantly comprises obligate symbionts.

### Origin and evolution of Arsenophonus sequences across Cimicidae

We analyzed 14 genome assemblies representing 13 Cimicidae species (Hypsa et al. 2025) (Supplementary table S3) to address two key questions. First, do the *Arsenophonus* sequences identified across Cimicidae share a single common origin, or do they result from multiple independent HGT events? Second, is there evidence that these sequences have persisted throughout host diversification?

To investigate these questions, we combined two complementary approaches: GC-content analysis, which served as an initial screen to assess the consistency of individual HGT sets, and phylogenetic analyses to place them in an evolutionary context. GC-content screening revealed considerable heterogeneity among Cimicidae samples, consistent with the possibility of multiple independent HGT origins. Within individual samples, most HGT sets were characterized by a single dominant GC class. The only exceptions were C57-*Cimex lectularius* H and C51-*Cimex hirundinis*, whose GC distributions appeared weakly bimodal. In both cases, the secondary component was small and was reduced or disappeared after excluding loci shorter than 150 bp (Figure 3).

**Figure 3.**
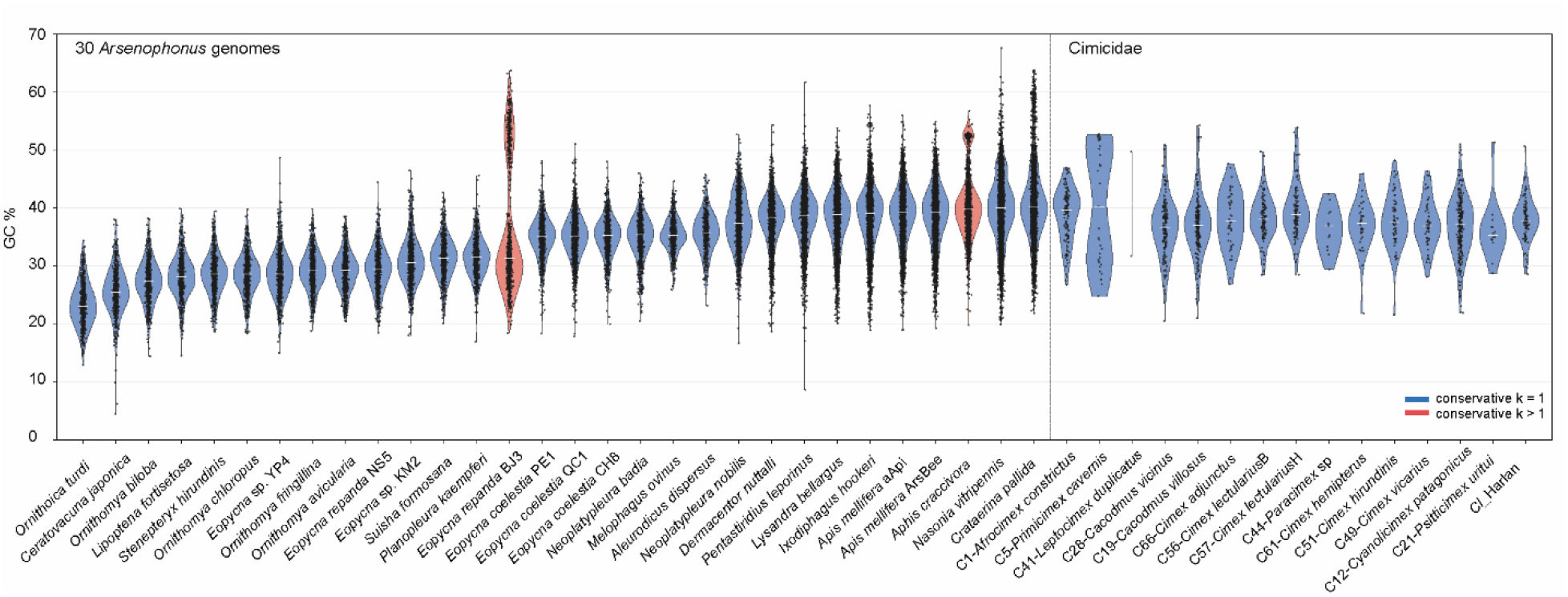
GC-content distributions for reference Arsenophonus genomes and Cimicidae-derived HGT loci after excluding sequences shorter than 150 bp. Violin plots show the distributions of GC percentages for coding sequences (reference genomes) or retained HGT loci (Cimicidae samples), with individual loci indicated by black dots. Genome and sample labels follow the nomenclature used in Figure 2 (Supplementary table S2).

Once the HGT sets were confirmed to be GC-consistent, we placed them in a phylogenetic framework. In the first approach, candidate HGTs were mapped to their best matches on a reference *Arsenophonus* phylogeny, revealing a pattern consistent with multiple independent origins of HGTs across Cimicidae (Figure 4A). Furthermore, the clustering of HGTs on the reference tree broadly reflected the phylogenetic relationships among bed bug hosts, suggesting that individual HGT lineages have been retained through subsequent host diversification. As shown in Figure 4A, HGT loci within each GC-consistent set produced best matches to multiple *Arsenophonus* reference genomes rather than to a single strain. This pattern most likely reflects the use of short, often fragmented HGT sequences and the high sequence similarity among closely related *Arsenophonus* genomes, which results in best hits being distributed among several reference strains. Accordingly, these mappings were interpreted as measures of phylogenetic affinity rather than direct donor assignments. More informative than the individual best hits was the overall distribution of matches: each HGT set was characterized by a dominant reference genome and a similar spectrum of secondary matches. This pattern was especially pronounced in the six species of the subfamily Cimicinae, whose closely similar best-hit profiles provide additional support for a shared origin of their HGT sets.

**Figure 4.**
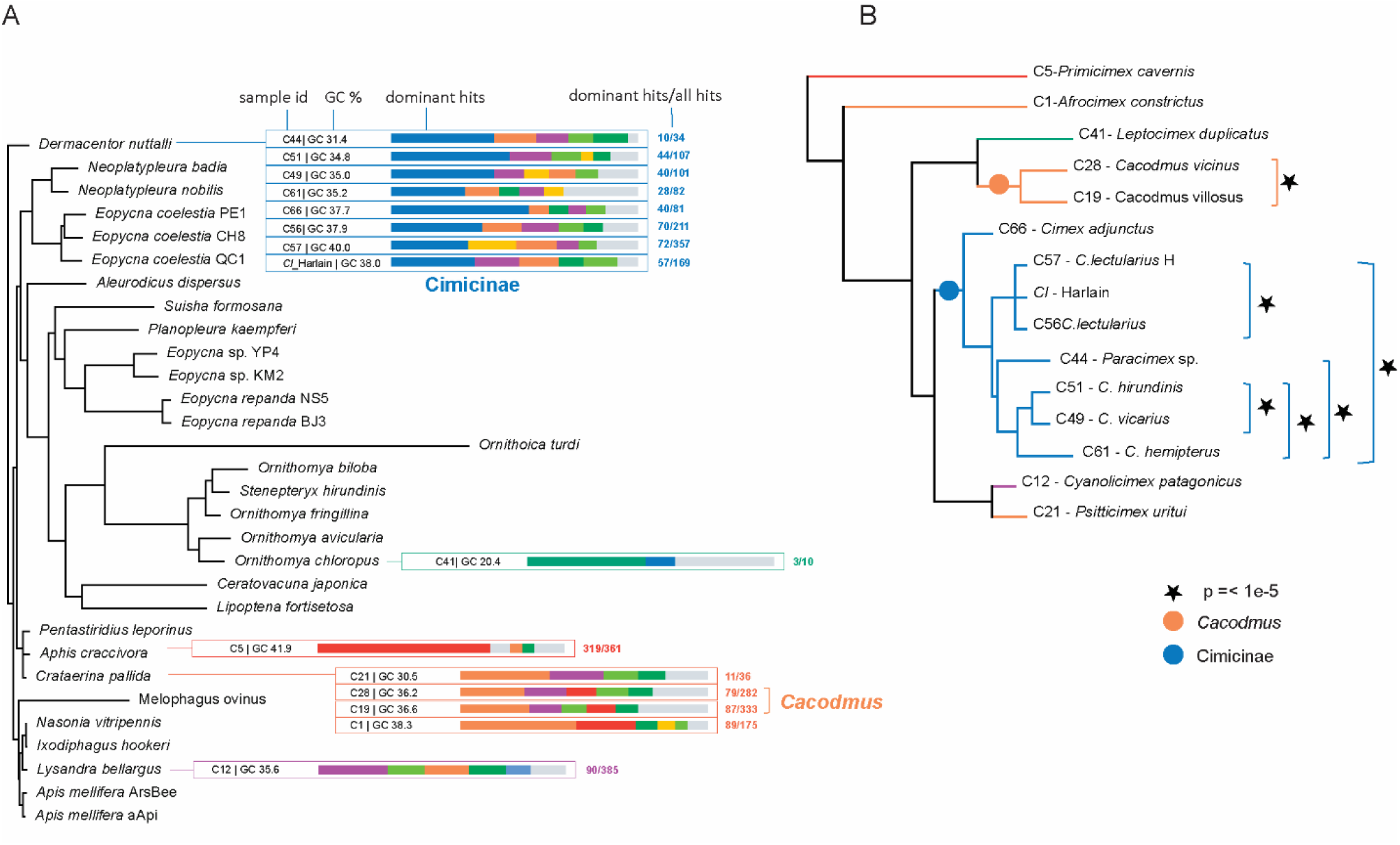
Evolutionary origin and persistence of Arsenophonus-derived HGTs across Cimicidae. (A) Best-hit analysis. The tree shows the phylogeny of 30 representative Arsenophonus genomes (Supplementary Table S2) together with the best-hit profiles of HGT sets from individual cimicid species. The distribution of best hits supports multiple independent origins of Arsenophonus-derived HGTs and a shared HGT lineage among the Cimicinae species. (B) Bridge-compatibility analysis. The Cimicidae phylogeny is adapted from Hypsa et al. (2025), with sample colours corresponding to those in panel A. Statistically supported bridge-compatible host clades are indicated on the right. The Cacodmus clade was recovered as bridge-compatible in 31 of 31 informative trees (P = 3.66 × 10^−52^), whereas the core Cimicinae clade was recovered in 30 of 34 informative trees (P = 4.38 × 10^−121^).

To test the coevolutionary pattern suggested by the preceding analysis, we applied a second phylogenetic approach. Using homolog sets for the HGT sequences, 30 *Arsenophonus* genomes, and four outgroups, we reconstructed phylogenetic trees for each retained representative HGT locus. Bridge-compatibility analysis of 238 maximum-likelihood trees identified two host clades that were recovered far more often than expected under the random-topology null model (Figure 4B, Supplementary figure S3). The first comprised the two *Cacodmus* species, the sub-Saharan C19-*Cacodmus villosus* and the Mediterranean C28-*Cacodmus vicinus*. This pair was bridge-compatible in all 31 informative trees, compared with a random expectation of 0.78 compatible trees (exact Poisson-binomial upper-tail P = 3.66 × 10^-52). The second, and biologically most notable, was a core Cimicinae subfamily clade comprising seven samples from five species: C56-*Cimex lectularius* B, C57-*Cimex lectularius* H, *Cl*_Harlain, C44-*Paracimex* sp., C61-*Cimex hemipterus*, C51-*Cimex hirundinis*, and C49-*Cimex vicarius*. This core Cimicinae group was bridge-compatible in 30 of 34 informative trees, compared with a random expectation of 0.05 compatible trees (exact Poisson-binomial upper-tail P = 4.38 × 10^-121). The basal Cimicinae sample C66-*Cimex adjunctus* was not recovered within this bridge-compatible cluster, although its affinity to the Cimicinae-associated HGT lineage is supported by the best-hit mapping analysis (Figure 4A). This discrepancy is most likely explained by the smaller number of informative homologous loci retained in C66-*Cimex adjunctus*, reducing its recovery under the more stringent bridge-compatibility framework. A third host-compatible pair, C12-*Cyanolicimex patagonicus*+C21-*Psitticimex uritui*, was represented by only one informative tree and was not bridge-compatible (0/1; P = 1). The remaining samples, including C1-*Afrocimex constrictus*, C5-Primicimex cavernis, and C41-Leptocimex duplicatus, showed no comparable host-phylogeny-compatible signal and instead were associated with different regions of the broader *Arsenophonus* phylogenetic context. Together, these results support multiple independent origins of *Arsenophonus*-derived HGTs across Cimicidae while demonstrating long-term persistence of one HGT lineage throughout diversification of the core Cimicinae.

The phylogenetic distribution of this shared Cimicinae HGT lineage is consistent with acquisition in the common ancestor of the core Cimicinae followed by vertical inheritance during subsequent diversification. This scenario closely parallels the acquisition and long-term coevolution of the *Wolbachia* symbiont across the same species group (Hypsa et al., 2025). Given the estimated divergence of *Cimex lectularius* and *C. hemipterus* approximately 47 Mya (Roth et al. 2019), the shared *Arsenophonus*-derived sequences must have persisted in Cimicinae genomes for at least 50 million years after their acquisition. Although long-term persistence of symbiont-derived DNA has been documented for *Wolbachia*-derived insertions (Nikoh et al., 2008; Klasson et al., 2009; Tvedte et al., 2022), few studies have traced the evolutionary history of apparently non-functional bacterial HGT fragments across a well-resolved host radiation.

### The main conclusions

This study revealed two notable patterns in the distribution of *Arsenophonus*-derived HGTs in bed bug genomes. First, *Arsenophonus*-derived sequences were detected across all sampled Cimicidae species despite the absence of any known extant *Arsenophonus* symbiont in bed bugs. Together with the multiple independent origins of the extant *Wolbachia, Symbiopectobacterium*, and *Sodalis* symbioses (Hypsa et al., 2025), this finding further supports the remarkable tendency of Cimicidae to repeatedly establish associations with a restricted pool of bacterial lineages.

Second, *Arsenophonus*-derived HGTs exhibit exceptional evolutionary persistence within bed bug genomes. This is most clearly demonstrated by the monophyletic group of *Arsenophonus*-derived sequences shared by Cimicinae species, whose phylogenetic placement repeatedly supports the monophyly of the corresponding host clades. We therefore propose that an *Arsenophonus* genomic region was horizontally transferred into the genome of the common ancestor of Cimicinae at least ∼50 million years ago and subsequently inherited by these descendant lineages. During this time, repeated genomic rearrangements and deletions fragmented the original insertion, leaving numerous short, apparently non-functional remnants dispersed throughout the genome. The persistence of these HGT-derived fragments over such an extended evolutionary timescale exceeds that documented for most non-functional bacterial insertions. These findings raise the possibility that similarly ancient HGT fossils have been overlooked in other insect genomes, although they may also reflect an unusual capacity of bed bug genomes to retain horizontally acquired DNA over evolutionary timescales.

## Methods

### General workflow

To identify and characterize candidate *Arsenophonus*-derived sequences in bed bug genomes, we used the following multi-step workflow: (i) an *Arsenophonus*-derived query dataset was used to identify homologous sequences in 14 previously published Cimicidae genome assemblies generated from Illumina short-read data (Hypsa et al., 2025), together with the chromosome-level reference genome of *Cimex lectularius* (Harlain isolate; ASM4990377v1; hereafter *Cl*_Harlain), (ii) the retrieved sequences were searched against the NCBI nt database to confirm their *Arsenophonus* origin; (iii) candidate HGT loci identified in *Cl*_Harlain were mapped onto the *Cl*_Harlain genome to verify that they represent genuine *Arsenophonus*-to-bed bug horizontal gene transfer (HGT) events; and (iv) the resulting HGT candidates for all screened cimicids were characterized by analyses of functional gene content, GC content, and phylogenetic relationships to assess their evolutionary origin and history. These steps are described in detail below.

### Identification of HGT candidates in bed bug genomes

#### Identification

We generated an *Arsenophonus* query dataset by extracting all unique coding sequences (CDSs) from 17 selected *Arsenophonus* genomes (Supplementary table S4), representing both gene-rich strains and obligate symbionts with reduced genomes. The *Cl*_Harlain genome and 14 previously published assemblies were screened for homologs to the 30,603 unique *Arsenophonus* CDSs using BLAST (Altschul et al. 1990, Camacho et al. 2009) with blastn algorithm (max_target_seqs = 20). To confirm *Arsenophonus* origin, all retrieved candidate hits were subjected to a reciprocal “back-BLAST” analysis against the NCBI nt database (blastn, evalue 1e-5, max_target_seqs 1).

#### Deduplication

Initial BLAST screening frequently recovered the same genomic locus multiple times because different *Arsenophonus* query sequences, or overlapping query fragments, matched the same genomic region with slightly different coordinates. Before final locus definition, records located on curated problematic contigs, including phage-associated, mobile-element-rich, and high-GC bacterial-island-like contigs, were excluded from the Cimicidae dataset. This retained 25,796 Cimicidae sequence-level records. These records were then collapsed to locus-level representatives by clustering hits on the same contig and strand that showed reciprocal overlap of their genomic intervals, allowing a 10-bp tolerance for small boundary differences. Within each cluster, a single representative was retained based on the highest BLAST bitscore, followed by the highest percentage identity and, if necessary, stable record ID. This produced 2,555 non-redundant Cimicidae HGT loci. The *Cl*_Harlain dataset was processed separately through an nt-confirmed intermediate set of 473 unique *Arsenophonus*-confirmed sequences, from which 169 final *Cl*_Harlain HGT loci were retained. Downstream analyses therefore used 2,555 Cimicidae loci and 169 *Cl*_Harlain loci, for a combined total of 2,724 unique HGT loci (Supplementary data S1).

### Integration of HGT sequences within the reference genome

To verify that these sequences were located within the bed bug genome, we used an additional alignment-visualization approach. For this step, we focused on *Cl_*Harlain chromosome-level genome to avoid potential artefacts associated with the 14 short-read assemblies. For each of the 169 back-BLAST-confirmed *Arsenophonus*-derived loci in the *Cl*_Harlain genome, we extracted the corresponding bed bug sequence together with 5 kb of flanking sequence on each side, and aligned it to the matching *Arsenophonus* sequence with equivalent 5 kb context. Sequence similarity across each alignment was then visualized using a 5 bp sliding window. The resulting plots allowed us to distinguish *Arsenophonus*-derived HGT fragments from the surrounding host genomic regions and to assess their genomic context (Supplementary figure S1).

### Functional characterization of the HGT repertoire

The aim of this analysis was to determine whether the frequencies of functional gene categories retained by HGT resembled the functional profiles of facultative or obligate *Arsenophonus* symbionts. Functional characterization was based on eggNOG-derived Cluster of Orthologous Groups (COG) classifications (Galperin et al. 2021). COG annotations were assigned with eggNOG-mapper (Cantalapiedra et al. 2021) to the 169 *Cl_*Harlain HGT candidate coding sequences (Supplementary table S1), 30 reference *Arsenophonus* genomes, and seven outgroup genomes. Sequences assigned to multiple COG categories retained all assignments, whereas sequences lacking a COG annotation were classified as unassigned. For each dataset, COG-category counts were converted to within-dataset relative abundances. Functional dissimilarity among datasets was quantified using Bray-Curtis distances (Bray and Curtis 1957) calculated from these relative-abundance profiles, and Principal Coordinates Analysis (PCoA) was used to visualize relationships among functional profiles. Associations between individual COG categories and PCoA axes were evaluated using Spearman rank correlations. Reference functional clusters were inferred only from the 37 complete bacterial genomes, comprising 30 *Arsenophonus* genomes and seven outgroups; the *Cl_*Harlain HGT repertoire was excluded from cluster inference. Reference-genome functional profiles were clustered by average-linkage hierarchical clustering on the Bray-Curtis distance matrix. The resulting hierarchy was cut into k = 1 to k = 6 clusters, and these alternative cluster solutions were evaluated using the silhouette coefficient calculated from Bray-Curtis distances, and the Calinski-Harabasz and Davies-Bouldin indices calculated from PC1/PC2 coordinates (Caliński and Harabasz 1974, Davies and Bouldin 1979). The optimal solution was selected by the lowest combined rank across these criteria. *Cl_*Harlain was then evaluated relative to the established clusters by Euclidean distance to cluster centroids in PC1/PC2 space and by pairwise Bray-Curtis dissimilarities to individual bacterial genomes (Supplementary table S2).

### Evolutionary analysis across Cimicidae: GC content

The analyses were designed to address two related questions: whether *Arsenophonus*-like sequences in Cimicidae (*Cl*_Harlain plus the remaining Cimicidae assemblies) are compatible with a single origin or instead represent multiple independent acquisitions, and whether any of the sequences show host-phylogenetic structure consistent with persistence through host diversification. We used two complementary approaches. First, GC content was used as a composition-based screen for heterogeneity among and within Cimicidae samples. This analysis was intended as an initial test of whether the selected *Arsenophonus*-like sequence pool contained one or more compositional classes. Because GC composition is influenced by gene-to-gene variation, fragment length, and donor genome characteristics, differences in GC composition were interpreted descriptively rather than as direct evidence of donor identity or evolutionary origin. Second, phylogenetic analyses were used to place the candidate HGT sequences in the context of *Arsenophonus* diversity and the Cimicidae host phylogeny. GC analyses were performed on the final non-redundant HGT locus set after collapse of overlapping BLAST-derived records to locus-level representatives. The analysed dataset comprised 2,724 loci, including the 169 *Cl*_Harlain loci (Supplementary data S1). For each locus, fragment length and GC content were calculated, and for each Cimicidae sample we summarized the number of loci, mean GC, median GC, GC range, and GC distribution. Across-sample GC distributions were compared descriptively, whereas within-sample heterogeneity was evaluated using the empirical *Arsenophonus* reference baseline described below.

To establish a conservative empirical expectation for GC heterogeneity within single bacterial genomic backgrounds, we analysed CDS GC distributions from 30 representative *Arsenophonus* genomes after excluding short loci (<150 bp). One-dimensional Gaussian mixture models were fitted to CDS GC values for each reference genome, and one- and two-component models were compared using the Bayesian Information Criterion (BIC). Because weak apparent multimodality can arise within a single bacterial genome through ordinary gene-to-gene compositional variation, BIC support for two components was not interpreted directly as evidence for multiple compositional classes. Instead, the reference genomes were used to define an empirical single-genome range of GC heterogeneity. The two strongest outliers, *Arsenophonus* strains from *Eopycna repanda* BJ3 and *Aphis craccivora*, were treated as multiple-component reference genomes, whereas all remaining references were retained within this range. The operational threshold was defined by the most extreme reference genome still retained within the empirical single-genome range, genome of *Arsenophonus* from *Crataerina pallida*, with ΔBIC = BIC(k = 2) ™ BIC(k = 1) = ™338.13. Evidence for multiple GC components within individual Cimicidae samples was accepted only when heterogeneity exceeded this threshold. Final GC classifications were therefore based on the length-filtered data and their position relative to this empirical reference range, rather than on the nominal best-fitting mixture model alone (Supplementary table S5). For *Cl*_Harlain set, we used these data also to estimate GC differences between the HGT set and the corresponding *Arsenophonus* homologues.

### Analysis of evolutionary origin(s) across Cimicidae based on Arsenophonus reference phylogeny

To identify putative origins of the HGT sequences, we applied two reciprocal phylogeny-based approaches; one using reference phylogeny of the 30 representative *Arsenophonus* genomes (see above), the other reference phylogeny of Cimicidae species (Hypsa et al. 2025). The aim of the first approach was to place the GC classes determined above into an evolutionary context. We used a reference phylogeny constructed from the 30 representative *Arsenophonus* genomes and four outgroup taxa (*Neisseria, Haemophilus, Proteus*, and *Providencia*). Protein-coding genes from the 34 genomes were clustered into orthogroups using OrthoFinder (Emms and Kelly 2019). Single-copy orthogroups were aligned individually and concatenated into a core-protein alignment, from which a maximum-likelihood phylogeny was inferred using IQ-TREE2 (Minh et al. 2020).

Each retained Cimicidae HGT candidate was searched against a local database of the 30 *Arsenophonus* genomes using BLASTN (blastn, -evalue 1e-5, -max_target_seqs 100). For each query, the best-matching genome was identified using highest bitscore, with percent identity and query coverage used to resolve ties. The best-hit genome of every retained HGT locus was recorded and summarized for each sample (Supplementary table S6). These summaries were mapped onto the 30-genome *Arsenophonus* reference phylogeny. In the visualization, each callout represents a final GC class within a Cimicidae sample and is connected to its predominant best-hit genome on the reference tree. The stacked bar shows the base-pair-weighted distribution of best-hit genomes within that class, whereas the accompanying label reports the number of loci assigned to the predominant best-hit genome relative to the total number of loci in the class. These mappings were interpreted as descriptive measures of phylogenetic affinity rather than direct donor assignments. Short or degraded HGT fragments may match conserved regions shared among multiple *Arsenophonus* genomes, and the available references represent only a subset of the diversity within the genus. Accordingly, best-hit mapping was used to visualize whether HGT candidates from a given host sample preferentially matched particular regions of *Arsenophonus* diversity, whereas inference of HGT evolution across Cimicidae diversification relied primarily on phylogenetic analyses of individual HGT loci.

### Analysis of evolutionary origin(s) across Cimicidae based on Cimicidae reference phylogeny

In this second phylogenetic approach, phylogenetic relationships were reconstructed separately for individual HGT candidate loci to test whether independent *Arsenophonus*-like fragments repeatedly recovered host-phylogeny-compatible groupings, as expected if transferred fragments had been inherited during diversification of Cimicidae lineages. Homologous sequence sets were obtained using each HGT candidate as a BLASTN query. Query sequences were searched with BLAST (blastn, evalue 1e-5, max_target_seqs 1) against local nucleotide databases comprising the Cimicidae datasets (including *Cl*_Harlain), 30 representative *Arsenophonus* genomes, and four bacterial outgroups: *Neisseria, Haemophilus, Proteus*, and *Providencia*. The best hit from each database was retained, and hits from all databases were combined into a query-centred homologous sequence set. Because the same HGT locus could be recovered by more than one query sequence, redundant query-centred homolog sets were collapsed before alignment so that only one representative homologous sequence set was retained per non-redundant HGT locus. Records shorter than 100 bp were removed before alignment. Representative homologous sequence sets were retained for phylogenetic analysis only when, after this pruning, they contained at least four sequences and at least one outgroup sequence. This produced 238 representative homologous sequence sets. These sets were aligned with MAFFT (--auto) (Katoh and Standley 2013), trimmed with ClipKIT (smart-gap) (Steenwyk et al. 2020), and used for independent IQ-TREE2 inference under the GTR+G nucleotide substitution model. Trees were rooted using the first available outgroup in the order *Neisseria, Haemophilus, Proteus*, and *Providencia*. The resulting trees were used to score for the compatibility between the HGT and host phylogenies.

### Statistical evaluation of Cimicidae-phylogeny-compatible HGT clades

To distinguish support for a phylogeny backbone node from support arising solely from nested descendant clades, a “bridge-compatibility” criterion was applied. For each tested node of the Cimicidae reference phylogeny, the taxa descending from its immediate child branches defined the bridge partitions of that node (Supplementary figure S3A). Because the reference phylogeny is bifurcating, each node comprised two bridge partitions, except for the unresolved C57-*Cimex lectularius* H+C56-*Cimex lectularius* B+ *Cl*_Harlain trichotomy, which comprised three partitions. Tree manipulation, MRCA identification, and descendant-tip extraction were performed in R using the ape package v5.8.1 (Paradis and Schliep 2019). A tree was considered informative only when it contained sampled HGT loci from at least two bridge partitions of the tested node (Supplementary figure S3B). Compatibility was then assessed by strict monophyly: the sampled HGT sequences from the tested host group had to form an exclusive clade containing no additional Cimicidae sample, *Arsenophonus* reference sequence, or outgroup taxon. This criterion reflects the expectation that HGT loci inherited by vertical descent should remain monophyletic relative to their bacterial homologues.

Statistical support for host-phylogeny-compatible HGT groups was evaluated under a uniform random rooted binary topology null model. For each tested group, only bridge-informative trees were included, i.e. trees containing sampled HGT loci from at least two immediate descendant partitions of the corresponding node in the Cimicidae backbone. For each bridge-informative tree with N total tips and k sampled tips from the tested group, we calculated the exact probability that those k tips would form a monophyletic clade by chance under a random rooted binary topology, following the random-branching framework for tests of monophyly (Rosenberg 2007). Because bridge-informative trees differed in taxon sampling, the resulting per-tree probabilities were combined using a Poisson-binomial upper-tail test, which gives the probability of observing at least the observed number of bridge-compatible trees across the informative tree set (Hong 2013). These exact Poisson-binomial upper-tail probabilities were used as the reported P-values. Tree manipulation, MRCA identification, and descendant-tip extraction were performed in R using ape v5.8.1 (Paradis and Schliep 2019). As a diagnostic check on the same null model, we also generated 100,000 simulated null replicates using the per-tree monophyly probabilities; these simulations were used only to summarize the expected null distribution, not as the source of the reported P-values.

## Acknowledgments

Part of the computational resources were provided by the e-INFRA CZ project (ID:90254), supported by the Ministry of Education, Youth and Sports of the Czech Republic. This work was supported by the Grant Agency of the Czech Republic (grant number 24-10943S to V.H.).

## Ethics declarations

### Ethics approval and consent to participate

Not applicable.

