## Supplementary figure S1 for "*Arsenophonus* ghosts in bed bug genomes: multiple origins and 50 million years of persistence"

CI\_Harlain\_0434 | Arsenophonus CP123504.1

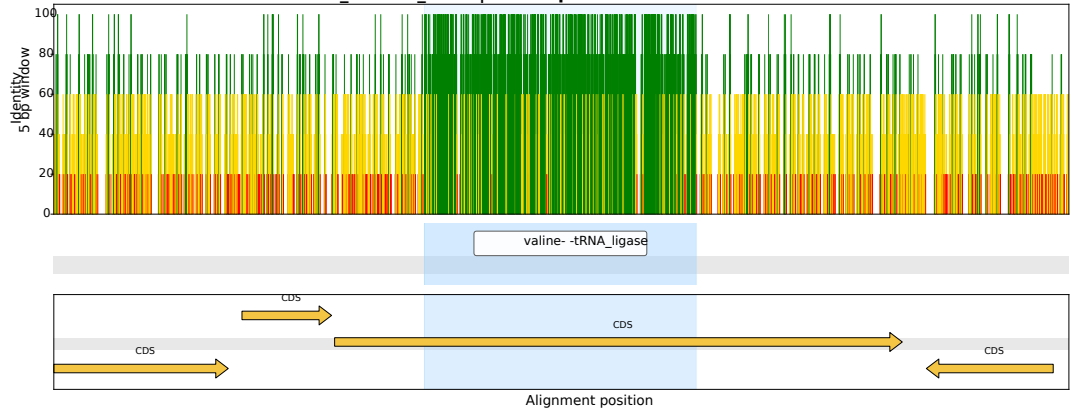

CI\_Harlain\_0143 | Arsenophonus OZ026540.1

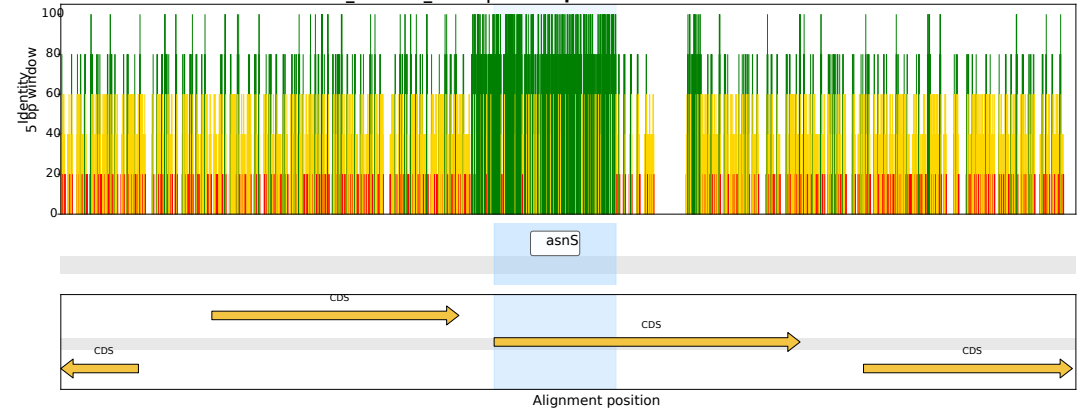

CI\_Harlain\_0279 | Arsenophonus CP123499.1

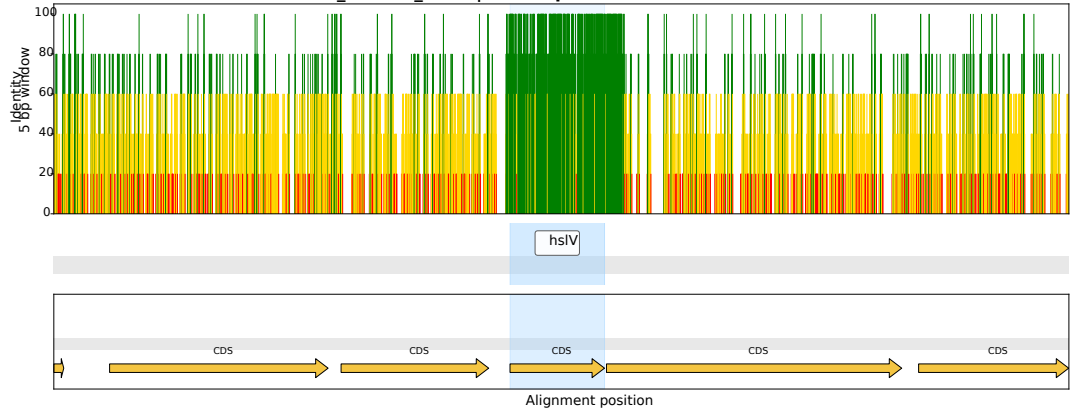

CI\_Harlain\_0026 | Arsenophonus CP123498.1

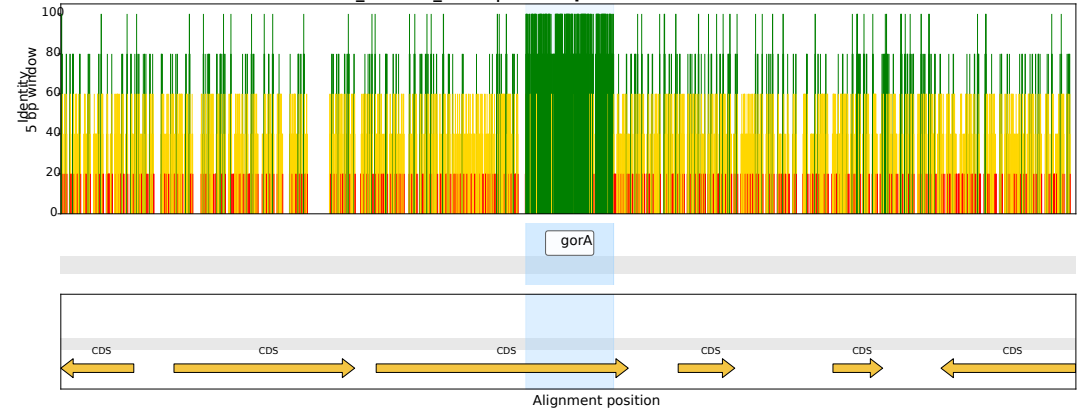

CI\_Harlain\_0133 | Arsenophonus OZ026540.1

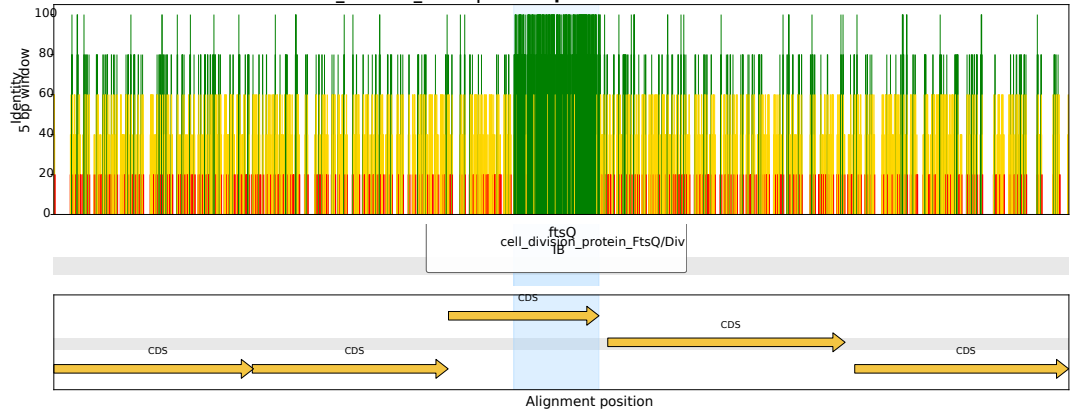

CI\_Harlain\_0264 | Arsenophonus CP123504.1

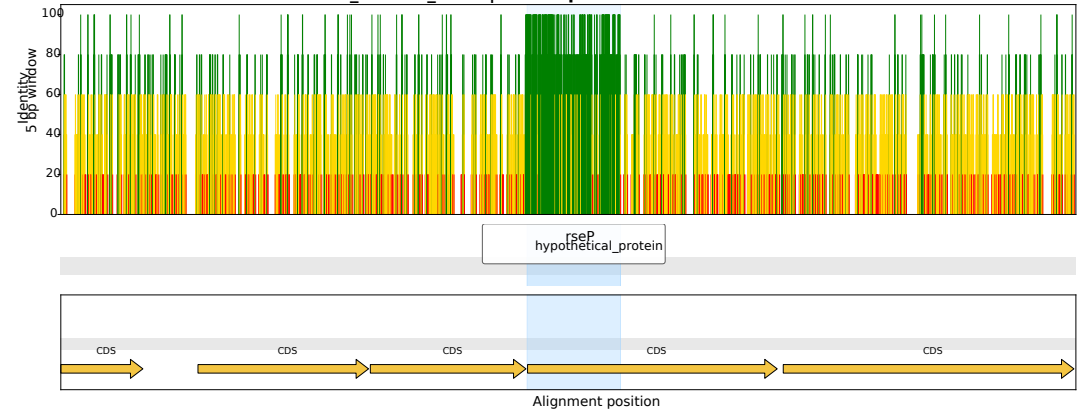

CI\_Harlain\_0358 | *Arsenophonus* OZ026540.1

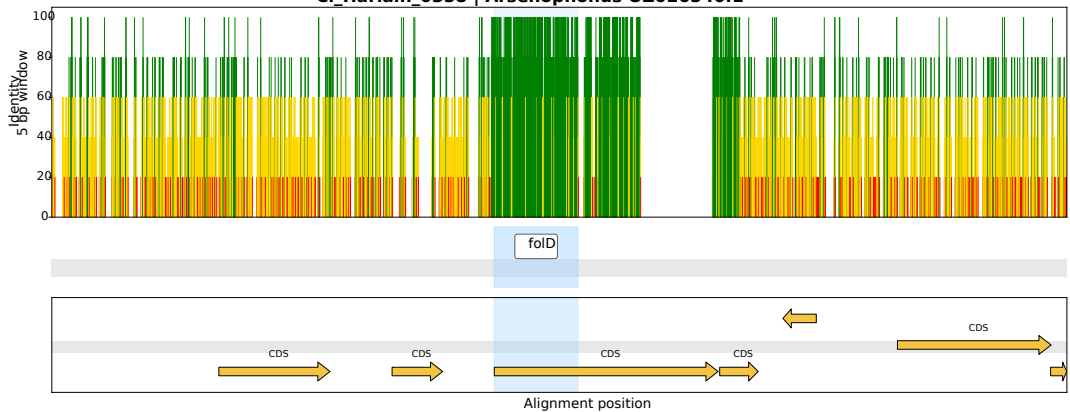

CI\_Harlain\_0017 | *Arsenophonus* CP123759.1

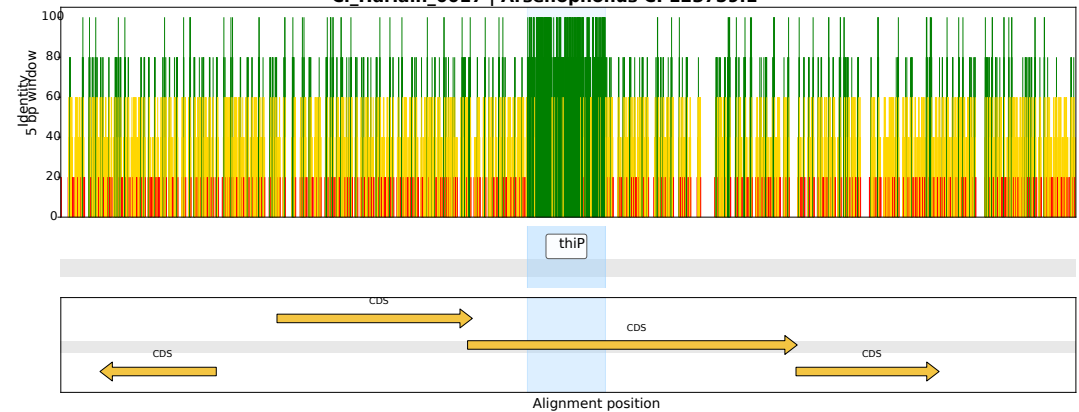

CI\_Harlain\_0276 | *Arsenophonus* CP038613.1

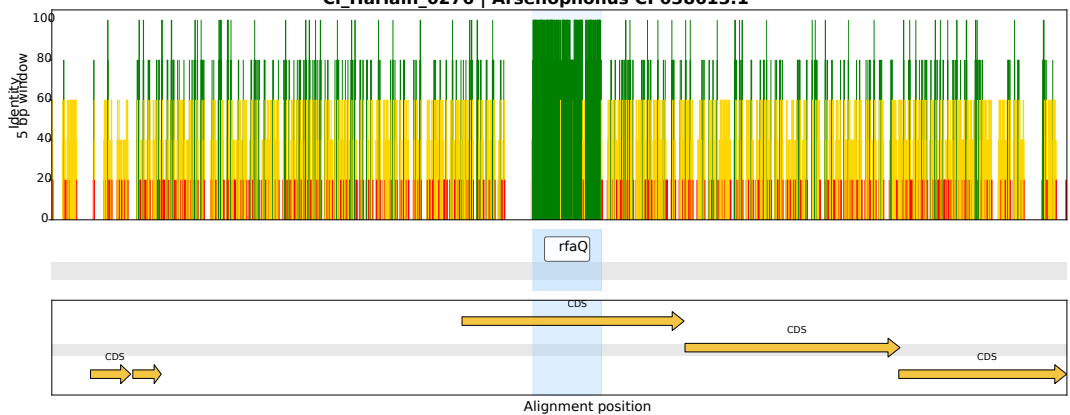

CI\_Harlain\_0072 | *Arsenophonus* CP123499.1

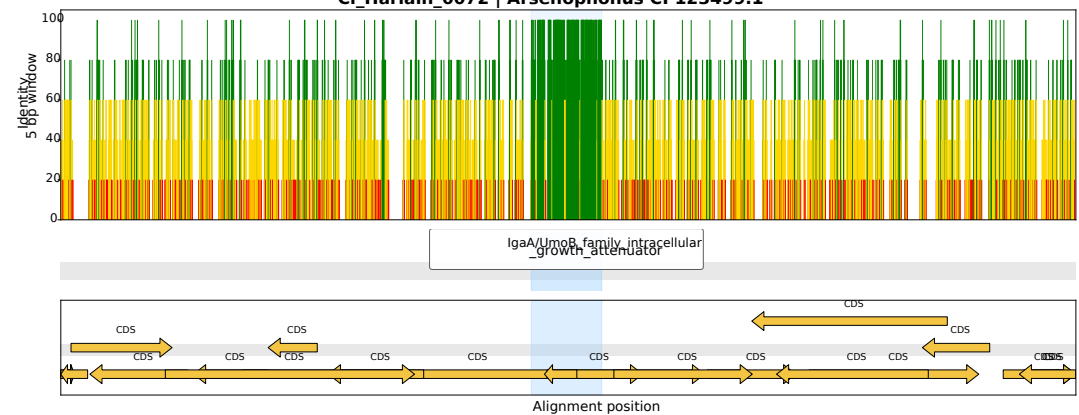

CI\_Harlain\_0342 | *Arsenophonus* CP038155.1

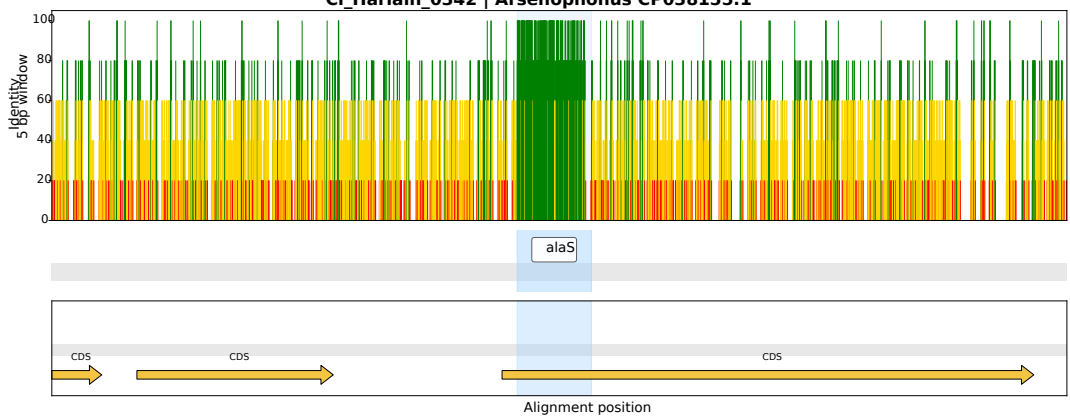

CI\_Harlain\_0375 | *Arsenophonus* CP123759.1

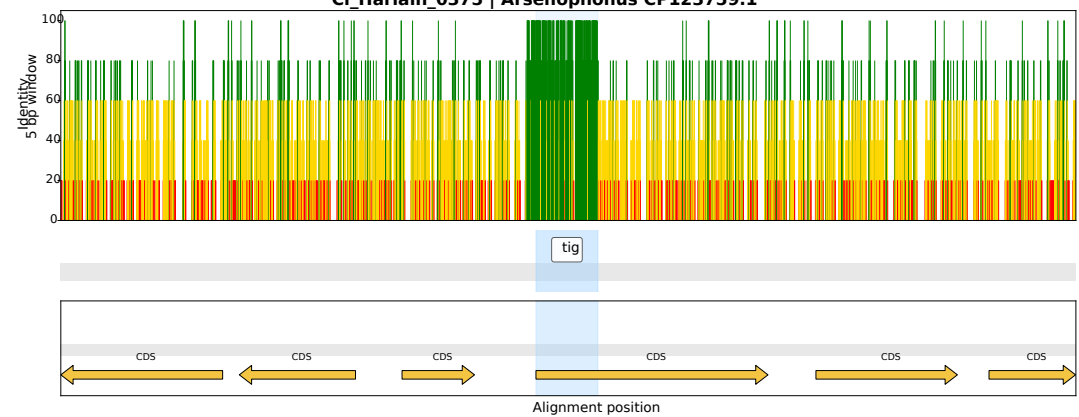

CI\_Harlain\_0066 | *Arsenophonus* CP084222.1

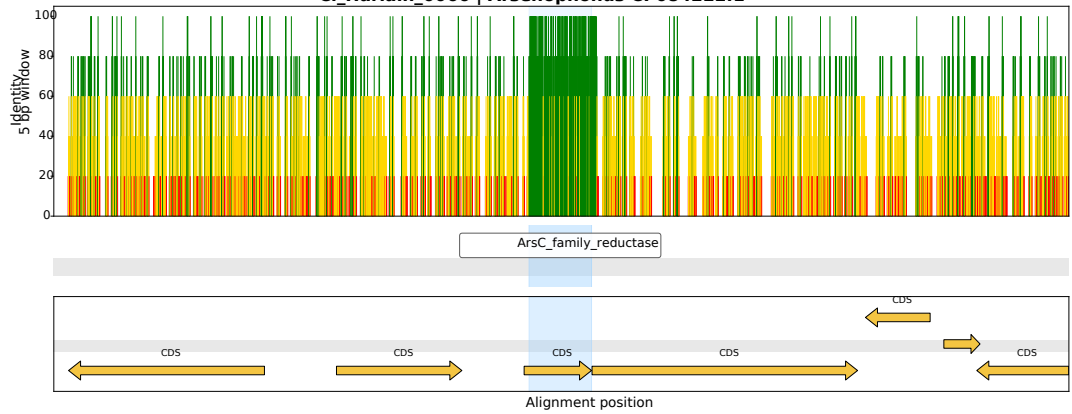

CI\_Harlain\_0144 | *Arsenophonus* CP038613.1

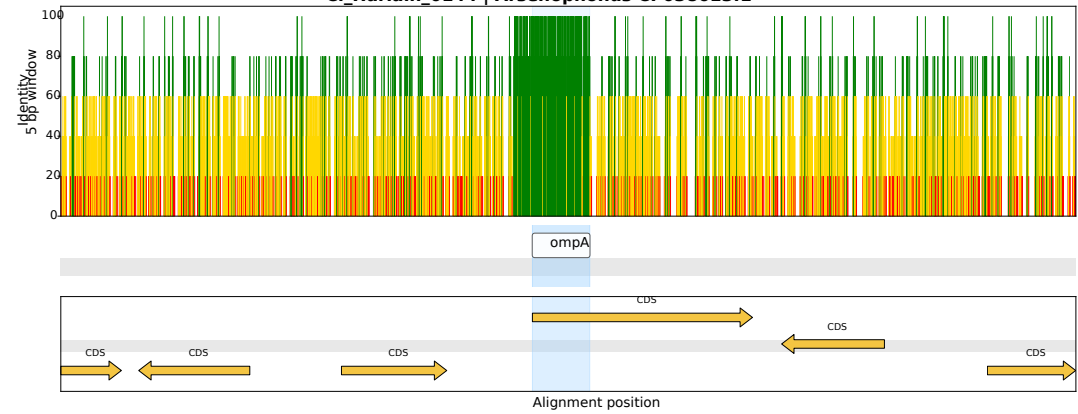

CI\_Harlain\_0287 | *Arsenophonus* CP038613.1

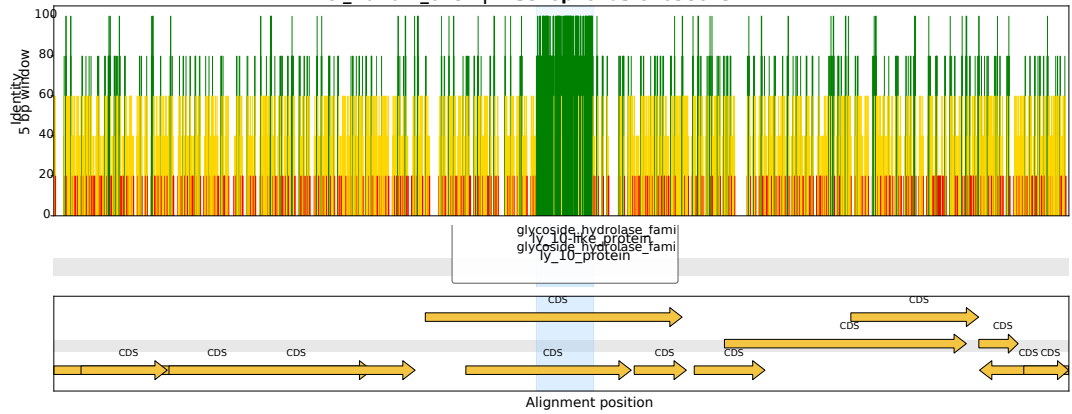

CI\_Harlain\_0394 | *Arsenophonus* CP084222.1

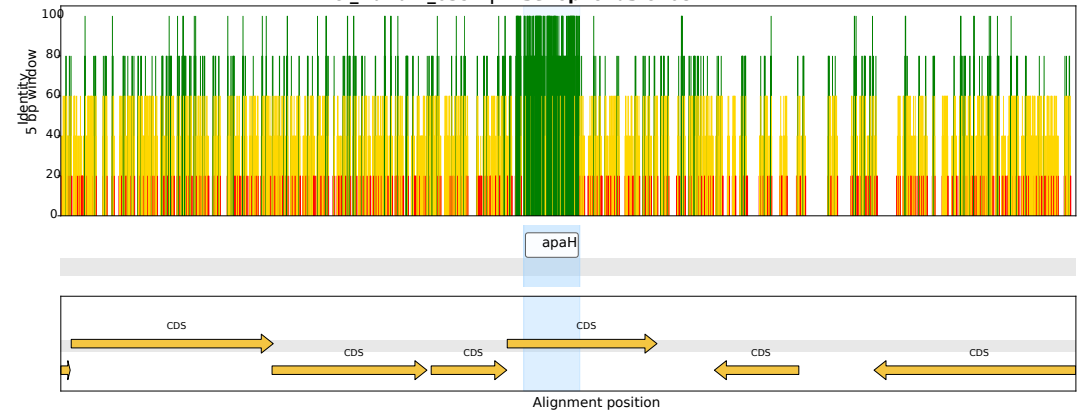

CI\_Harlain\_0451 | *Arsenophonus* LR025108.1

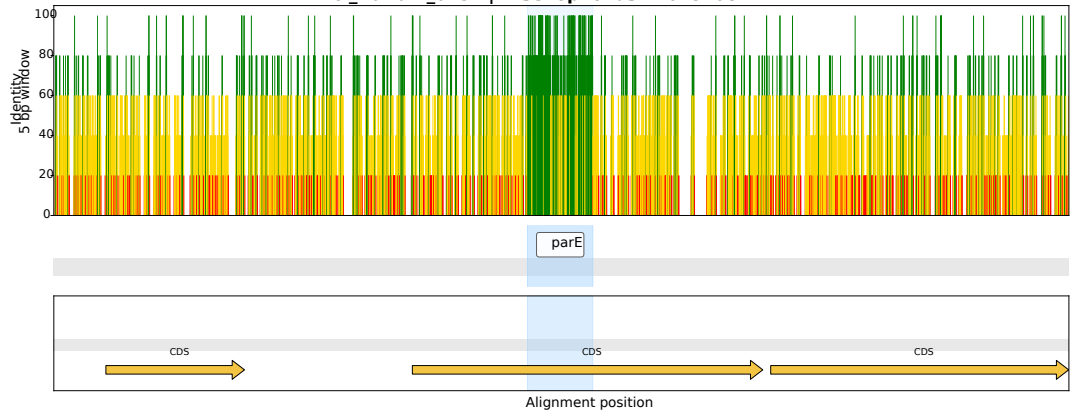

CI\_Harlain\_0328 | *Arsenophonus* CP084222.1

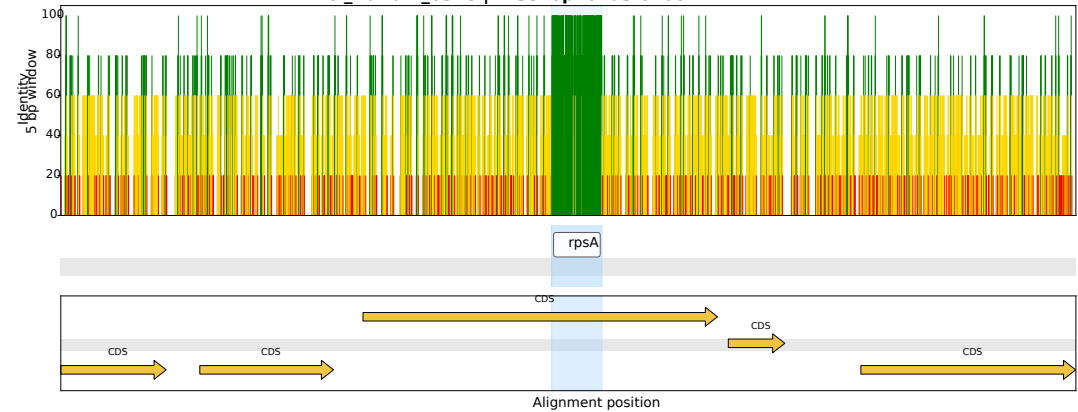

CI\_Harlain\_0295 | *Arsenophonus* CP123759.1

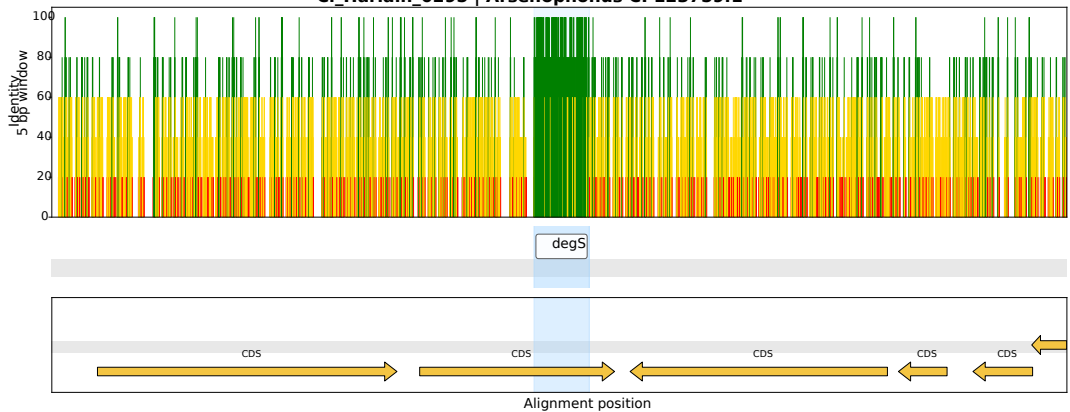

CI\_Harlain\_0355 | *Arsenophonus* CP123759.1

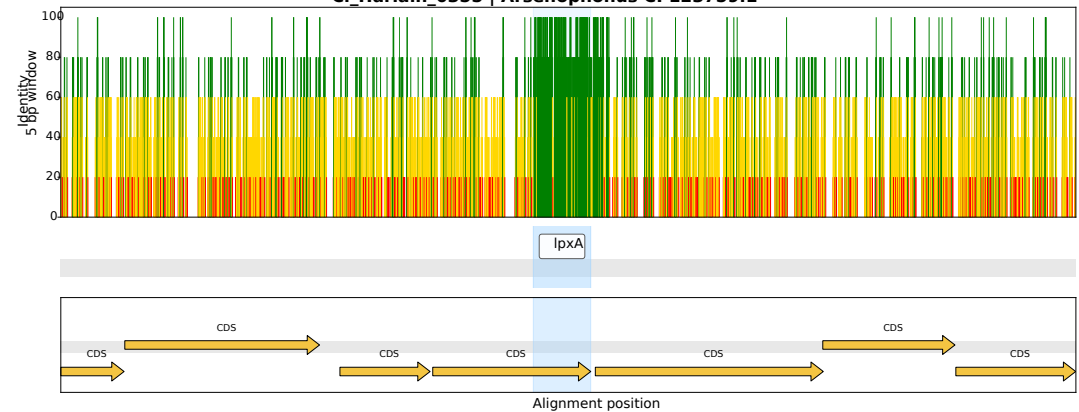

CI\_Harlain\_0224 | *Arsenophonus* OZ026540.1

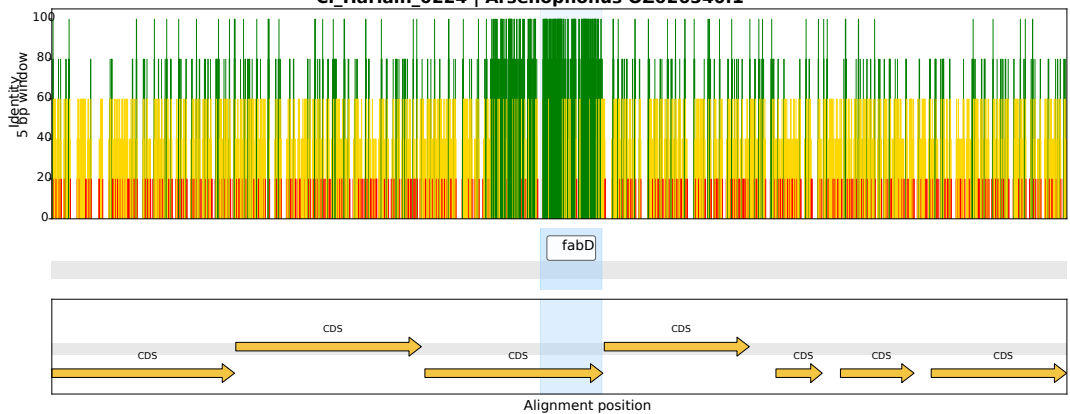

CI\_Harlain\_0439 | *Arsenophonus* CP123499.1

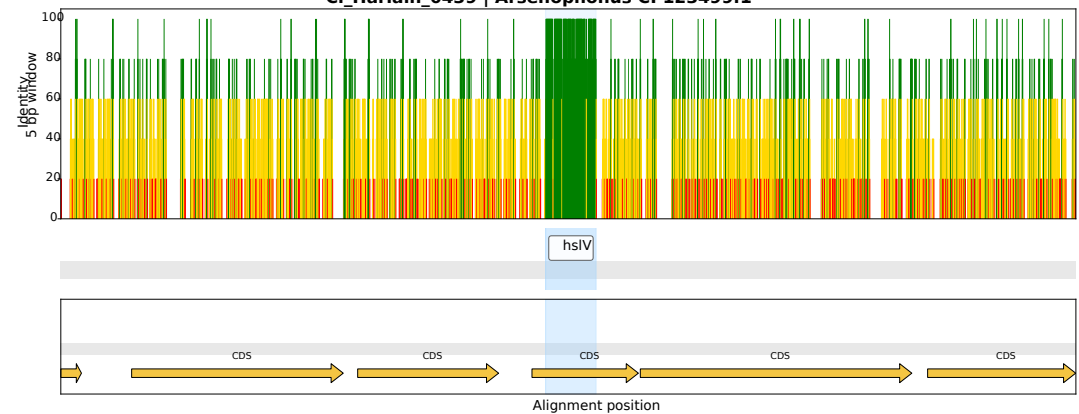

CI\_Harlain\_0380 | *Arsenophonus* OZ026540.1

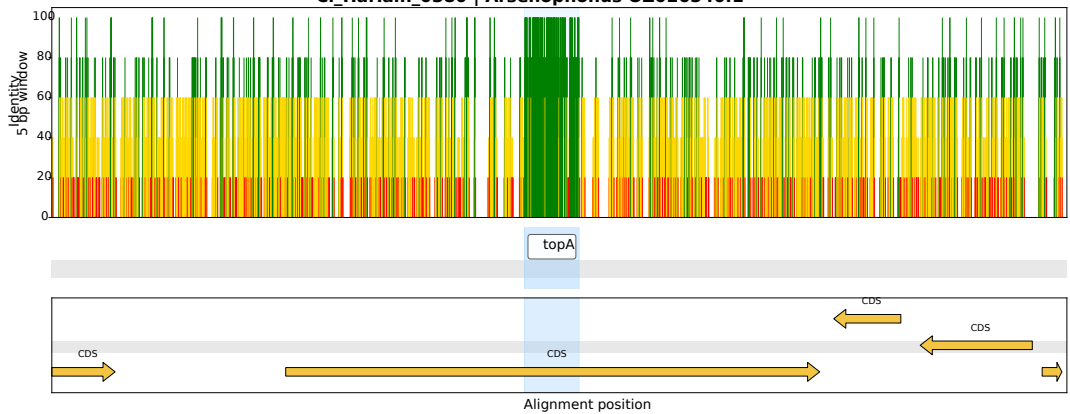

CI\_Harlain\_0362 | *Arsenophonus* CP038613.1

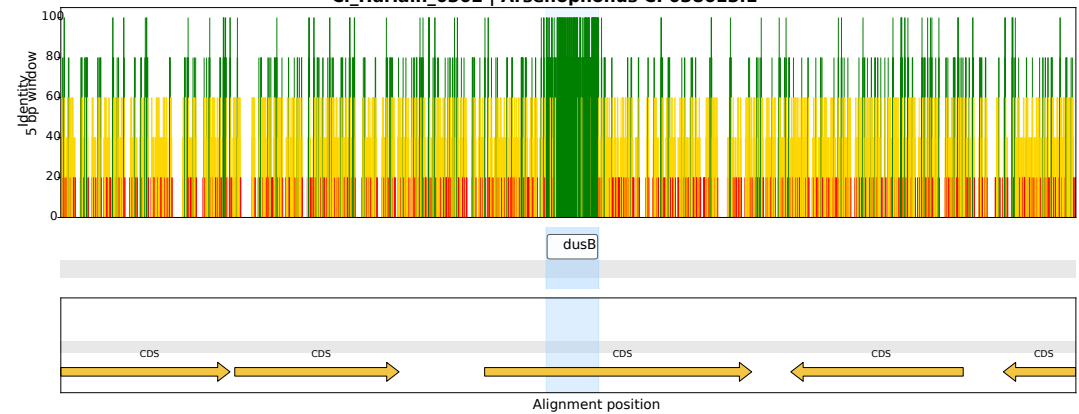

CI\_Harlain\_0102 | *Arsenophonus* OZ026540.1

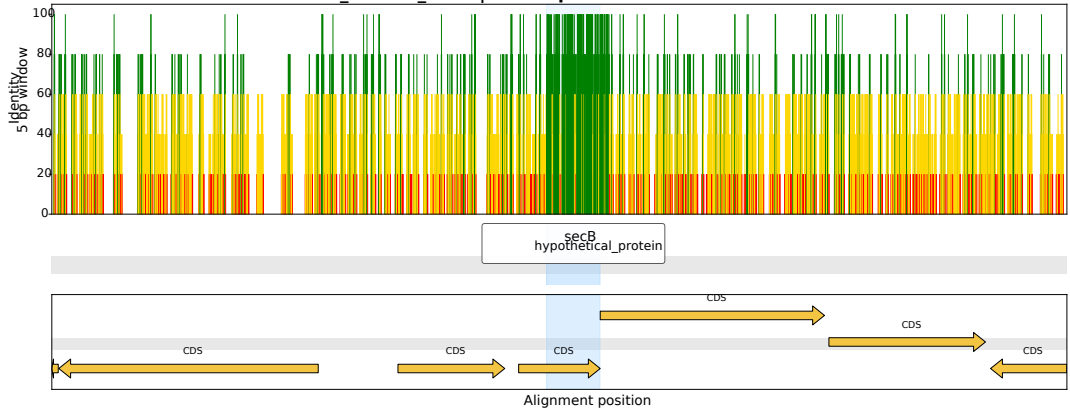

CI\_Harlain\_0148 | *Arsenophonus* CP123498.1

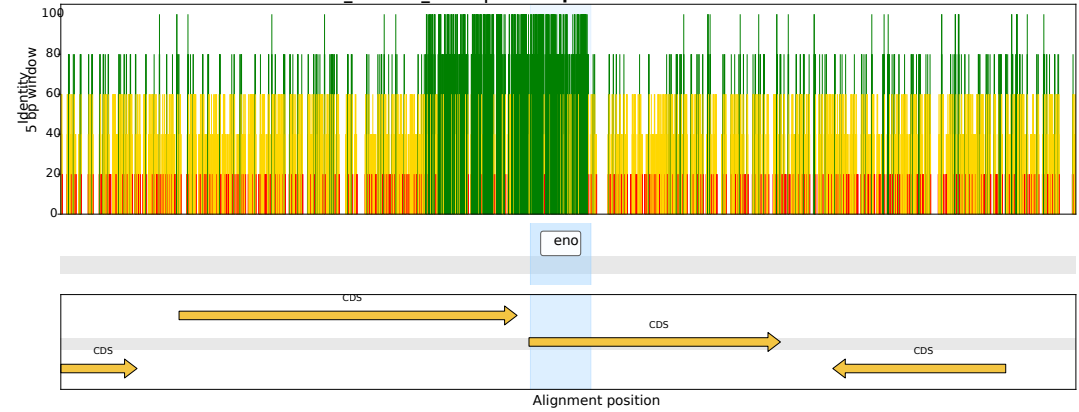

CI\_Harlain\_0424 | *Arsenophonus* OZ026540.1

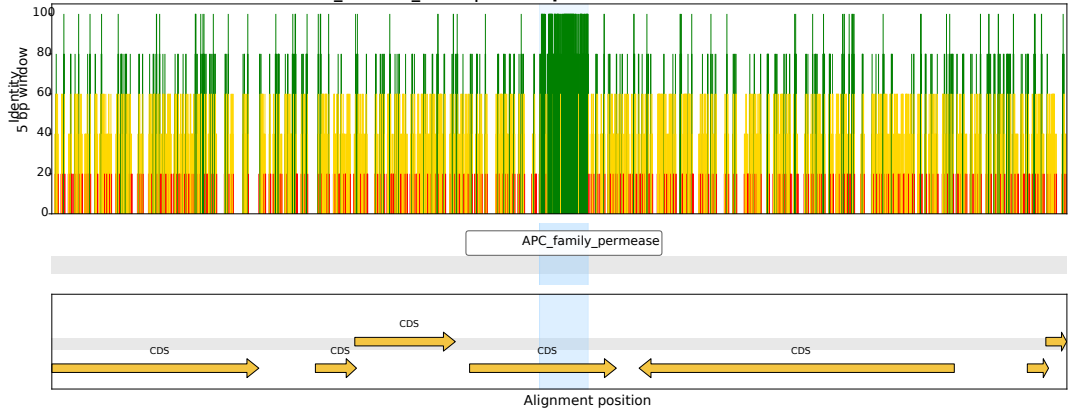

CI\_Harlain\_0071 | *Arsenophonus* CP038613.1

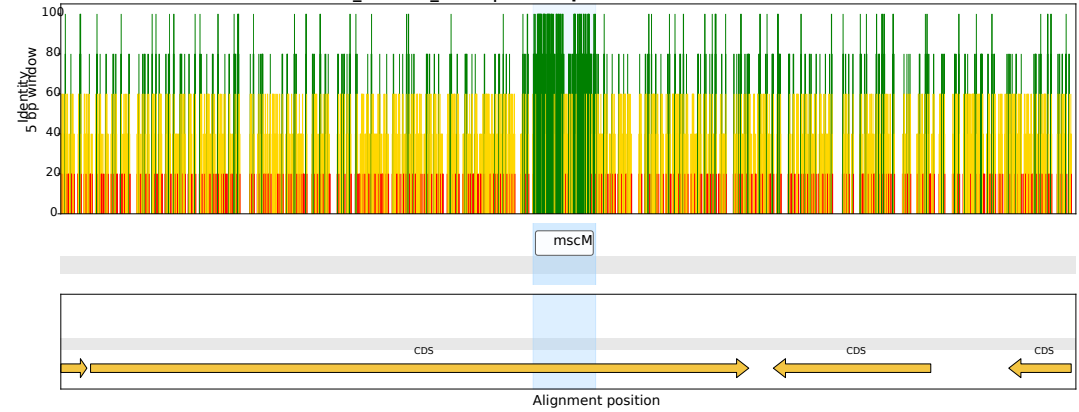

CI\_Harlain\_0233 | *Arsenophonus* CP123759.1

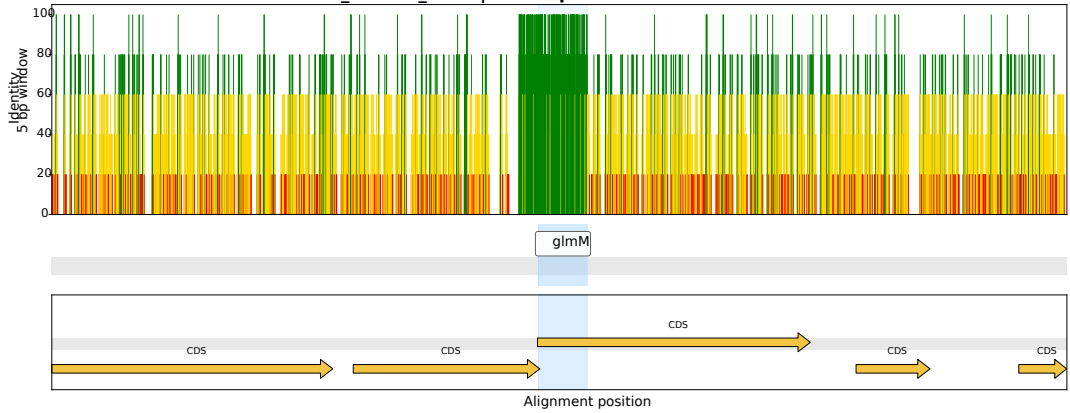

CI\_Harlain\_0408 | *Arsenophonus* OZ026540.1

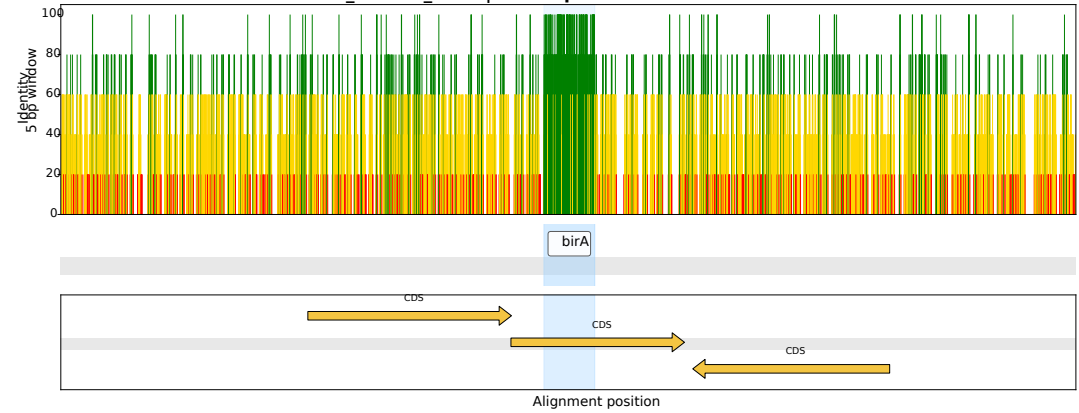

CI\_Harlain\_0208 | *Arsenophonus* CP084222.1

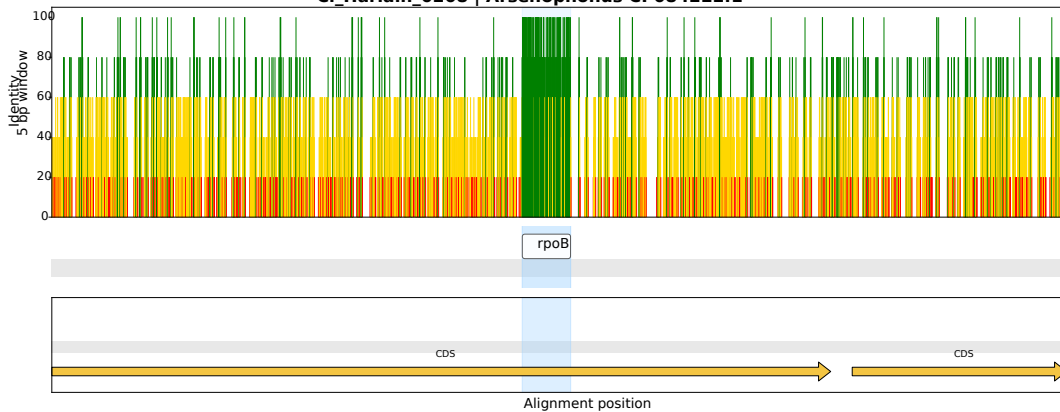

CI\_Harlain\_0169 | *Arsenophonus* CP123499.1

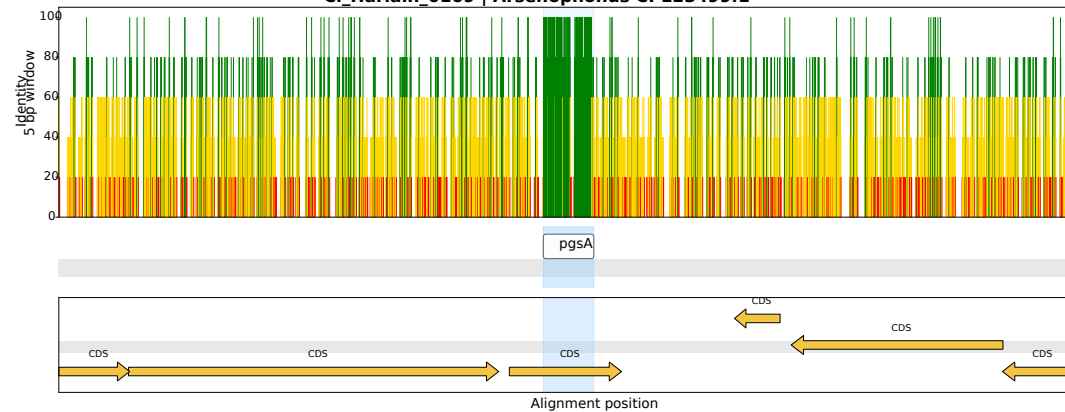

CI\_Harlain\_0469 | *Arsenophonus* CP038613.1

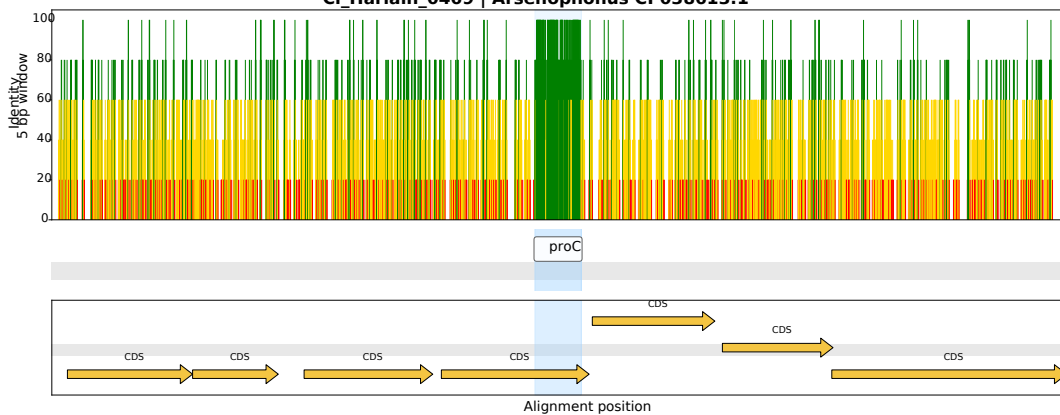

CI\_Harlain\_0442 | *Arsenophonus* CP123759.1

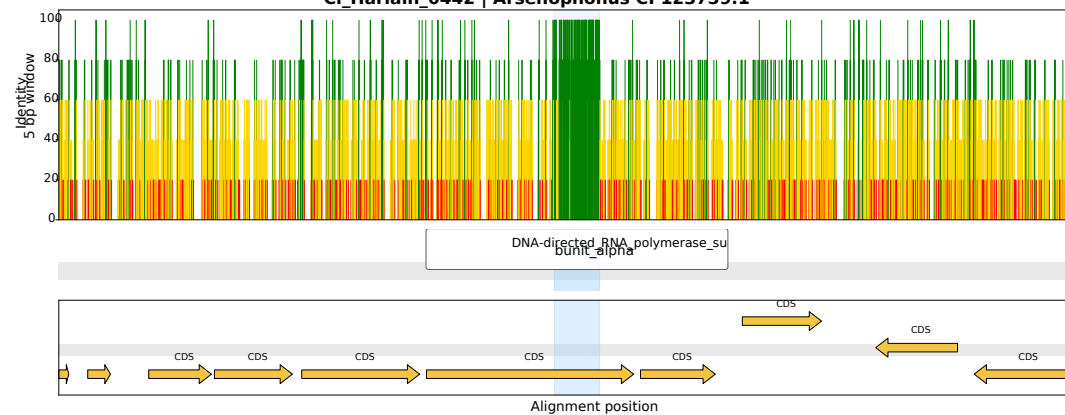

CI\_Harlain\_0245 | *Arsenophonus* CP123759.1

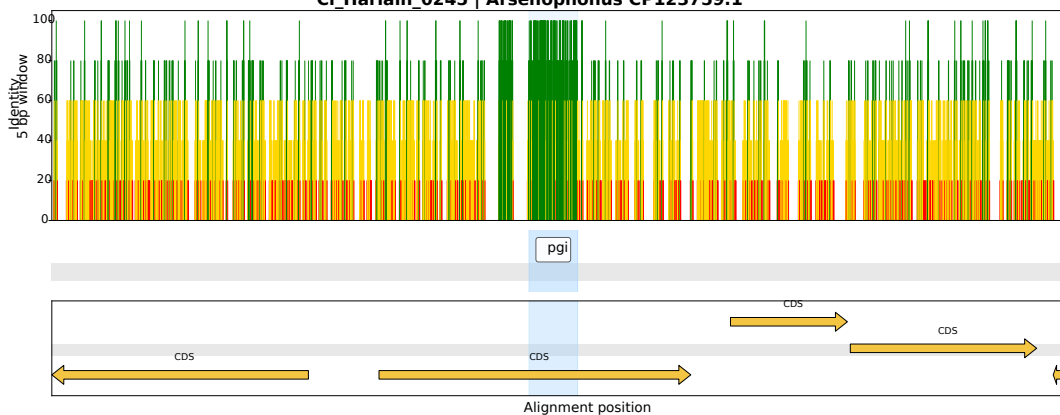

CI\_Harlain\_0432 | *Arsenophonus* CP123499.1

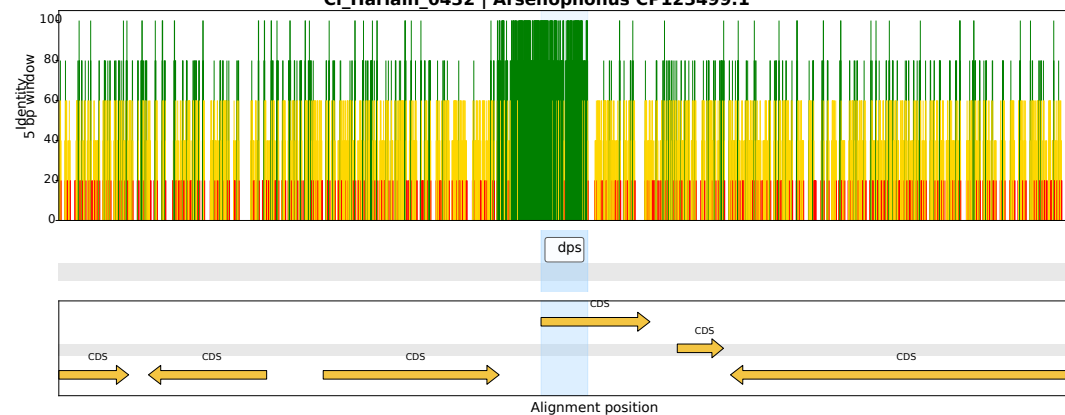

CI\_Harlain\_0242 | *Arsenophonus* CP038613.1

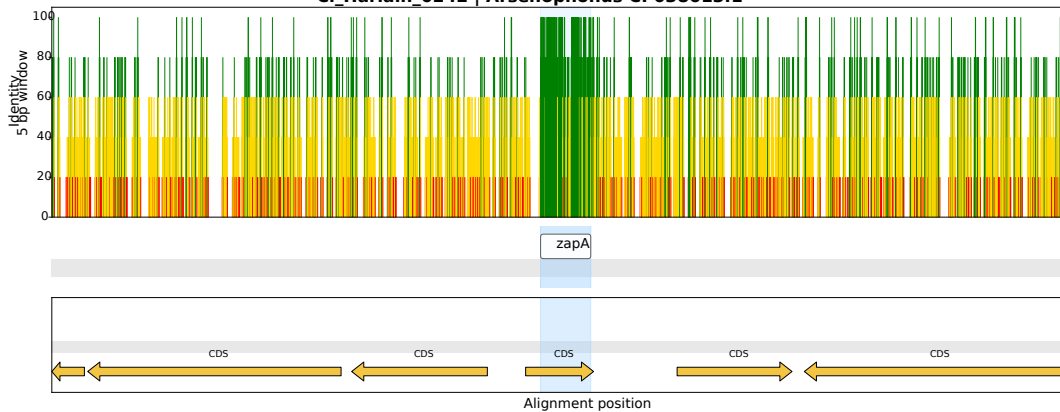

CI\_Harlain\_0046 | *Arsenophonus* CP084222.1

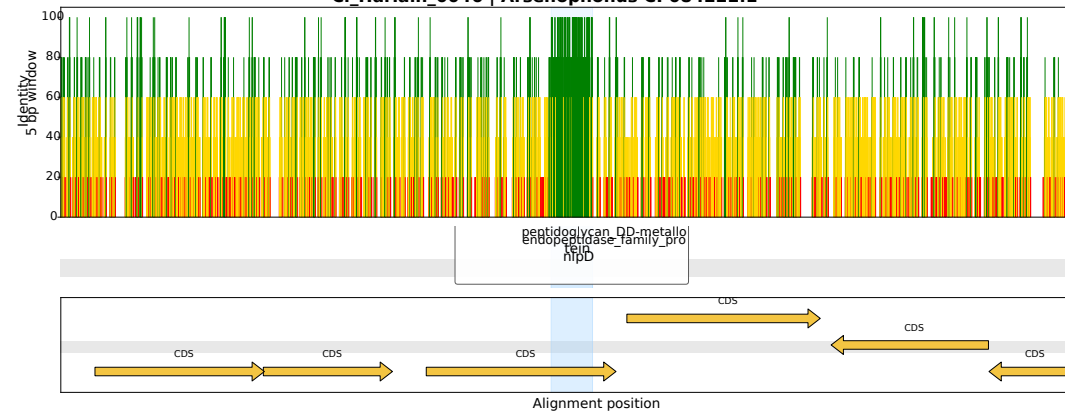

CI\_Harlain\_0161 | *Arsenophonus* CP123499.1

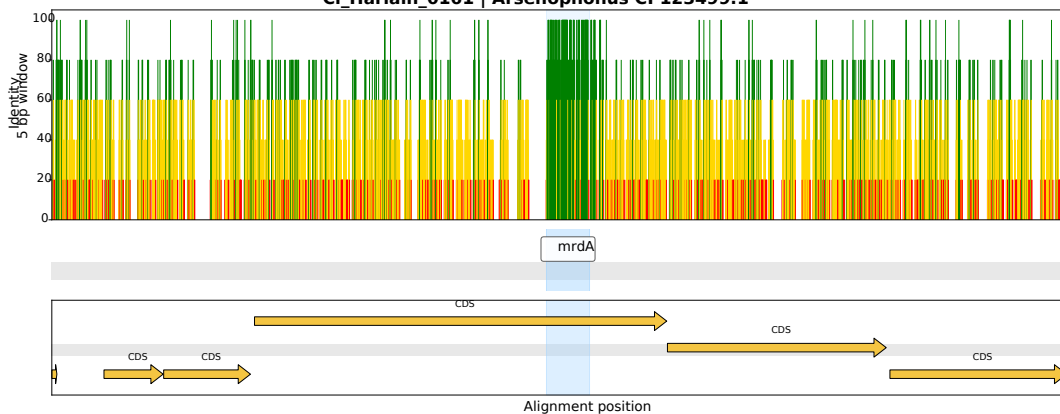

CI\_Harlain\_0307 | *Arsenophonus* CP123499.1

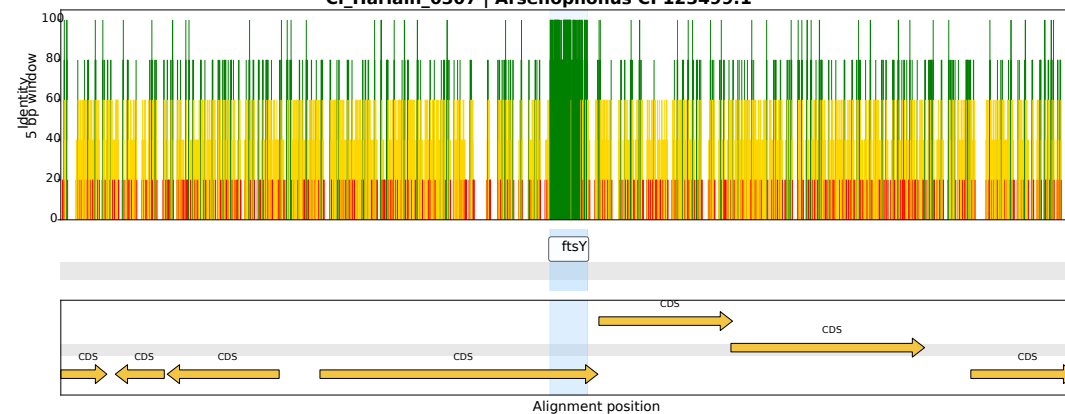

CI\_Harlain\_0080 | *Arsenophonus* OZ026540.1

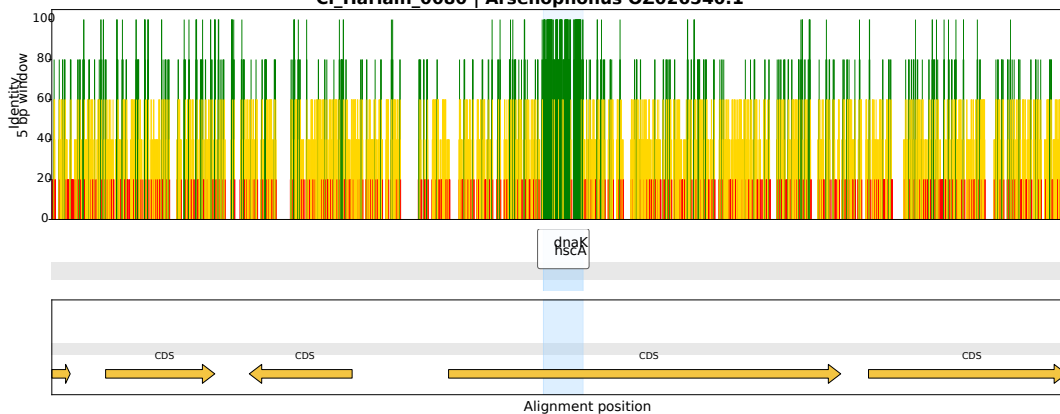

CI\_Harlain\_0095 | *Arsenophonus* CP038613.1

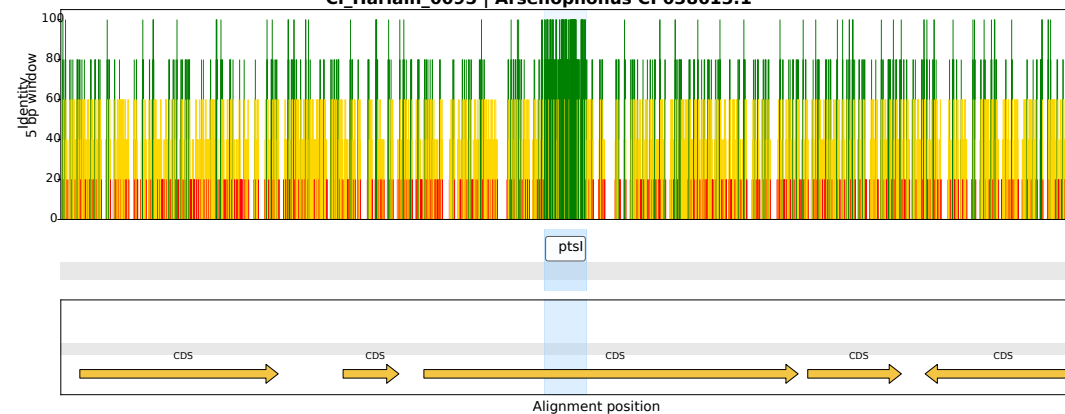

CI\_Harlain\_0462 | *Arsenophonus* CP123499.1

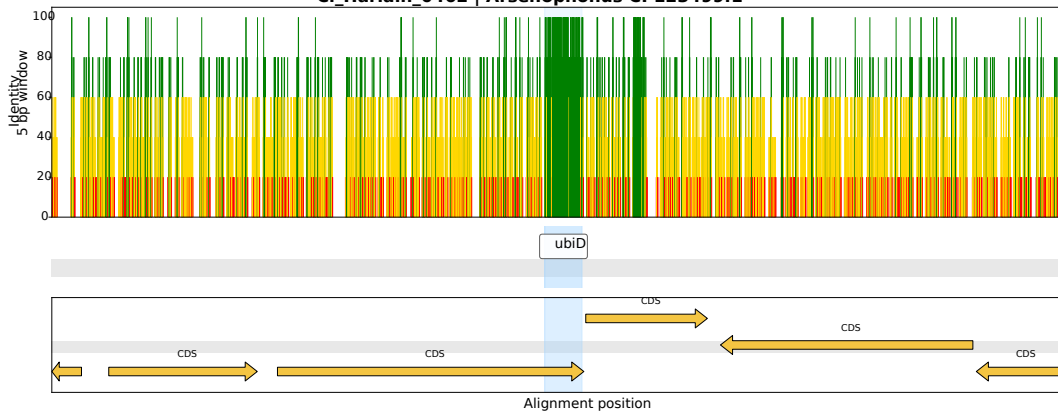

CI\_Harlain\_0269 | *Arsenophonus* OZ026540.1

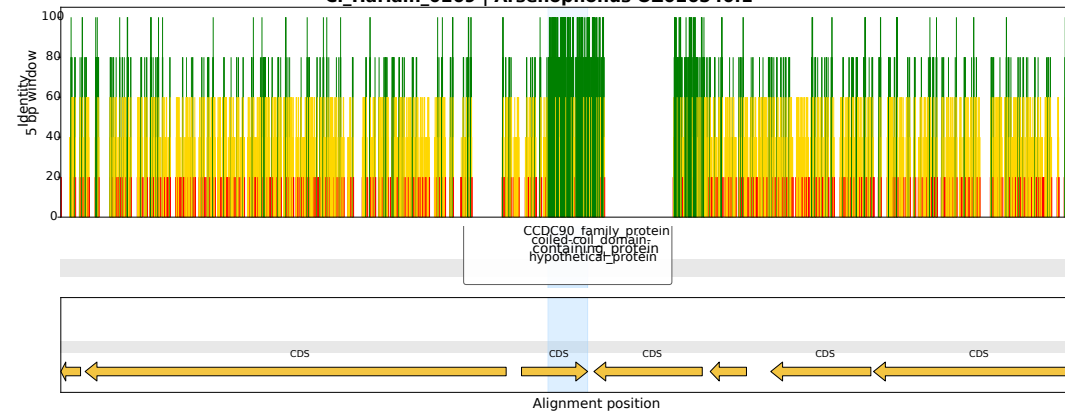

CI\_Harlain\_0310 | *Arsenophonus* CP123759.1

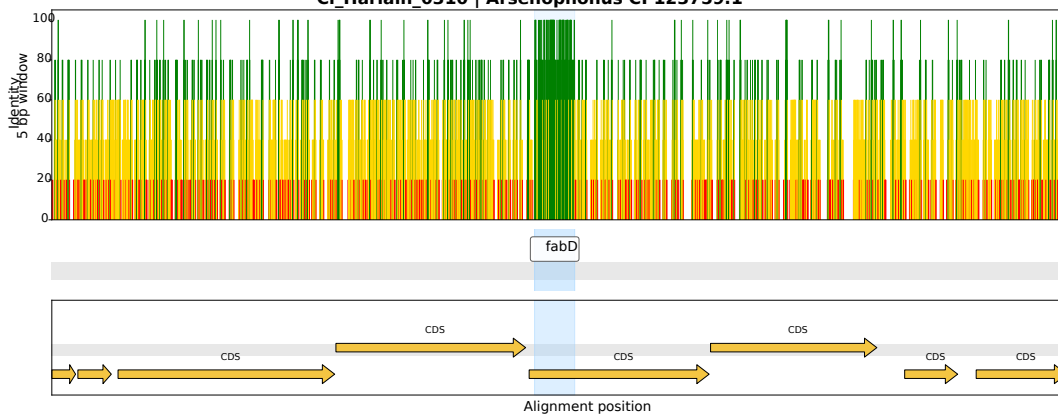

CI\_Harlain\_0191 | *Arsenophonus* CP123499.1

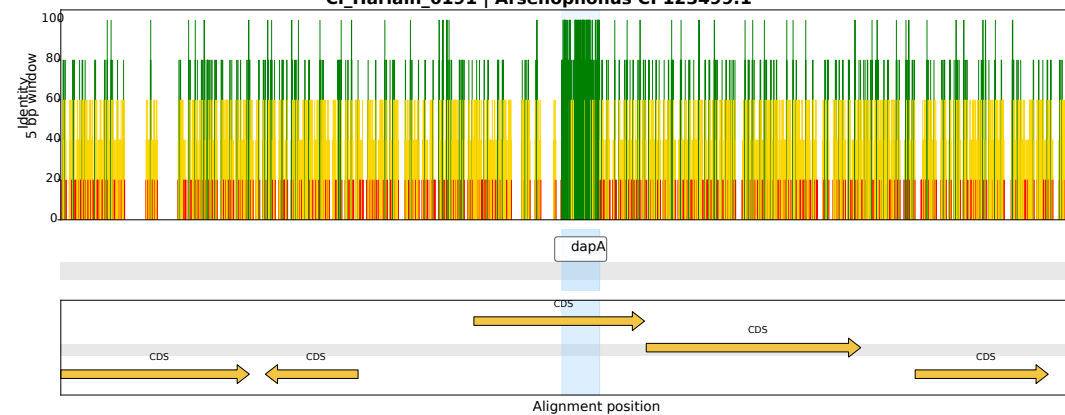

CI\_Harlain\_0167 | *Arsenophonus* CP123759.1

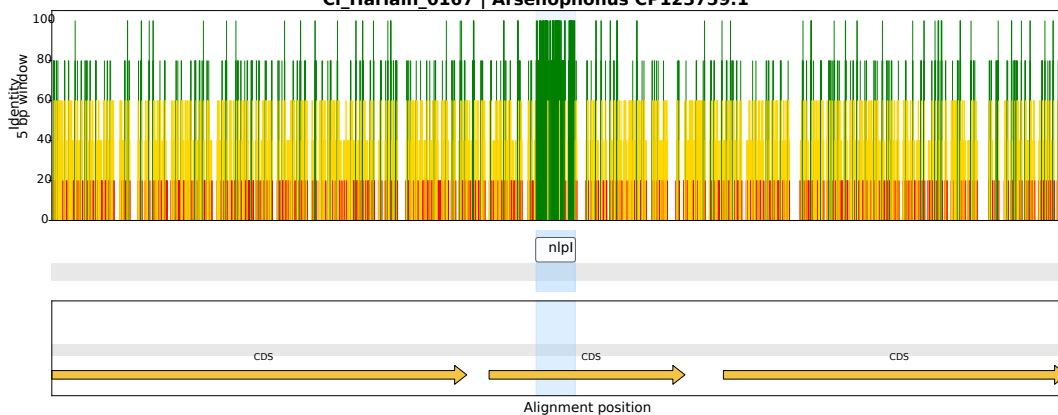

CI\_Harlain\_0041 | *Arsenophonus* CP038613.1

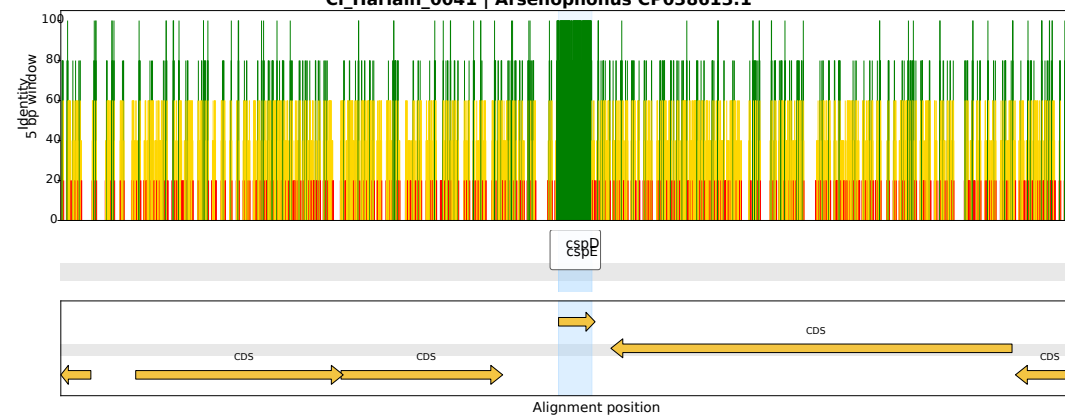

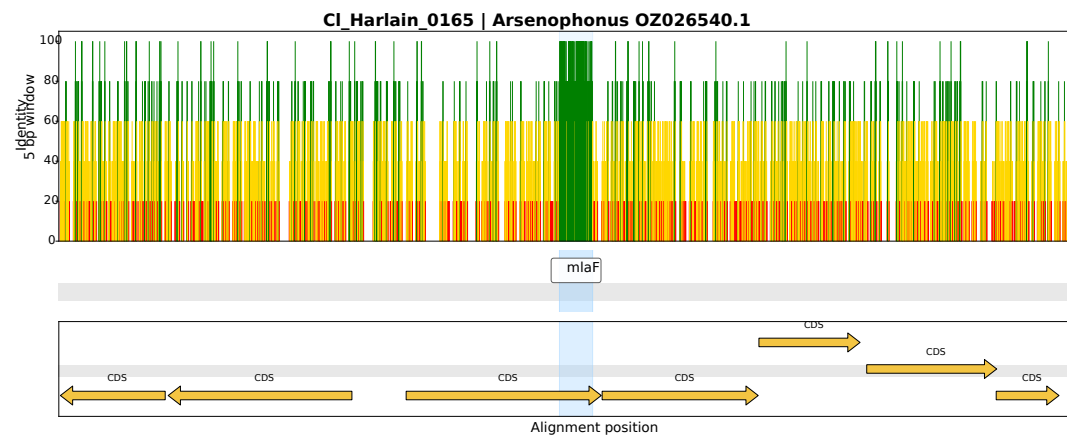

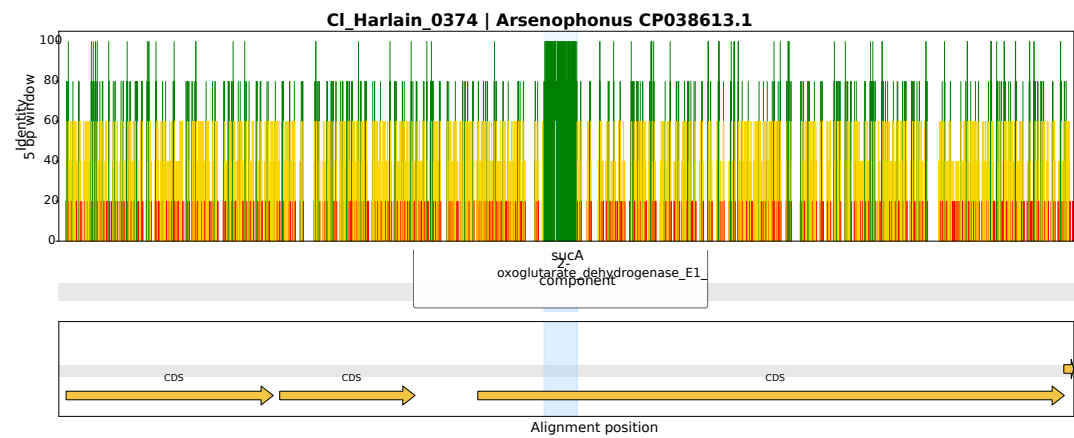

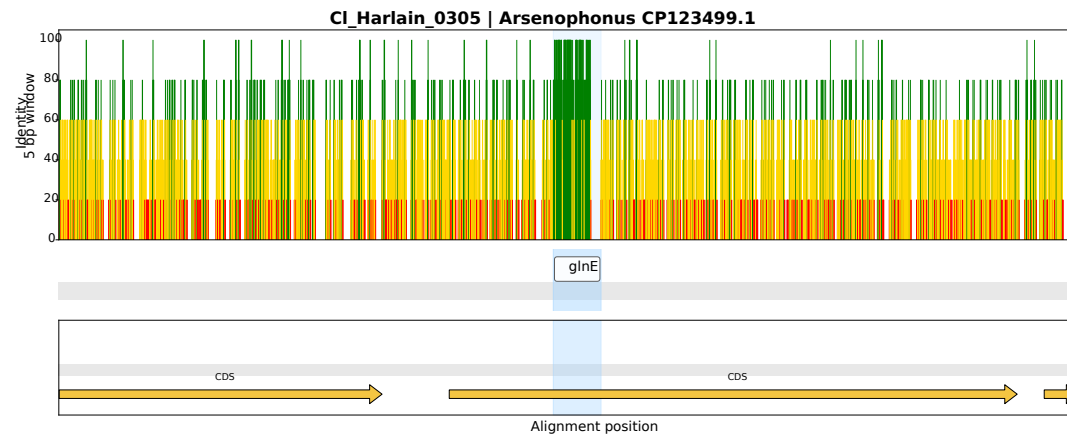

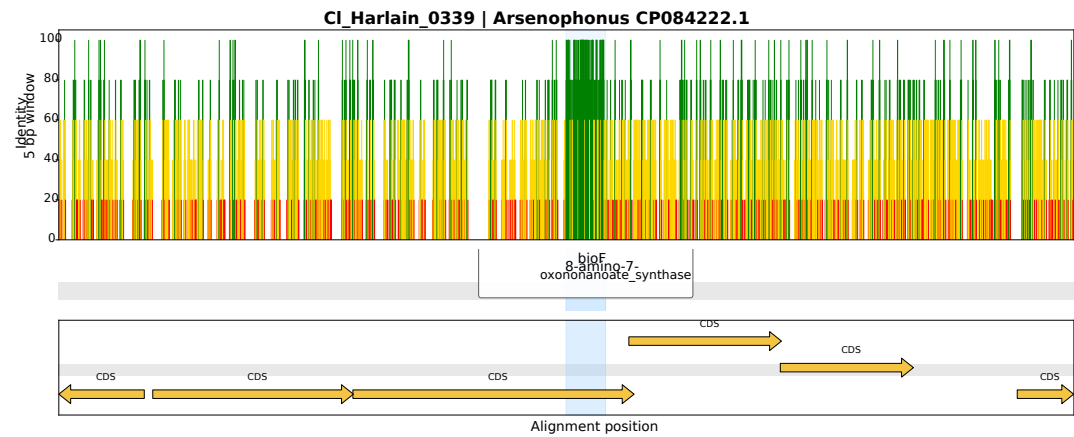

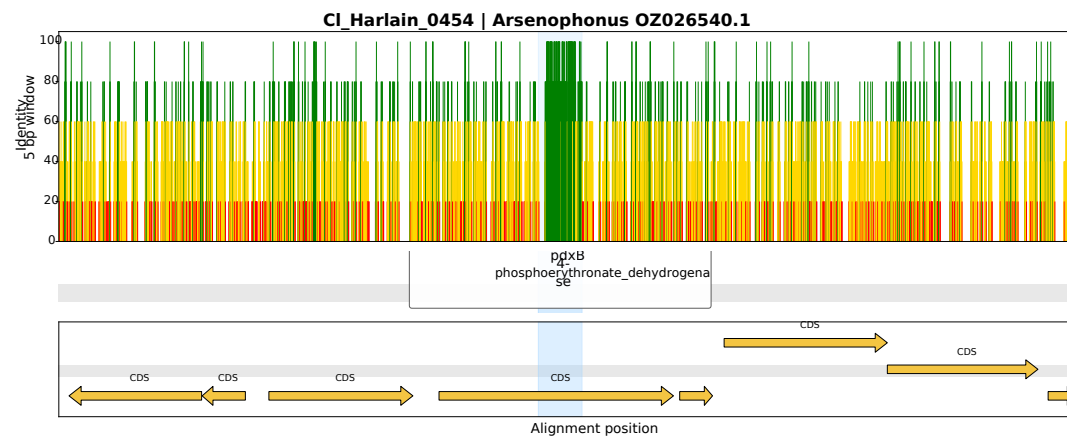

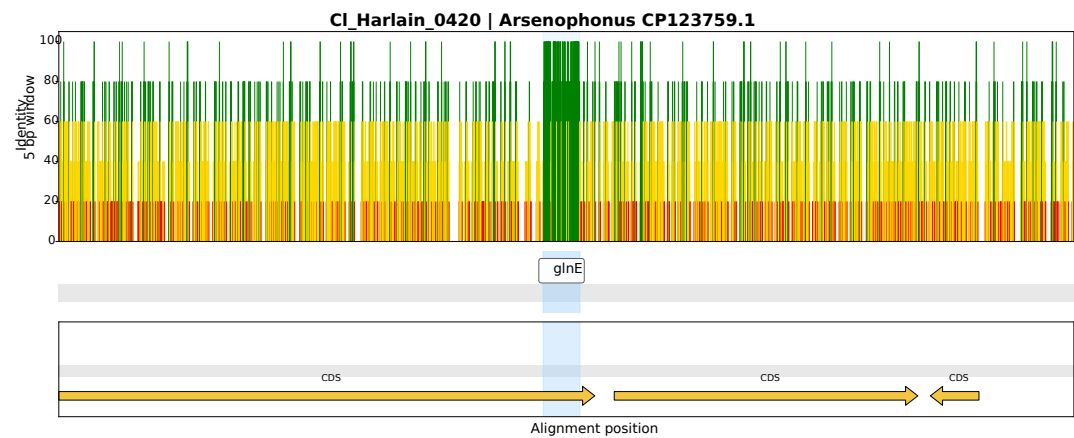

CI\_Harlain\_0416 | *Arsenophonus* CP038613.1

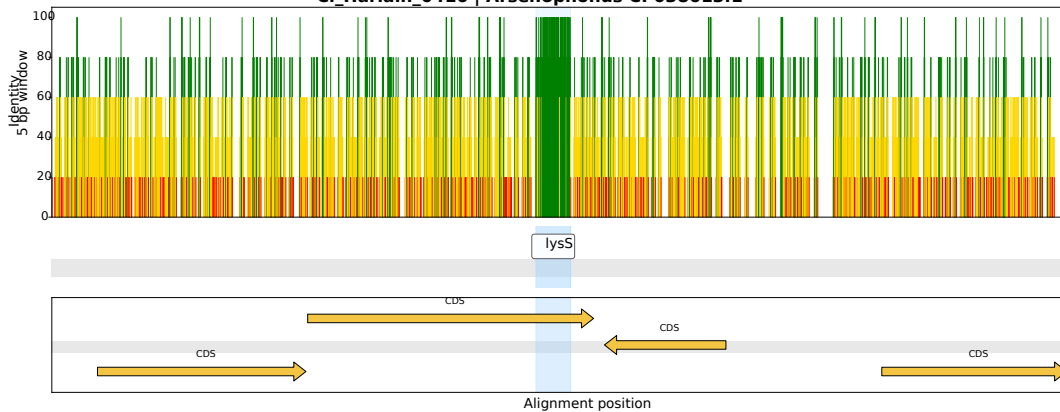

CI\_Harlain\_0016 | *Arsenophonus* OZ026540.1

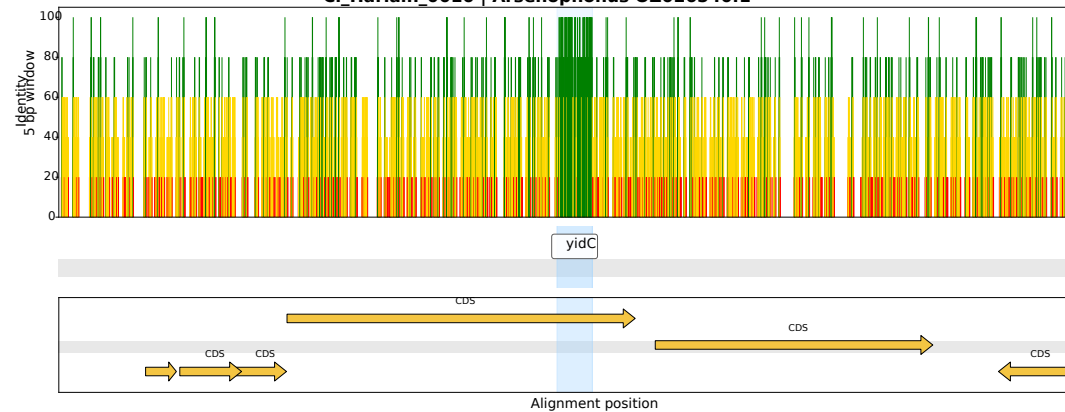

CI\_Harlain\_0128 | *Arsenophonus* CP038155.1

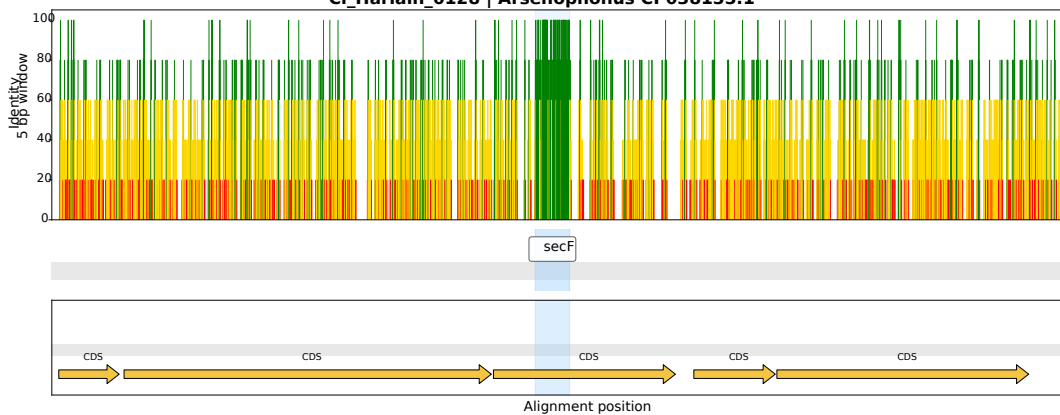

CI\_Harlain\_0155 | *Arsenophonus* OZ026540.1

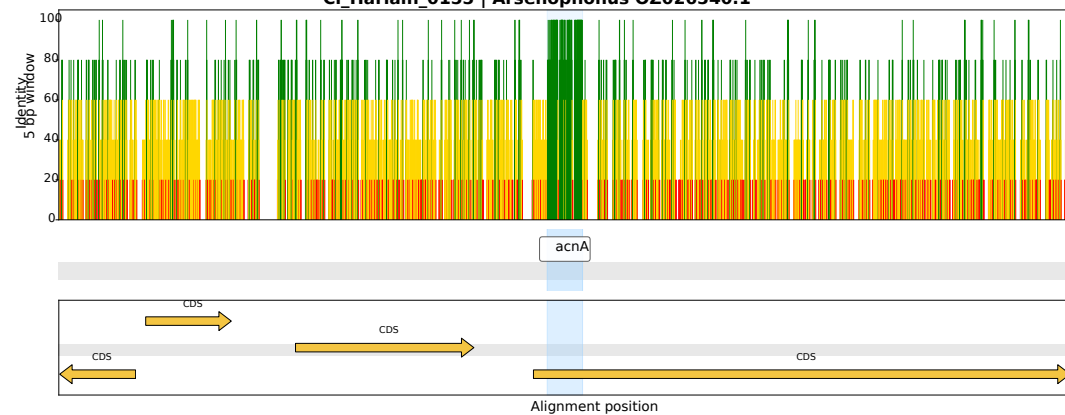

CI\_Harlain\_0404 | *Arsenophonus* CP084222.1

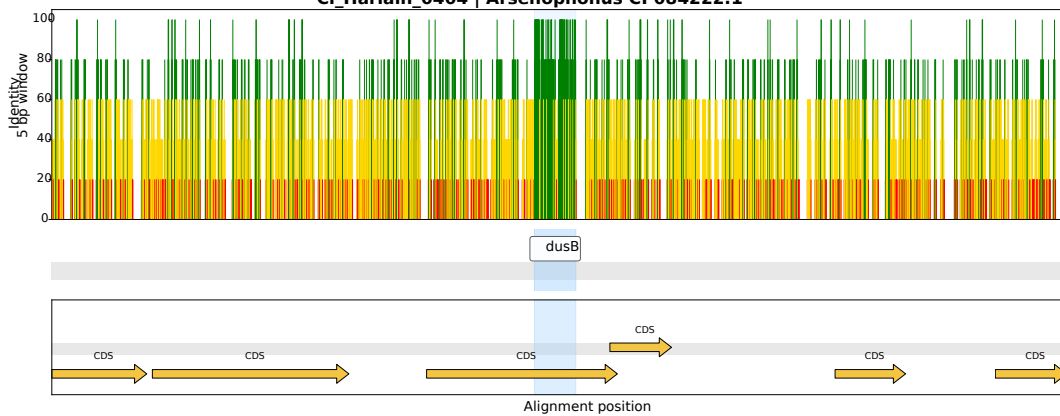

CI\_Harlain\_0347 | *Arsenophonus* CP123499.1

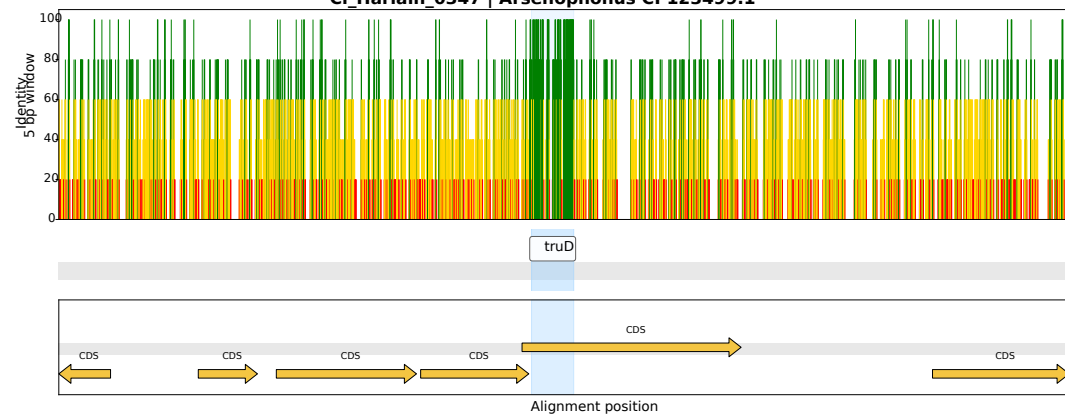

CI\_Harlain\_0337 | *Arsenophonus* OZ026540.1

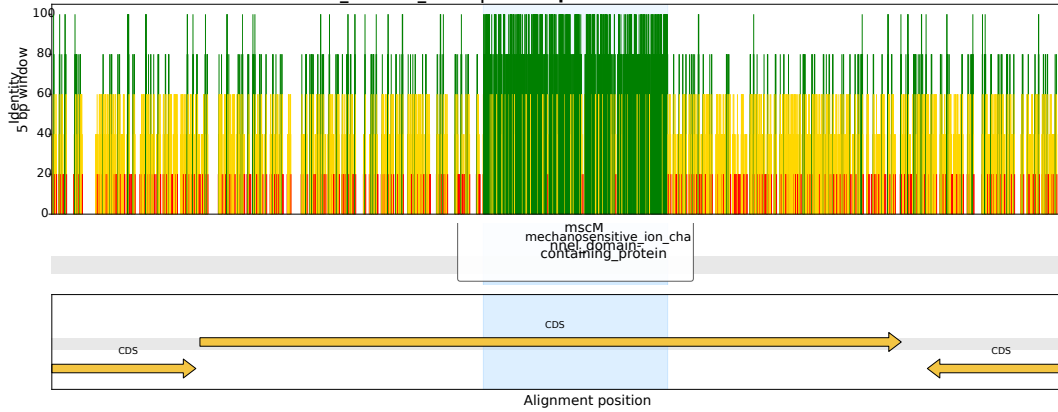

CI\_Harlain\_0073 | *Arsenophonus* OZ026540.1

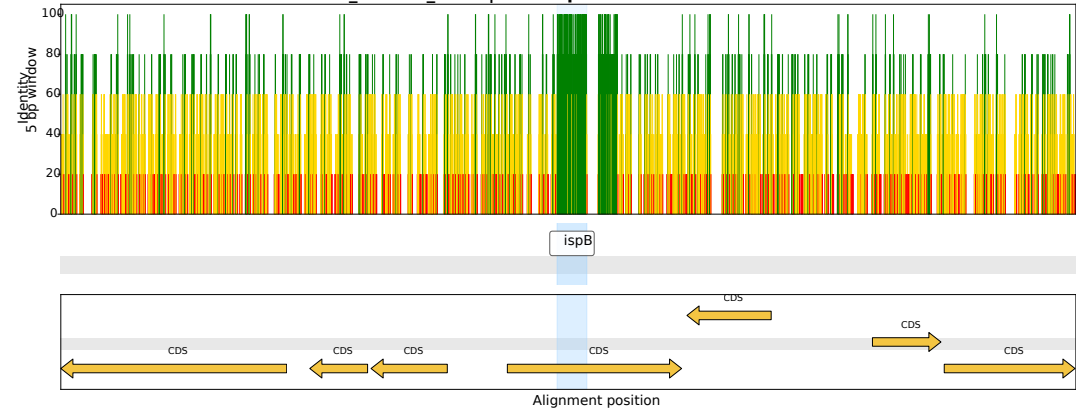

CI\_Harlain\_0419 | *Arsenophonus* CP038613.1

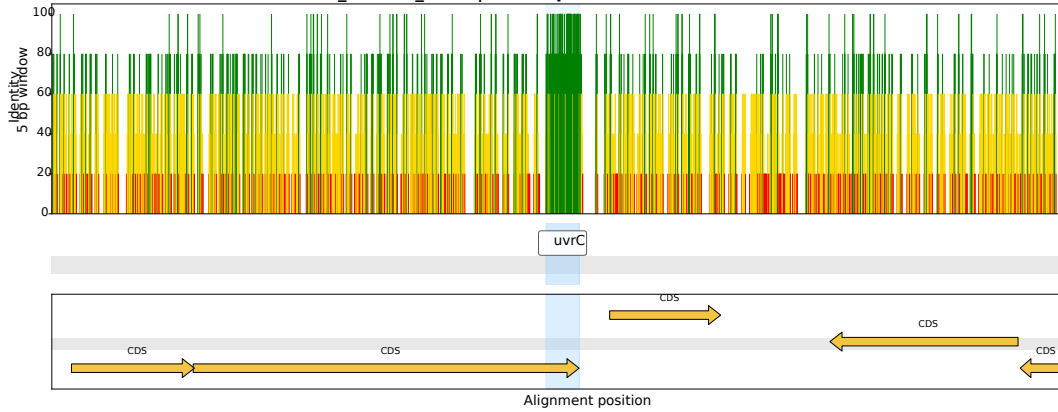

CI\_Harlain\_0108 | *Arsenophonus* CP038613.1

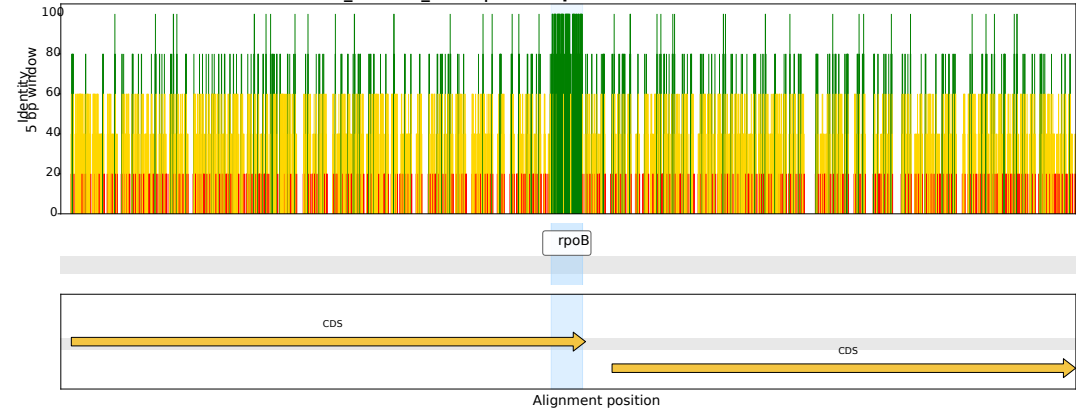

CI\_Harlain\_0247 | *Arsenophonus* CP123499.1

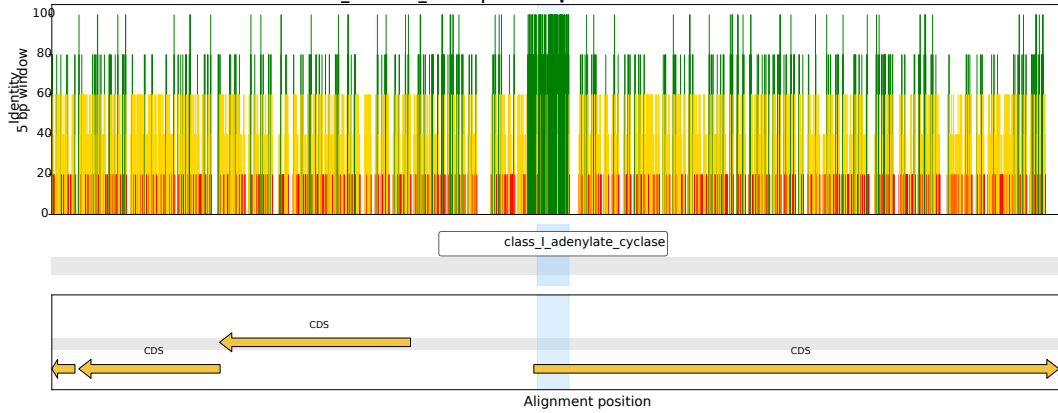

CI\_Harlain\_0239 | *Arsenophonus* CP038613.1

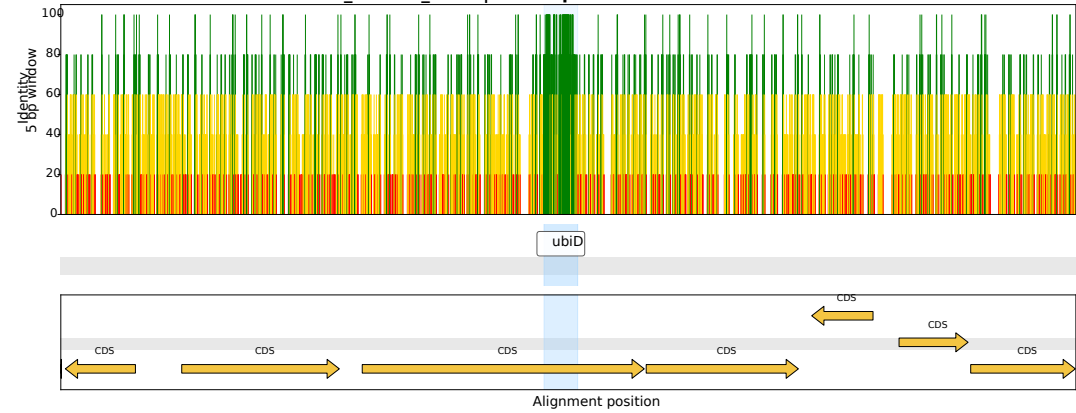

CI\_Harlain\_0022 | *Arsenophonus* CP123759.1

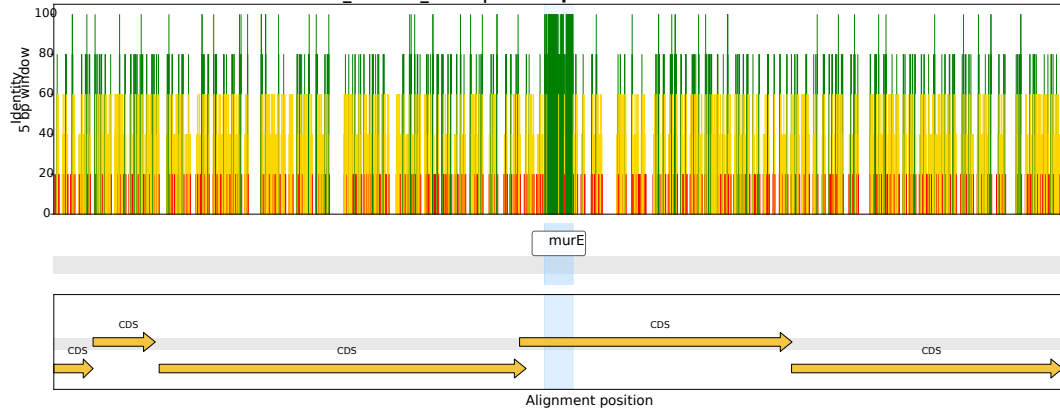

CI\_Harlain\_0378 | *Arsenophonus* OZ026540.1

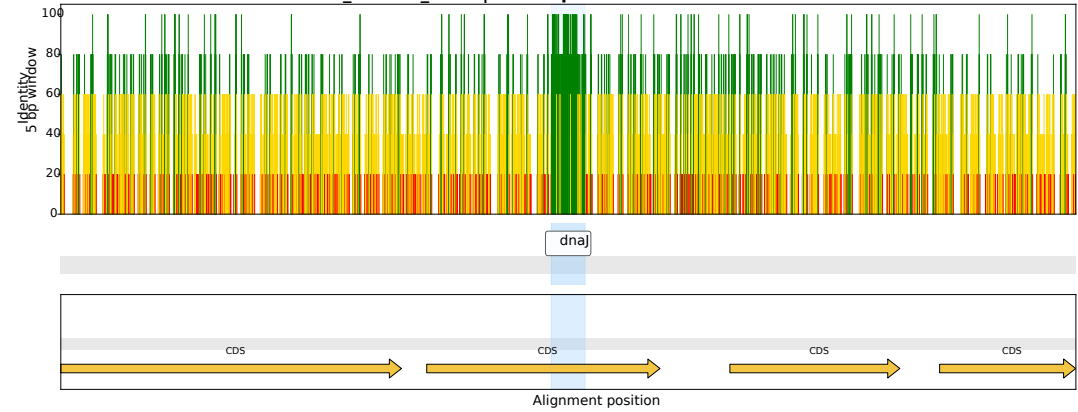

CI\_Harlain\_0039 | *Arsenophonus* CP123499.1

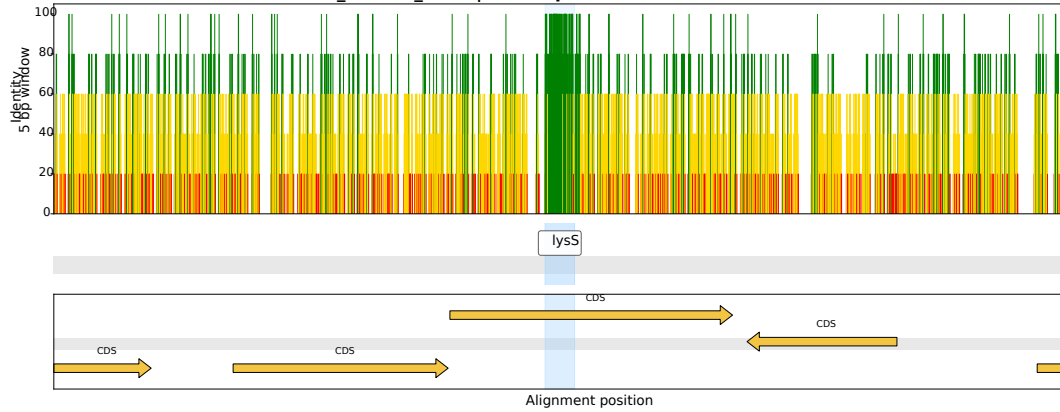

CI\_Harlain\_0052 | *Arsenophonus* CP123759.1

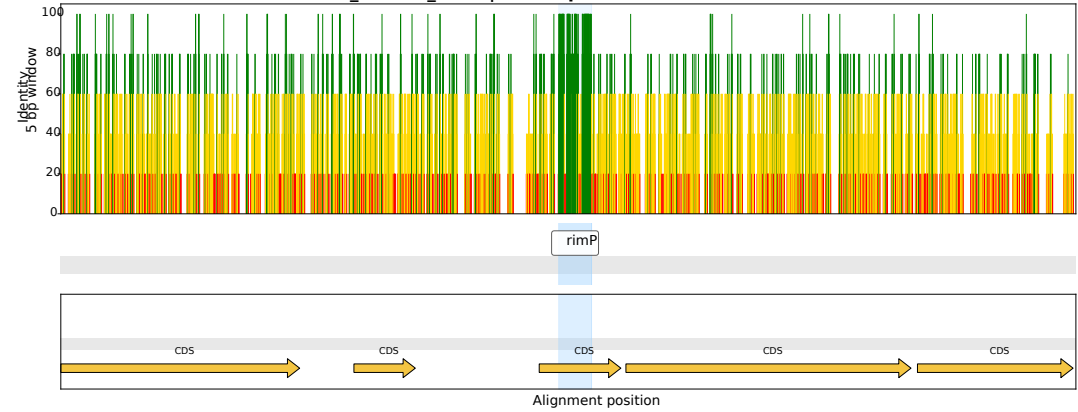

CI\_Harlain\_0136 | *Arsenophonus* OZ026540.1

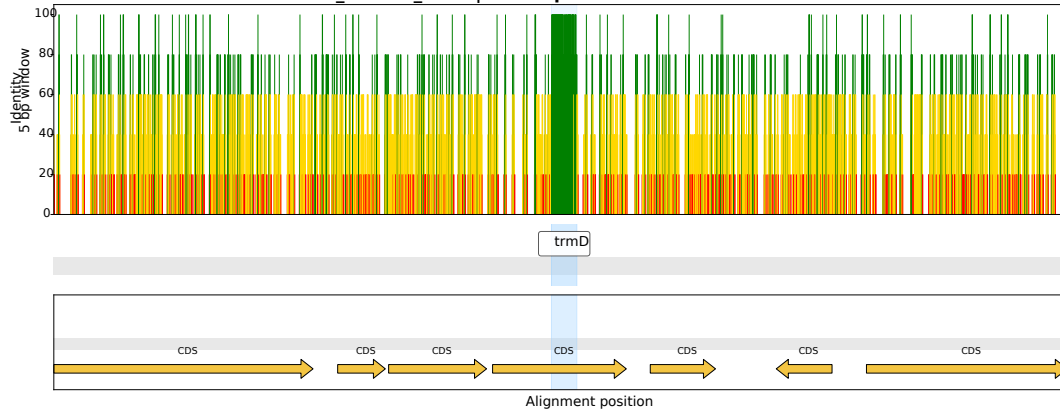

CI\_Harlain\_0032 | *Arsenophonus* CP038613.1

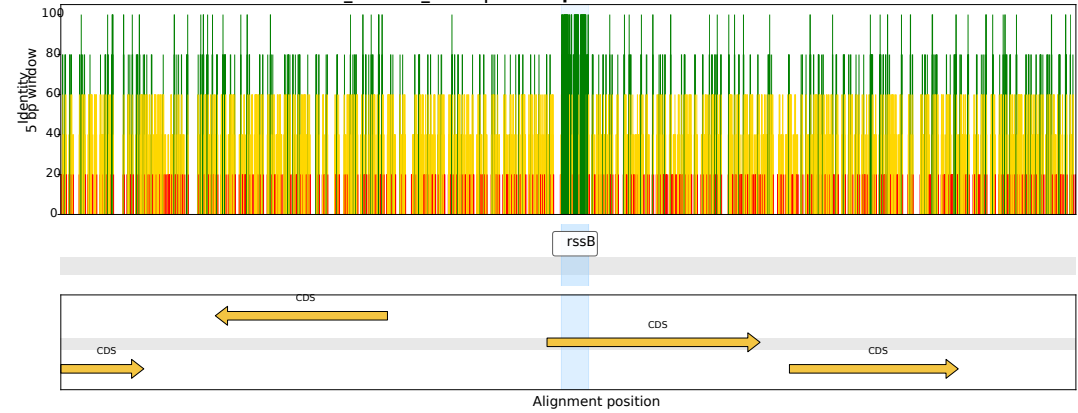

CI\_Harlain\_0196 | *Arsenophonus* CP123759.1

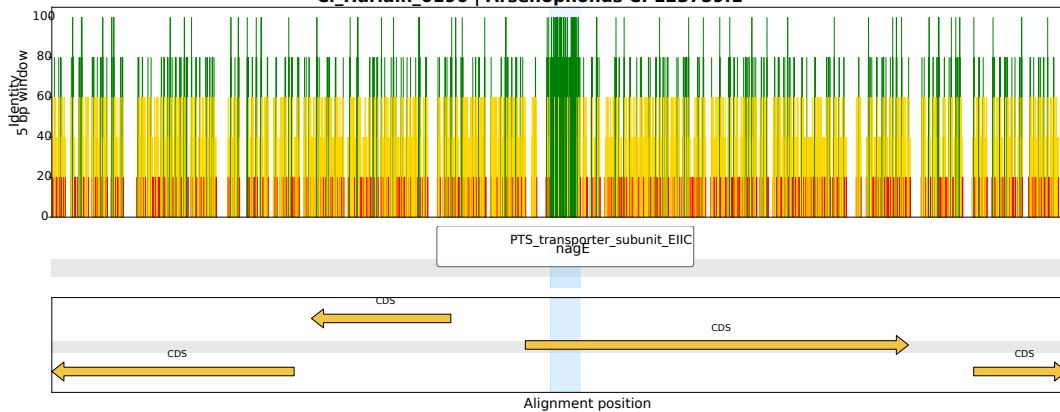

CI\_Harlain\_0001 | *Arsenophonus* CP038613.1

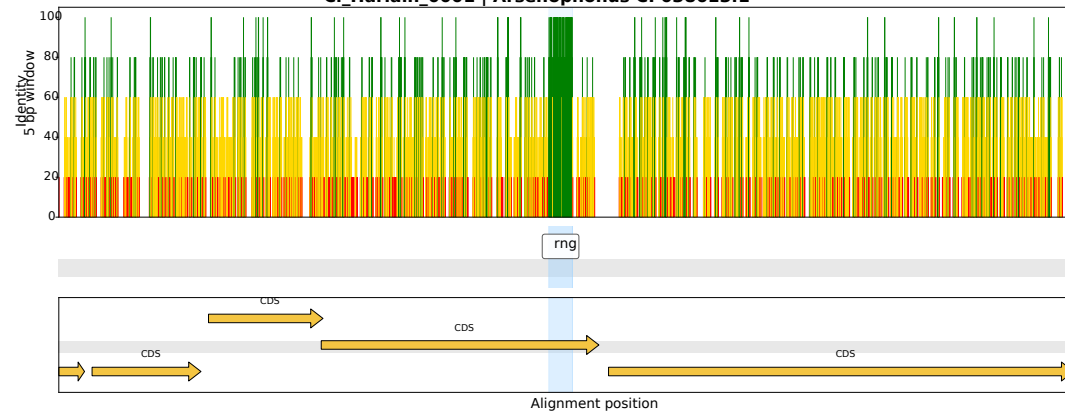

CI\_Harlain\_0120 | *Arsenophonus* CP123499.1

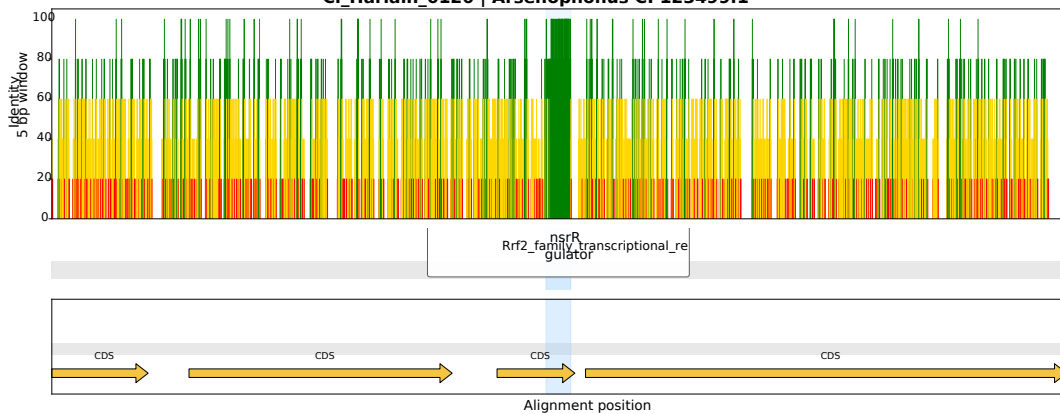

CI\_Harlain\_0390 | *Arsenophonus* OZ026540.1

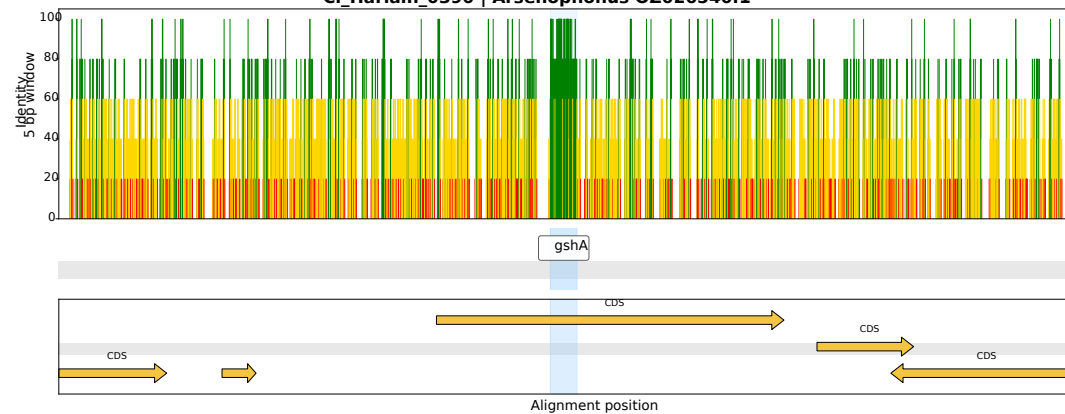

CI\_Harlain\_0024 | *Arsenophonus* OZ026540.1

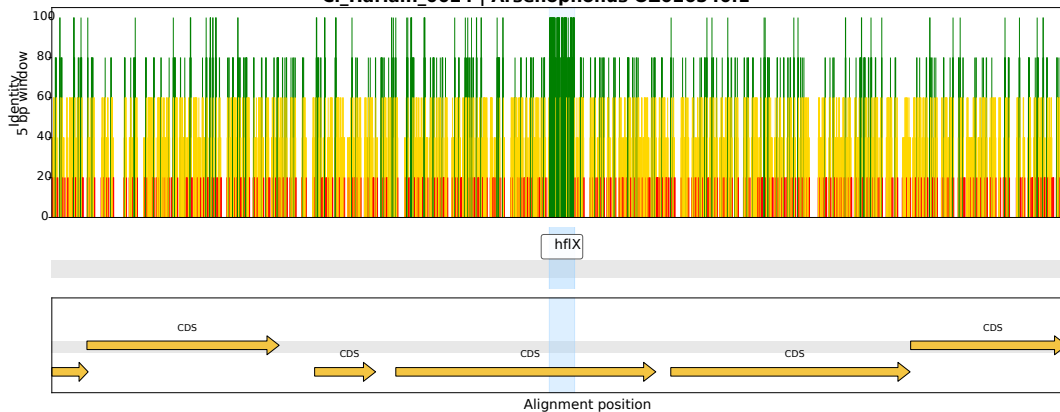

CI\_Harlain\_0252 | *Arsenophonus* OZ026540.1

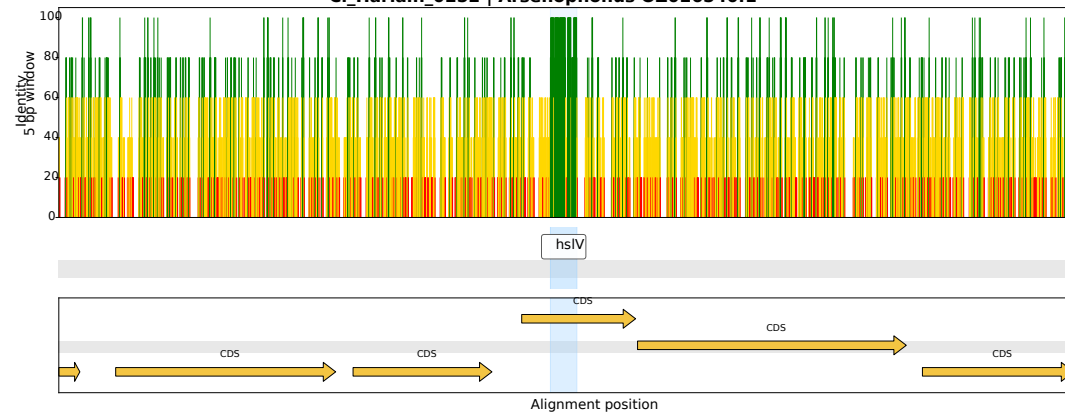

CI\_Harlain\_0231 | *Arsenophonus* CP123759.1

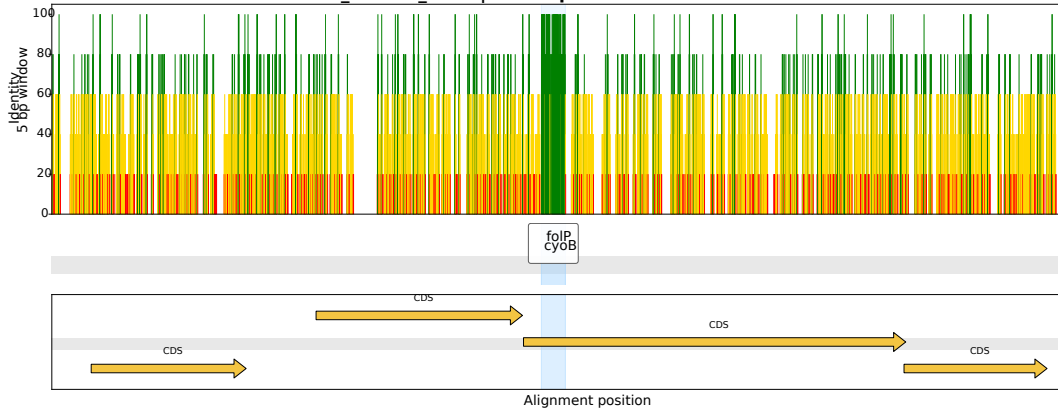

CI\_Harlain\_0241 | *Arsenophonus* CP084222.1

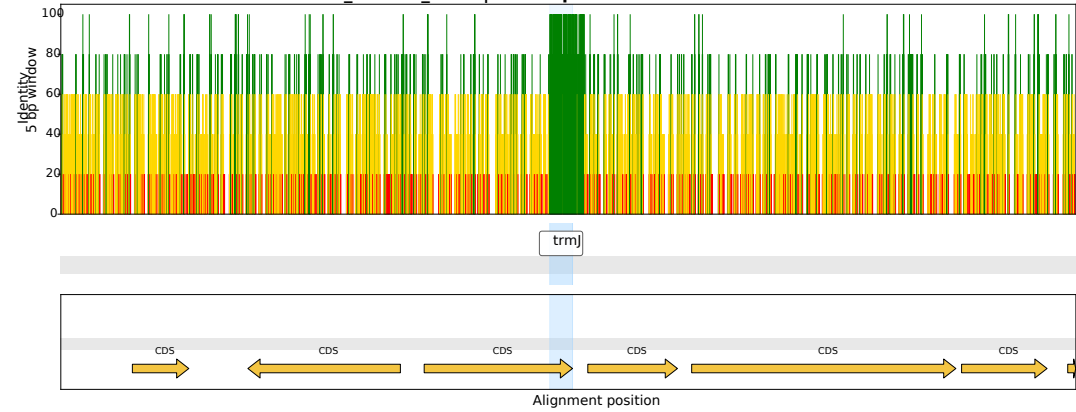

CI\_Harlain\_0159 | *Arsenophonus* CP123759.1

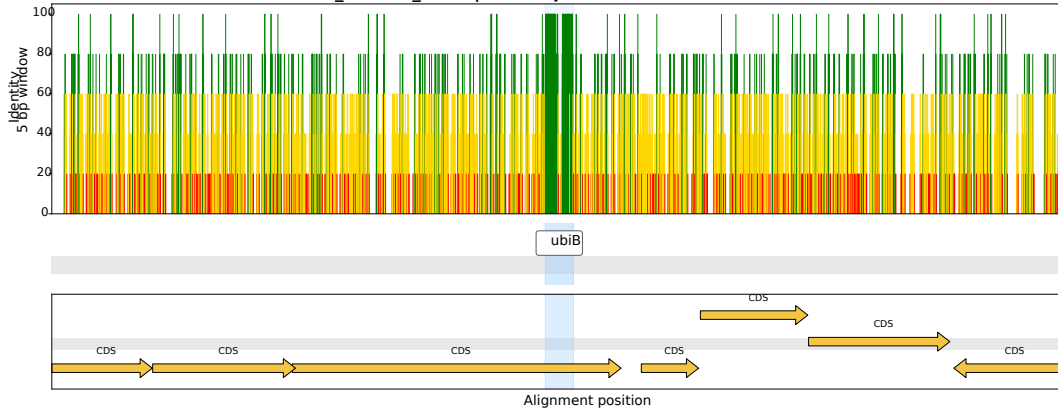

CI\_Harlain\_0300 | *Arsenophonus* OZ026540.1

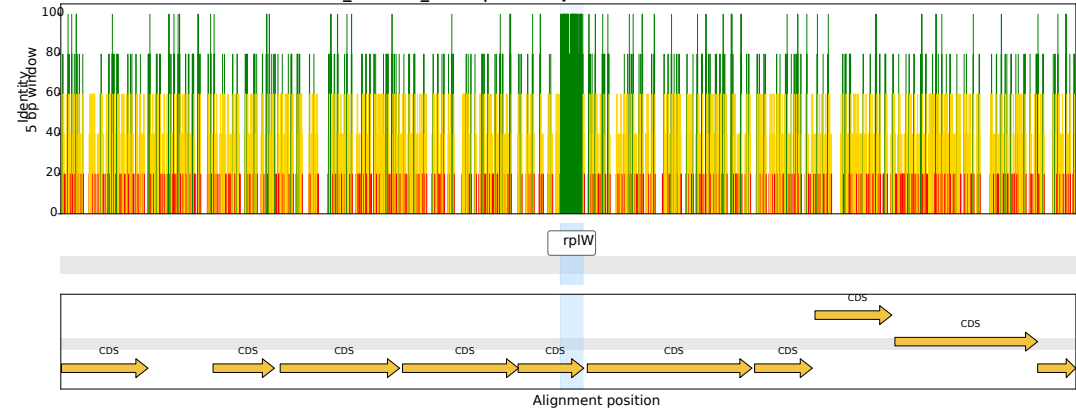

CI\_Harlain\_0219 | *Arsenophonus* OZ026540.1

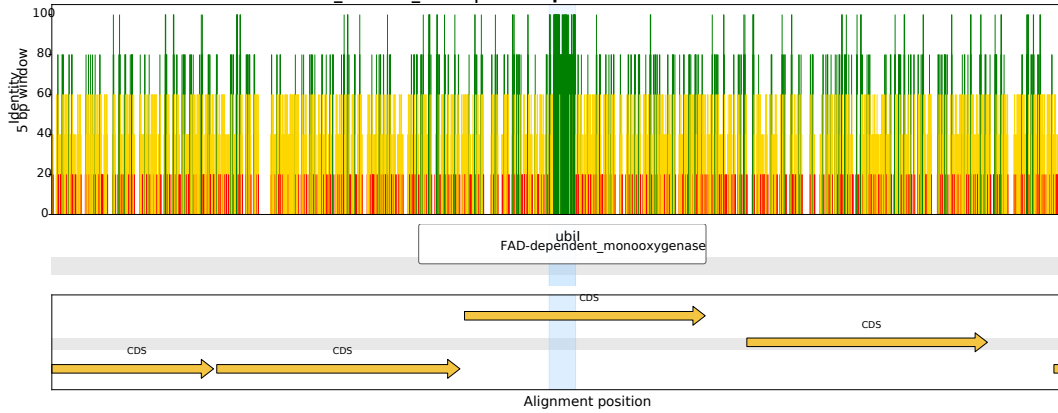

CI\_Harlain\_0114 | *Arsenophonus* CP123759.1

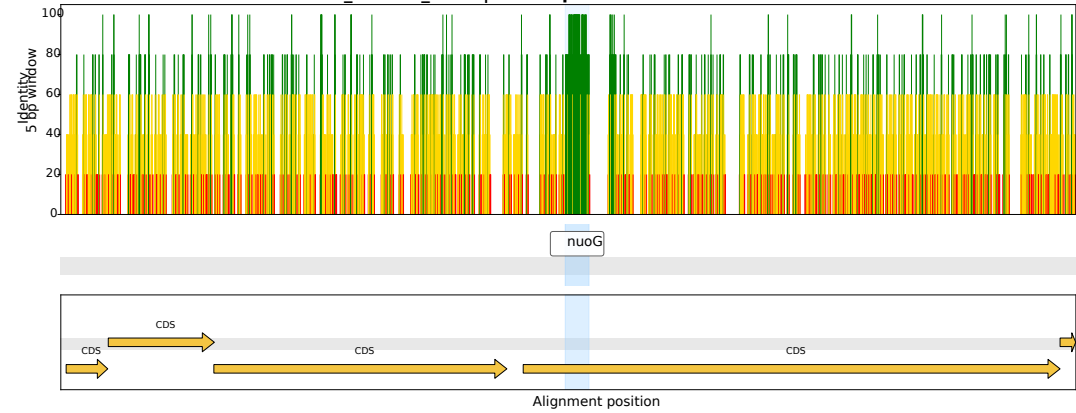

CI\_Harlain\_0081 | *Arsenophonus* CP123499.1

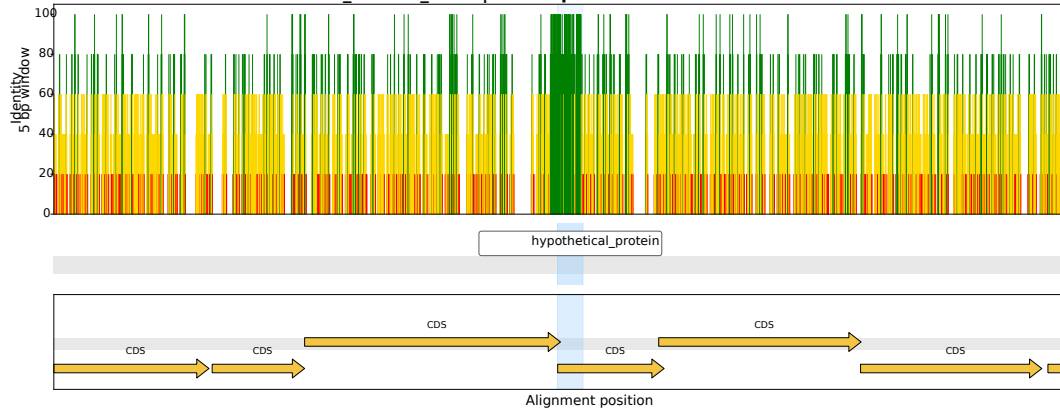

CI\_Harlain\_0004 | *Arsenophonus* CP038155.1

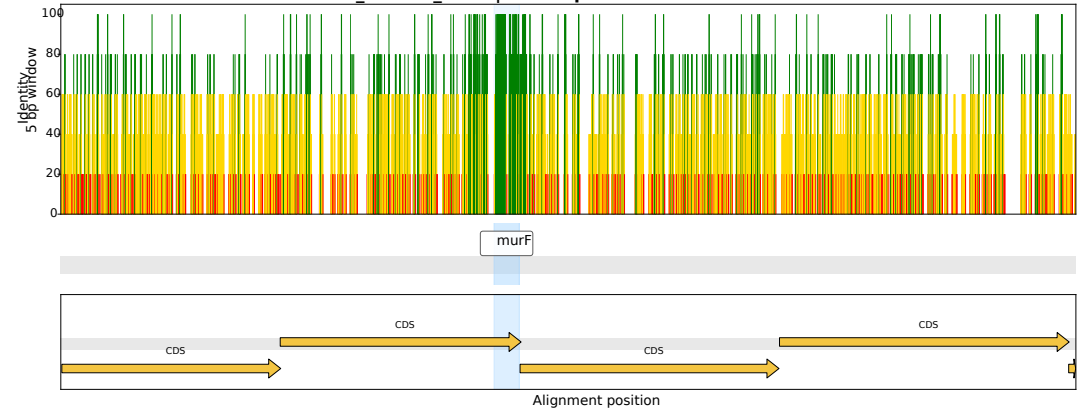

CI\_Harlain\_0123 | *Arsenophonus* OZ026540.1

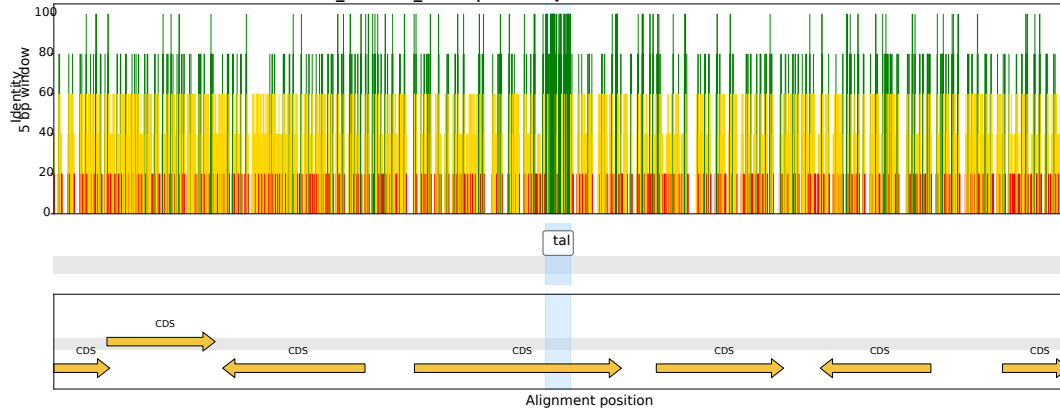

CI\_Harlain\_0065 | *Arsenophonus* CP038613.1

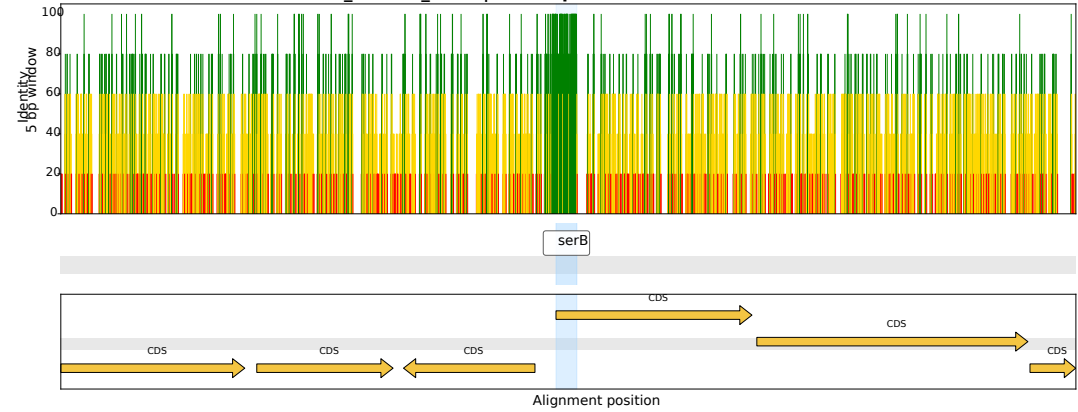

CI\_Harlain\_0262 | *Arsenophonus* CP084222.1

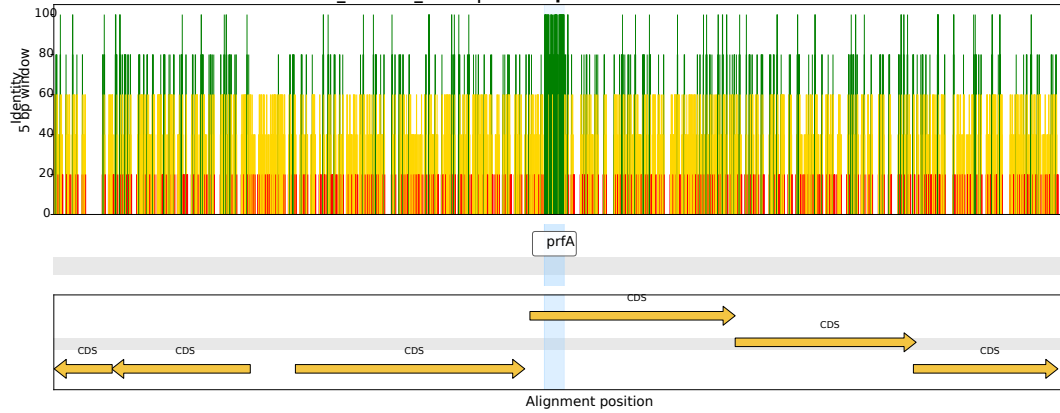

CI\_Harlain\_0005 | *Arsenophonus* CP038155.1

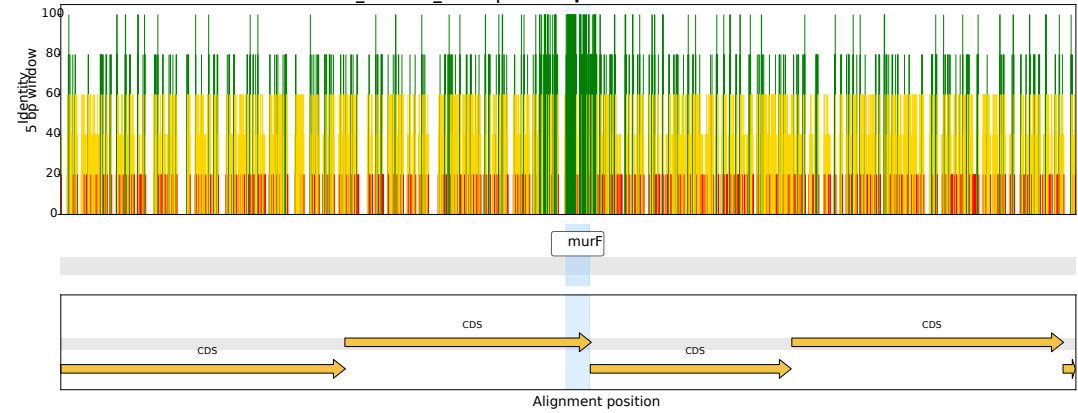

CI\_Harlain\_0450 | *Arsenophonus* CP123759.1

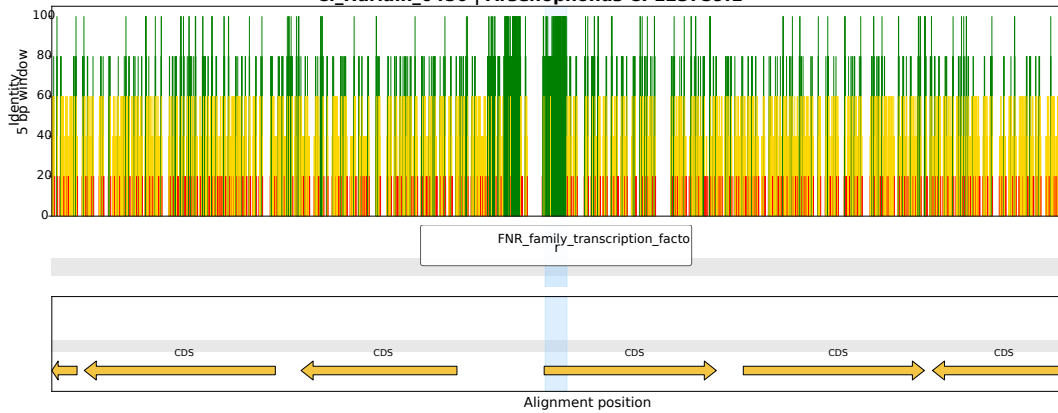

CI\_Harlain\_0151 | *Arsenophonus* CP123499.1

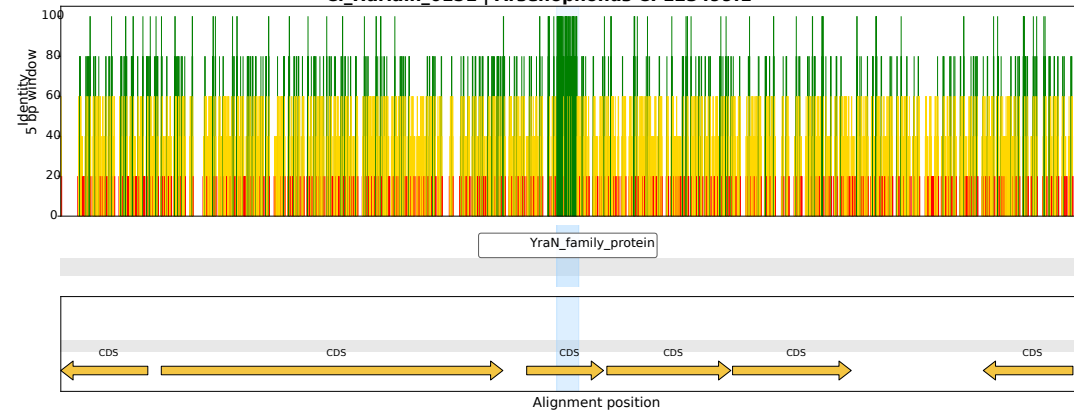

CI\_Harlain\_0423 | *Arsenophonus* OZ026540.1

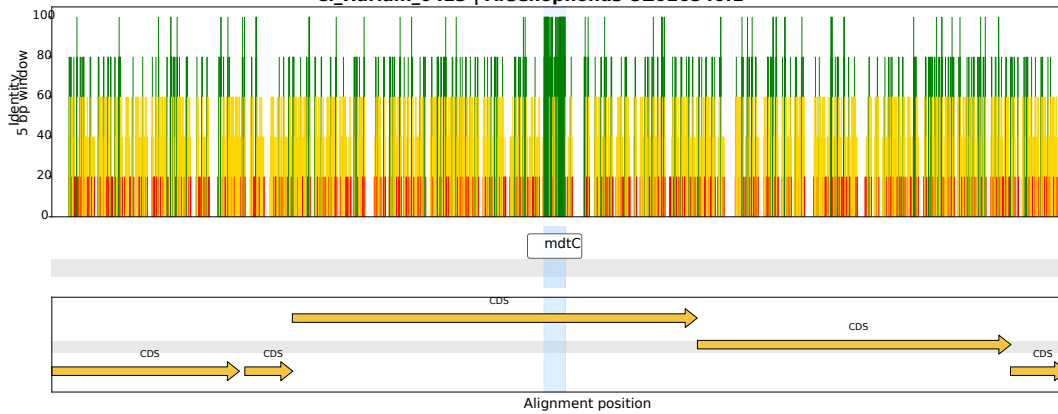

CI\_Harlain\_0111 | *Arsenophonus* CP013920.1

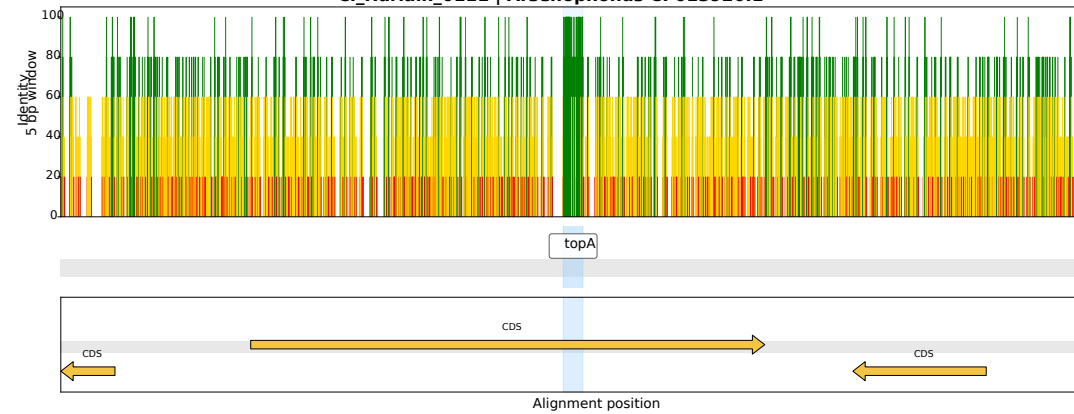

CI\_Harlain\_0221 | *Arsenophonus* CP084222.1

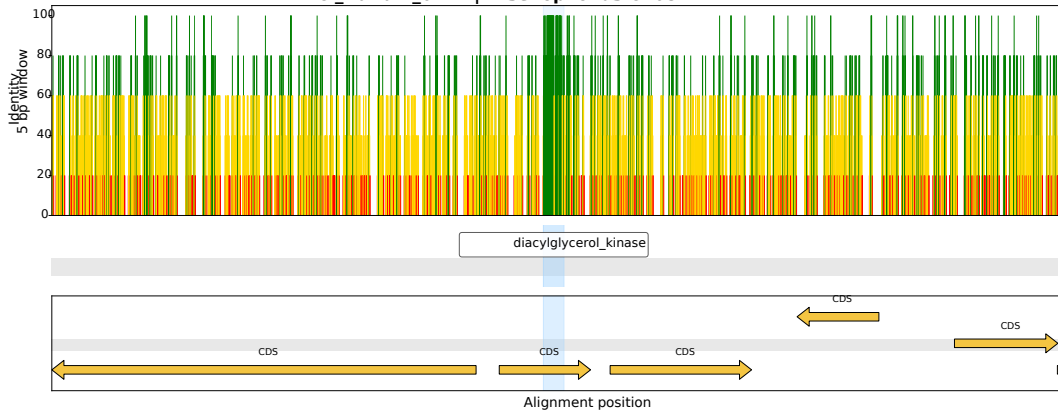

CI\_Harlain\_0237 | *Arsenophonus* CP038613.1

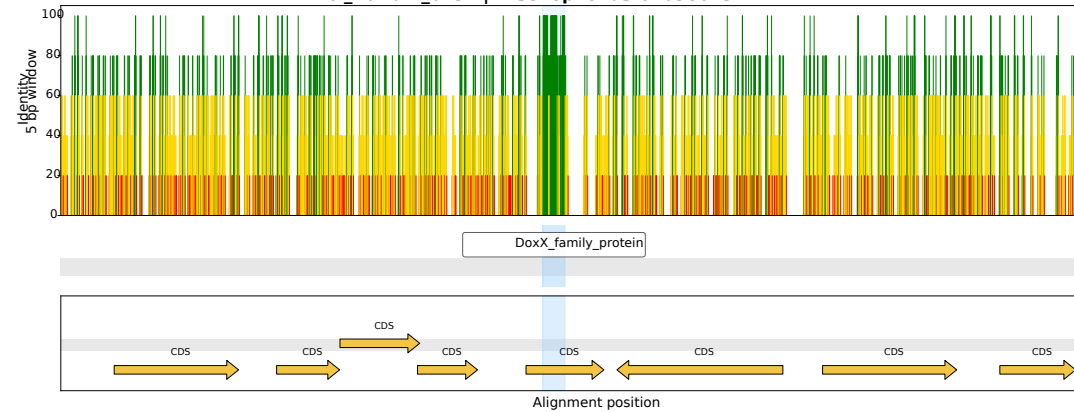

CI\_Harlain\_0351 | *Arsenophonus* OZ026540.1

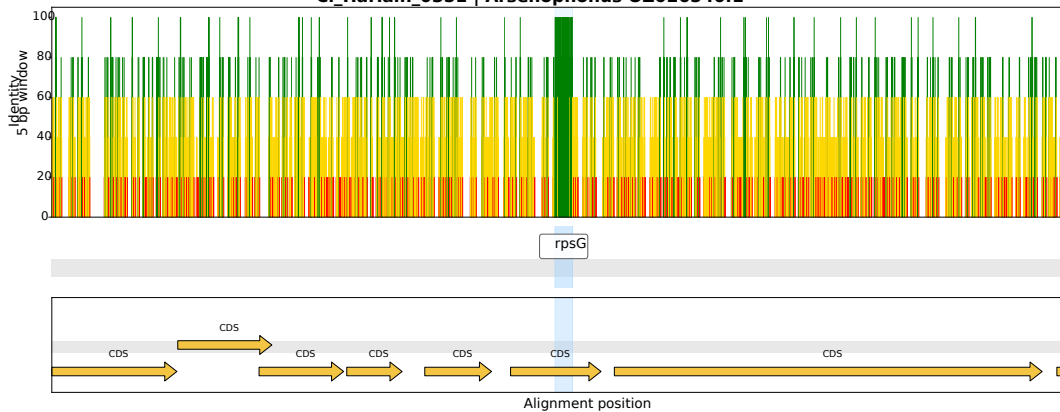

CI\_Harlain\_0069 | *Arsenophonus* CP038155.1

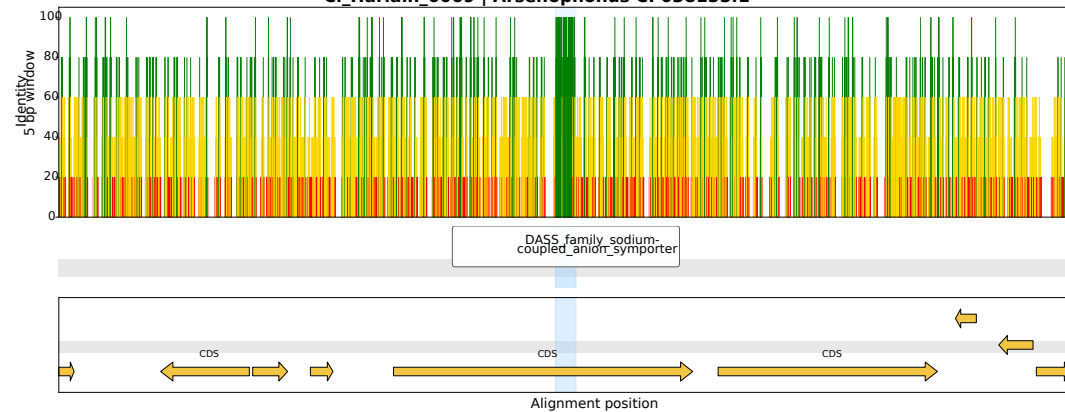

CI\_Harlain\_0291 | *Arsenophonus* CP038613.1

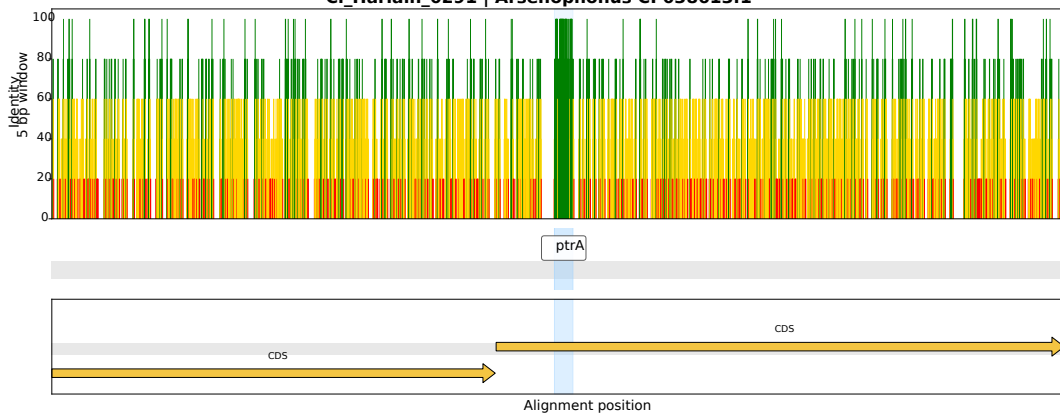

CI\_Harlain\_0333 | *Arsenophonus* OZ026540.1

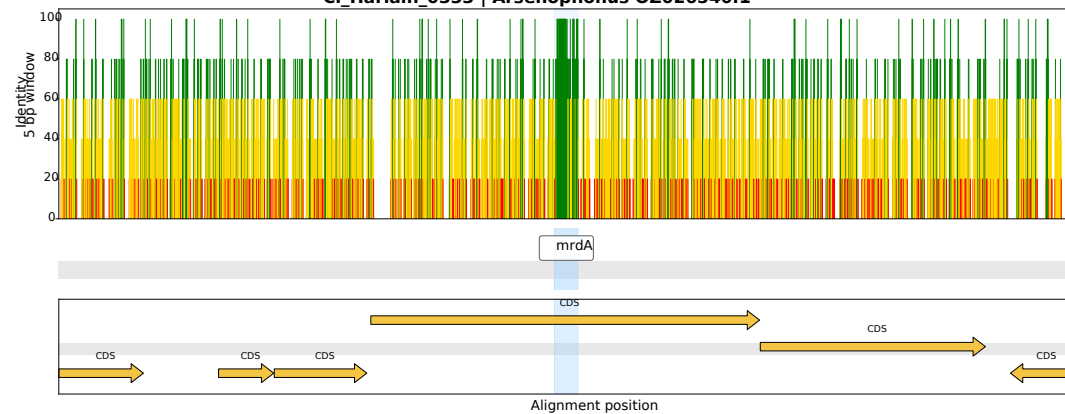

CI\_Harlain\_0349 | *Arsenophonus* CP038613.1

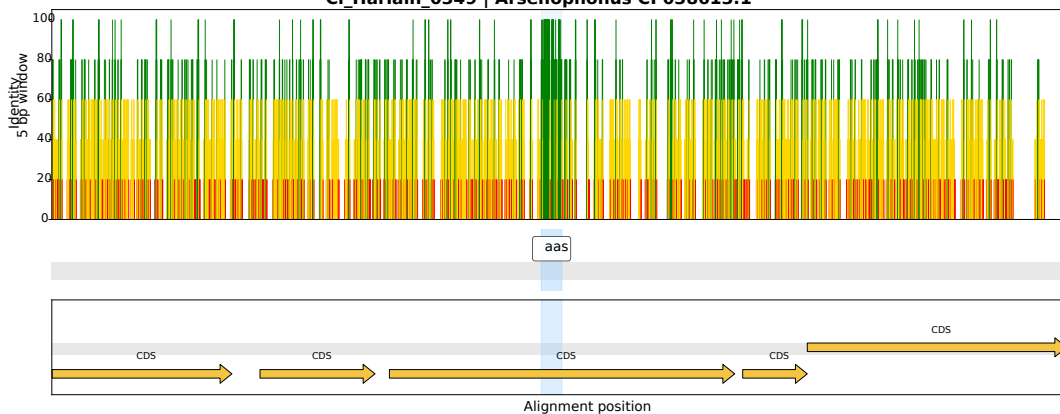

CI\_Harlain\_0204 | *Arsenophonus* CP123759.1

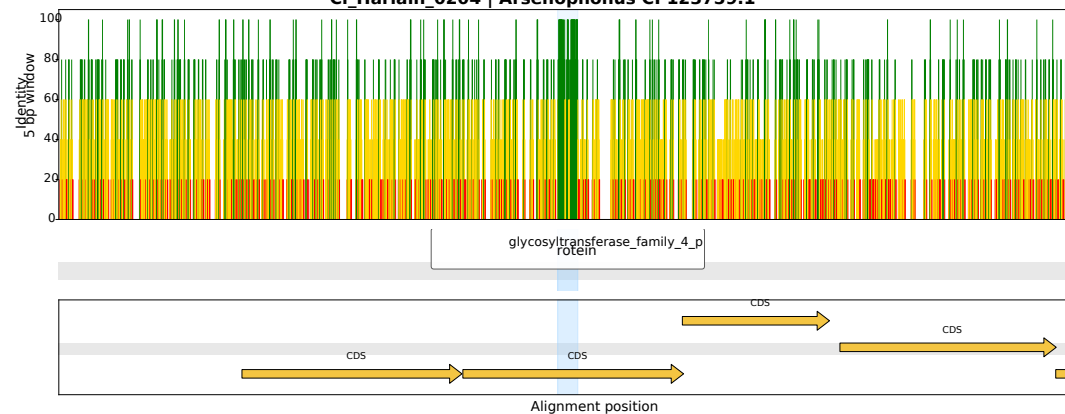

CI\_Harlain\_0278 | *Arsenophonus* OZ026540.1

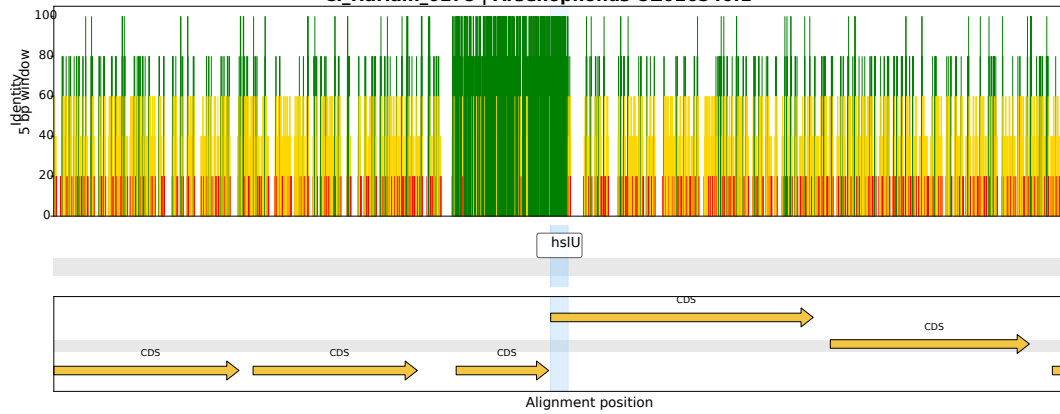

CI\_Harlain\_0297 | *Arsenophonus* CP038614.1

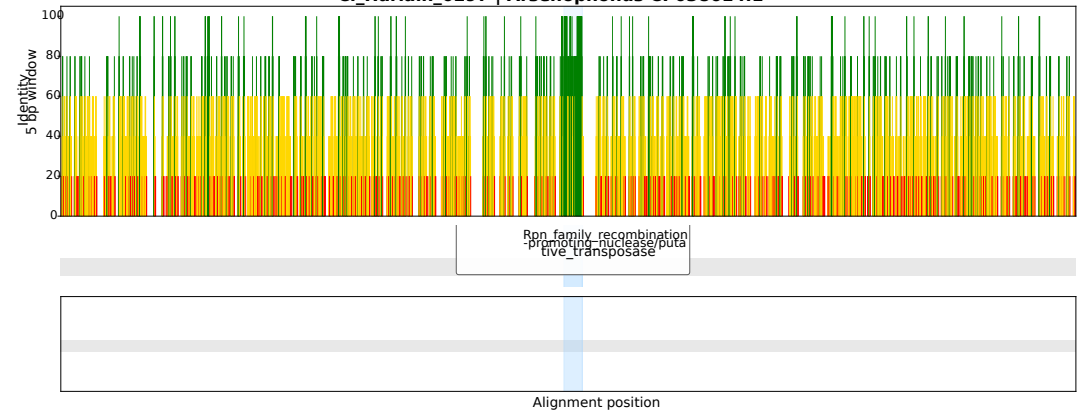

CI\_Harlain\_0084 | *Arsenophonus* OZ026540.1

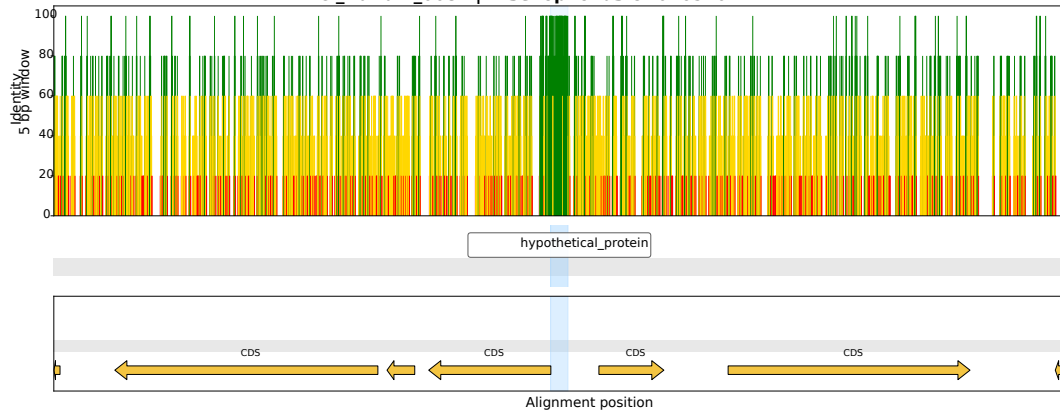

CI\_Harlain\_0187 | *Arsenophonus* CP038613.1

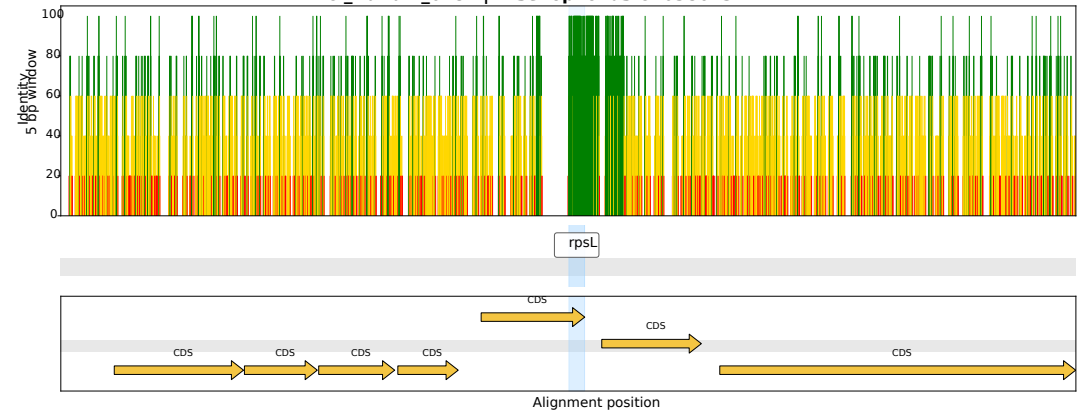

CI\_Harlain\_0433 | *Arsenophonus* OZ026540.1

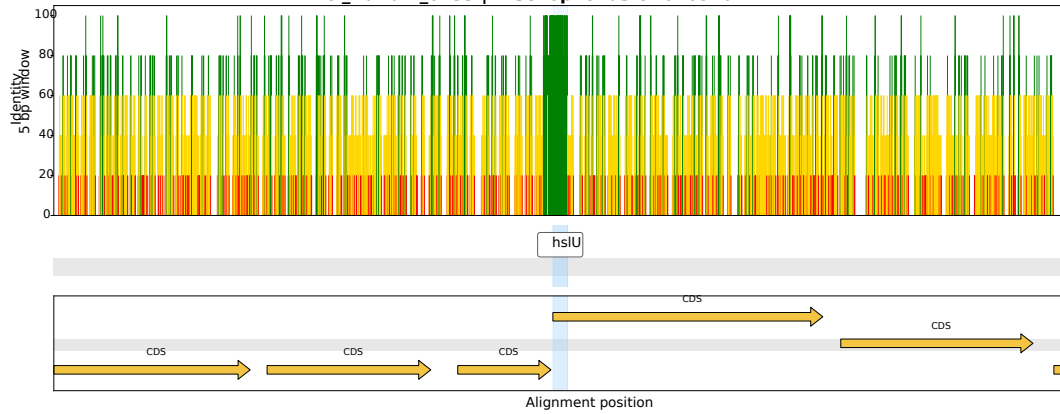

CI\_Harlain\_0368 | *Arsenophonus* CP123499.1

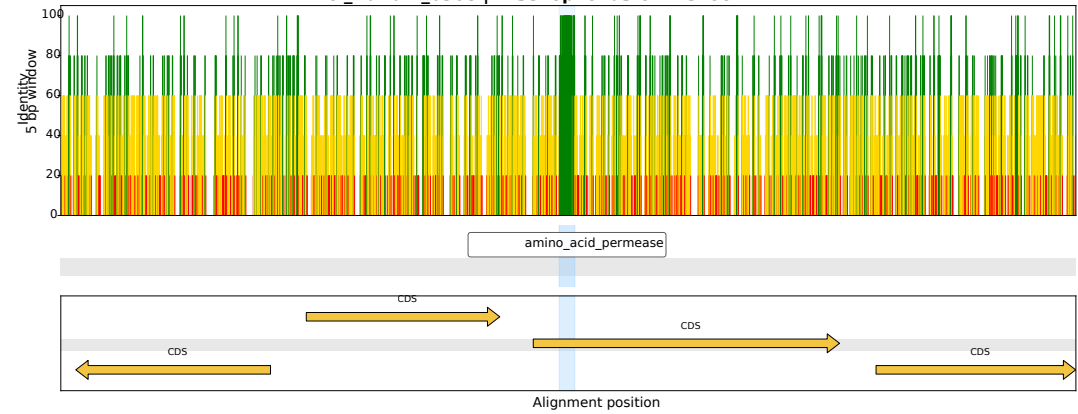

CI\_Harlain\_0058 | *Arsenophonus* CP123759.1

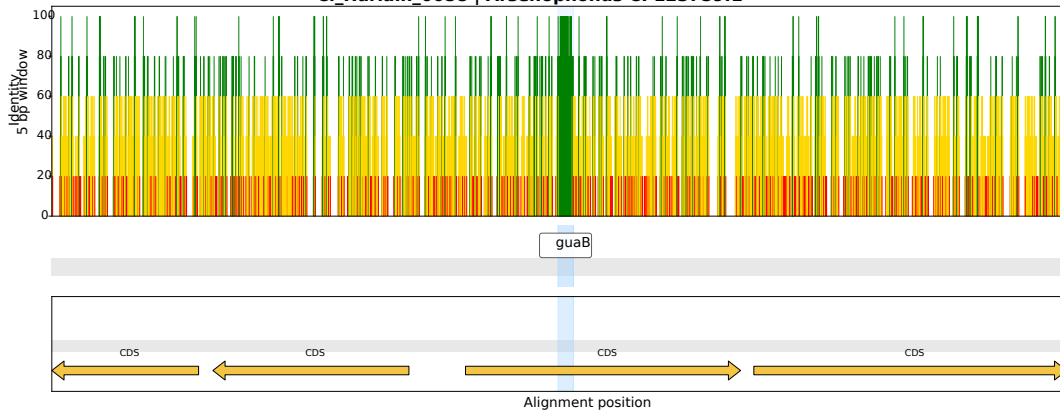

CI\_Harlain\_0317 | *Arsenophonus* OZ026540.1

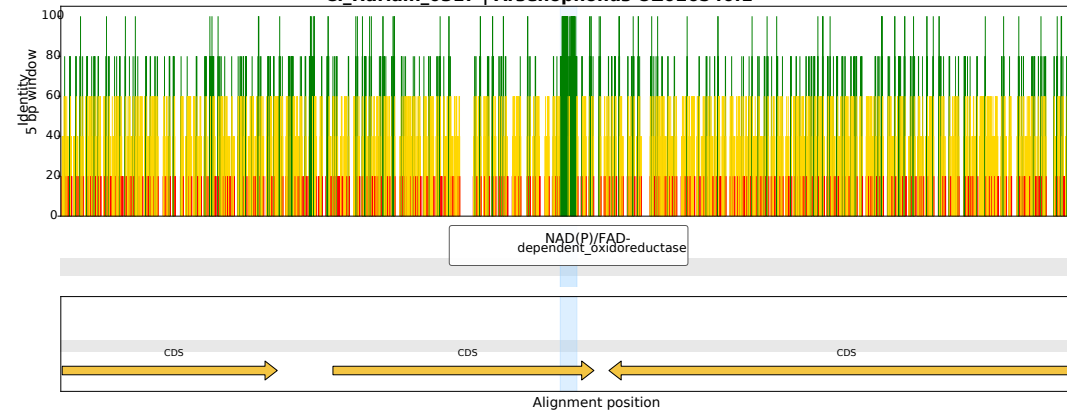

CI\_Harlain\_0198 | *Arsenophonus* CP038613.1

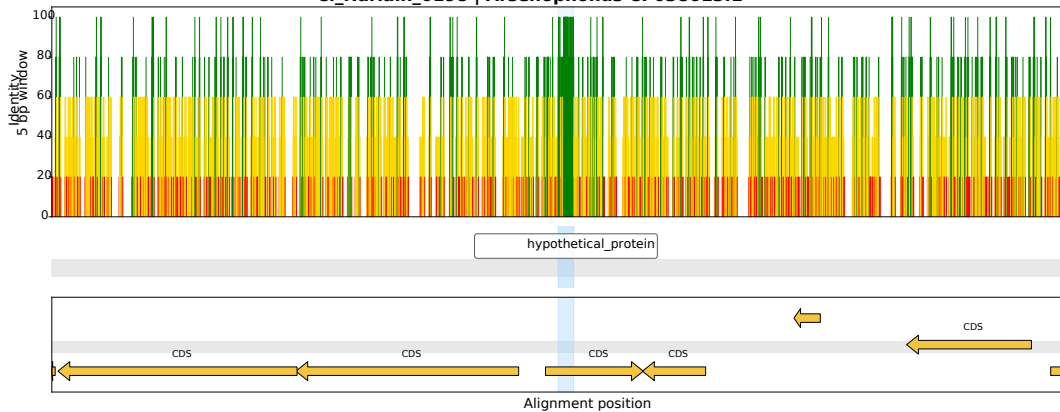

CI\_Harlain\_0086 | *Arsenophonus* OZ026540.1

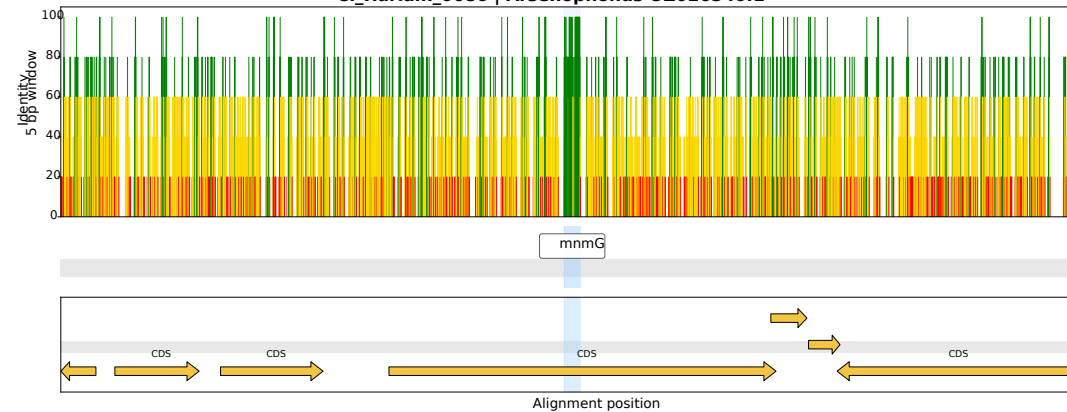

CI\_Harlain\_0200 | *Arsenophonus* CP038613.1

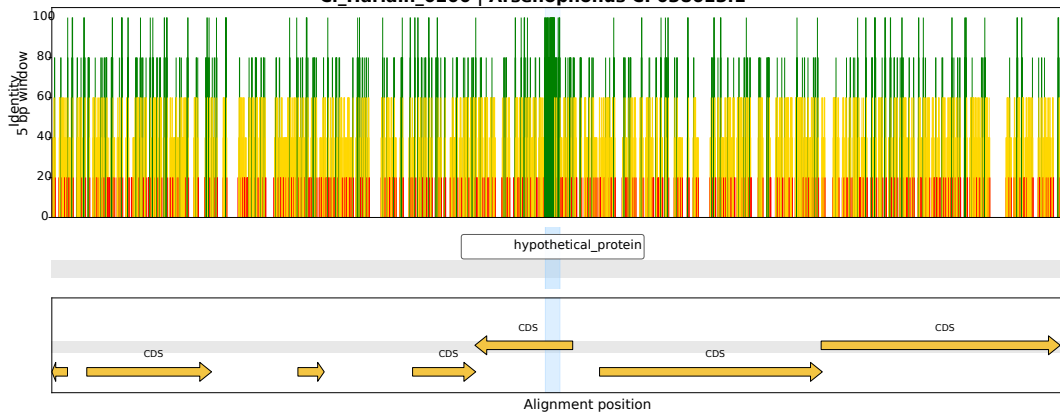

CI\_Harlain\_0282 | *Arsenophonus* CP123759.1

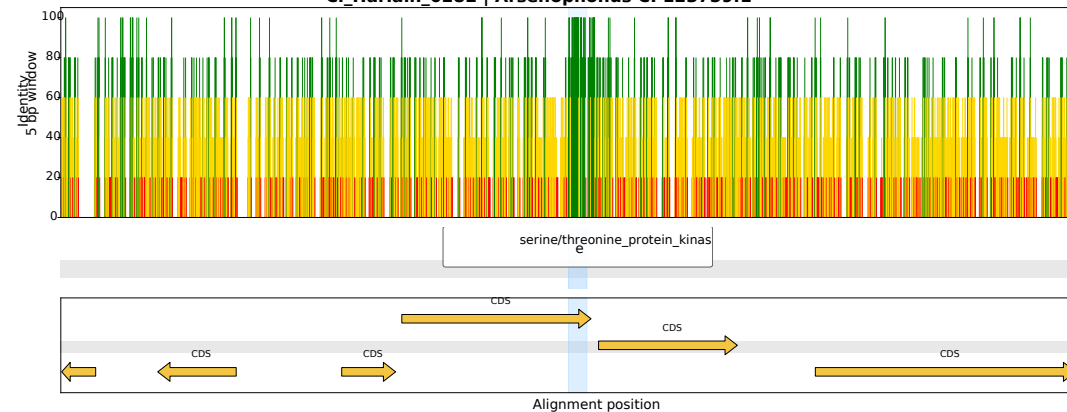

CI\_Harlain\_0409 | *Arsenophonus* CP123499.1

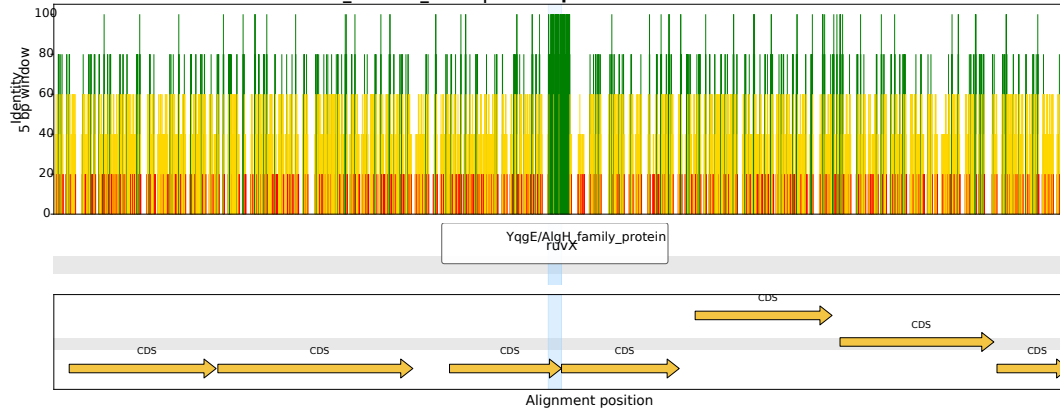

CI\_Harlain\_0471 | *Arsenophonus* CP038613.1

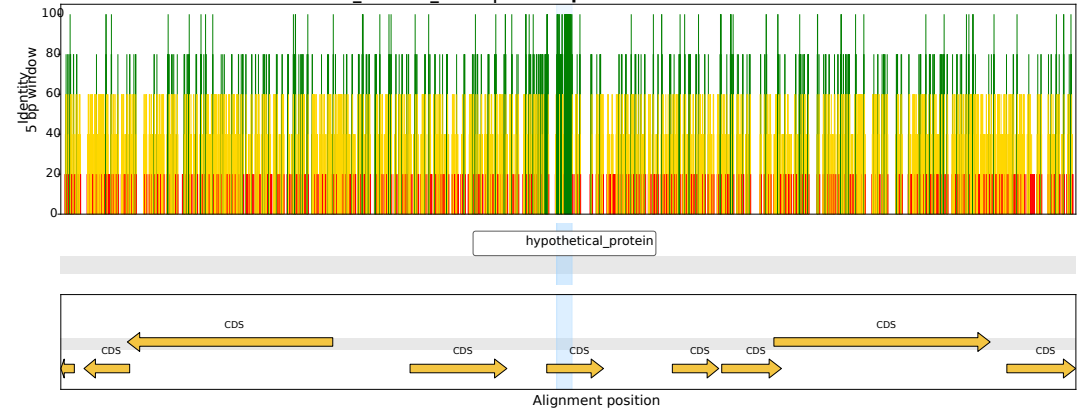

CI\_Harlain\_0365 | *Arsenophonus* CP038613.1

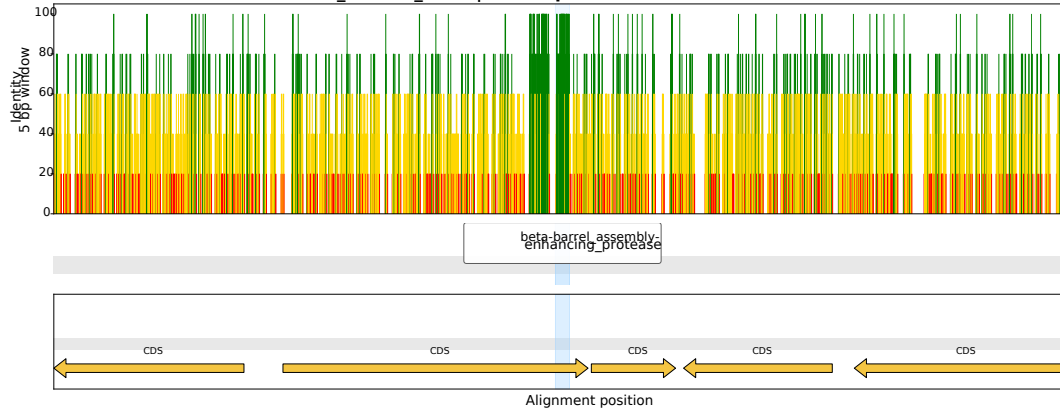

CI\_Harlain\_0156 | *Arsenophonus* CP123759.1

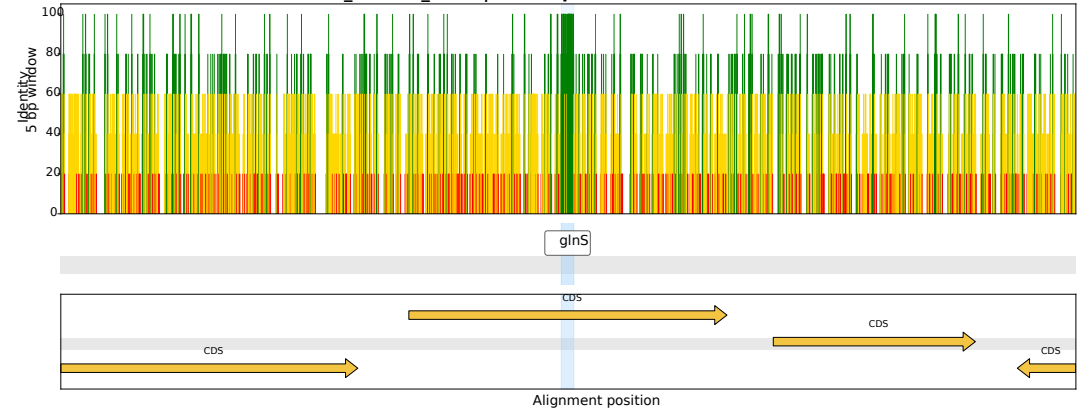

CI\_Harlain\_0064 | *Arsenophonus* CP038613.1

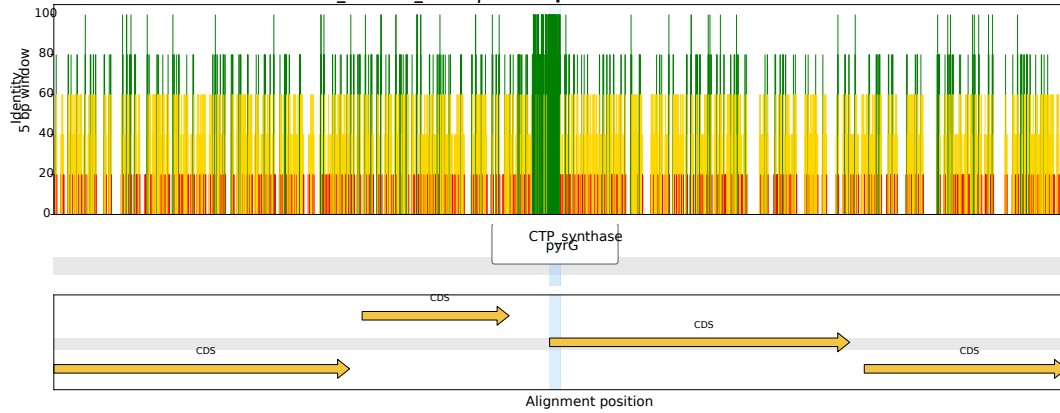

CI\_Harlain\_0186 | *Arsenophonus* CP123499.1

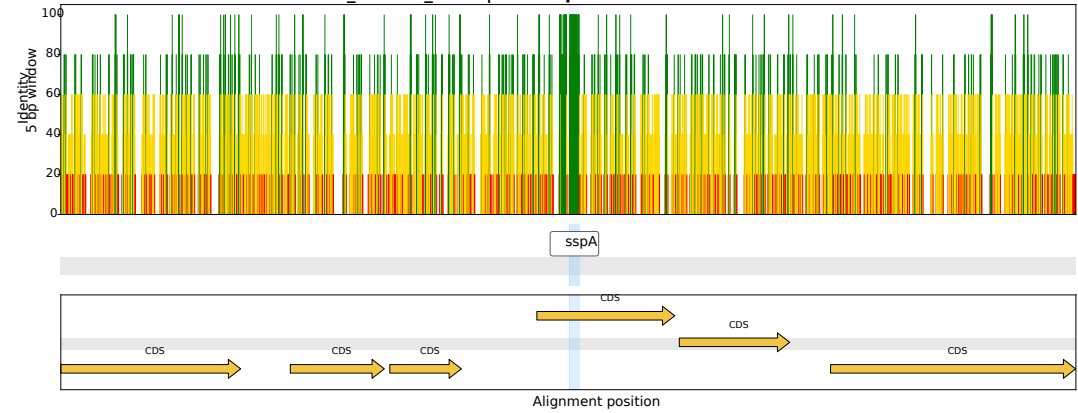

CI\_Harlain\_0194 | *Arsenophonus* CP038613.1

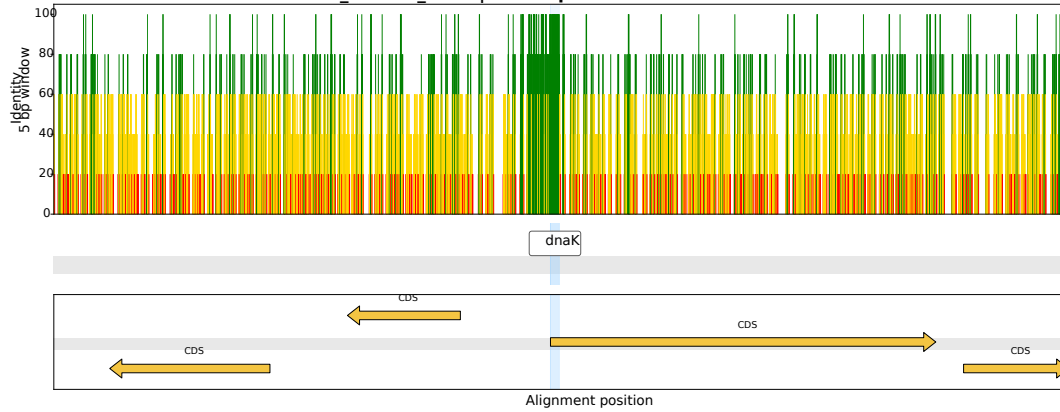

CI\_Harlain\_0034 | *Arsenophonus* CP123499.1

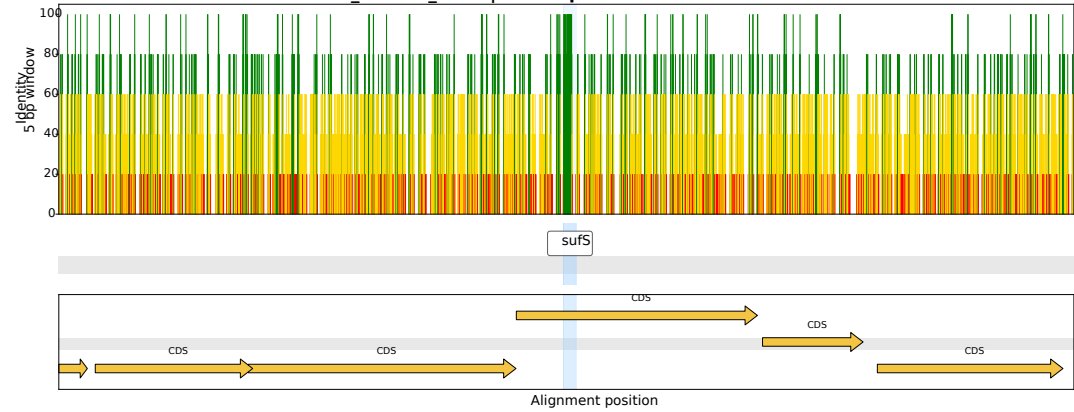

CI\_Harlain\_0255 | *Arsenophonus* OZ026540.1

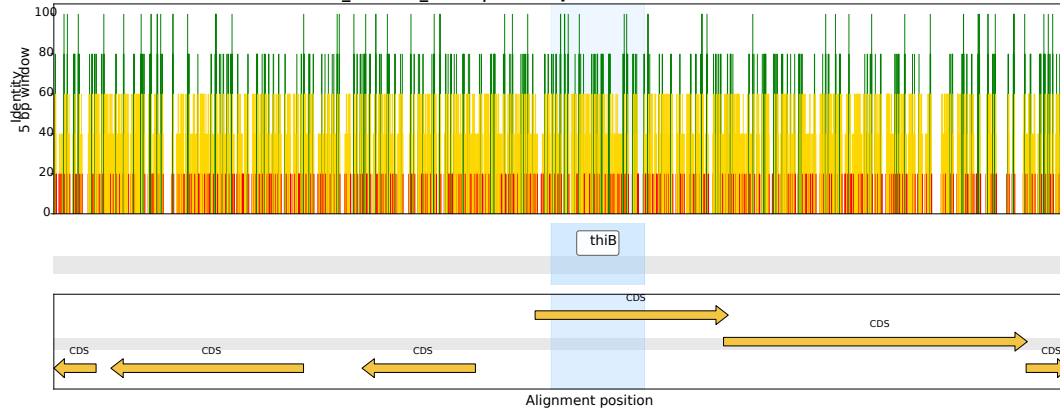

CI\_Harlain\_0105 | *Arsenophonus* CP038155.1

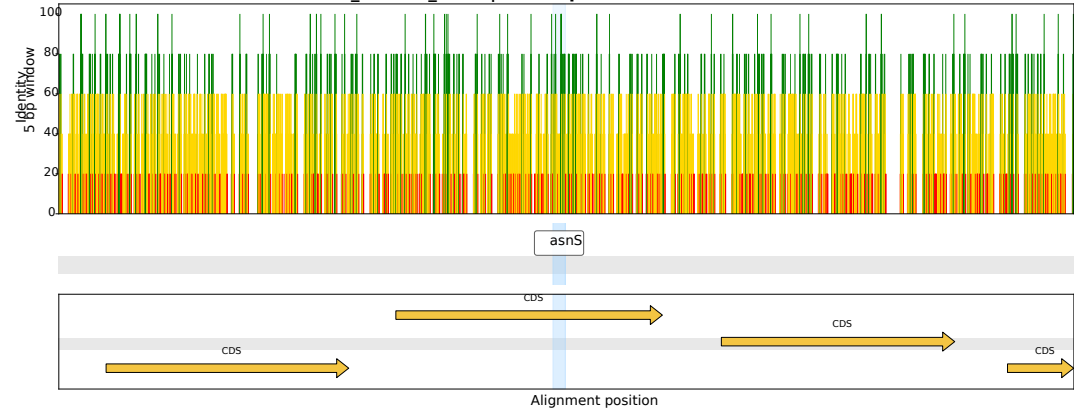

CI\_Harlain\_0399 | *Arsenophonus* OZ026540.1

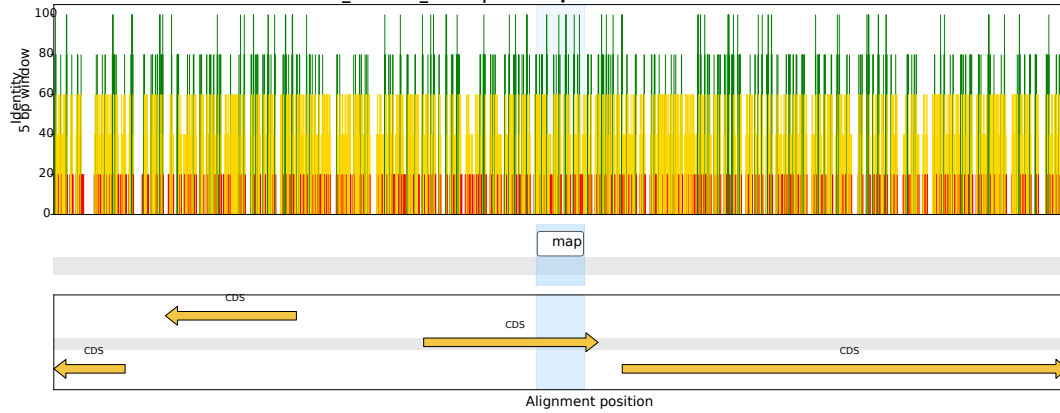

CI\_Harlain\_0031 | *Arsenophonus* OZ026540.1

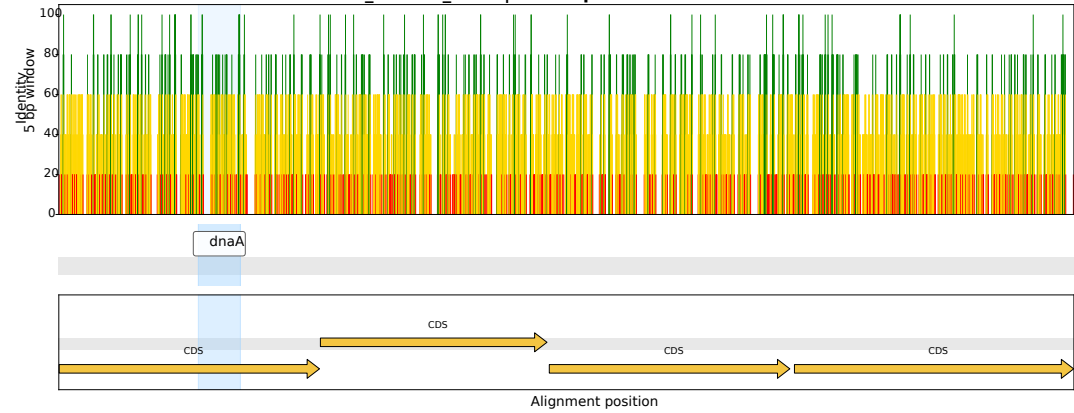

CI\_Harlain\_0382 | *Arsenophonus* CP038155.1

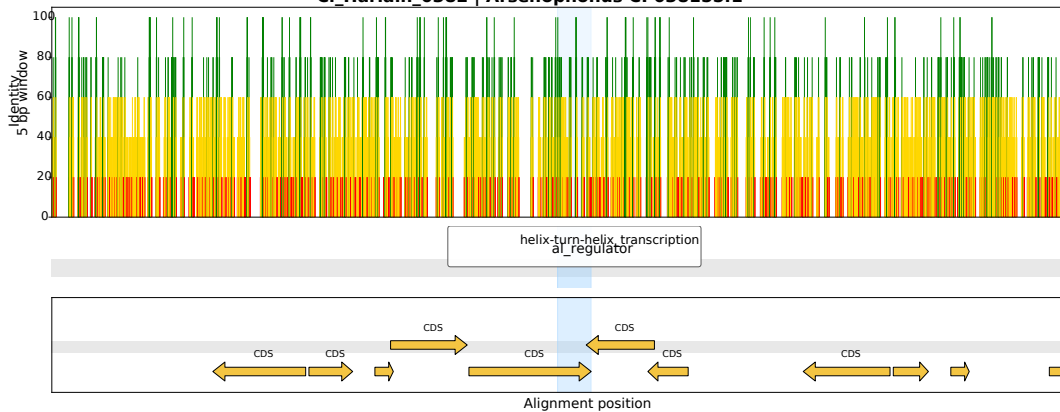

CI\_Harlain\_0459 | *Arsenophonus* CP123759.1

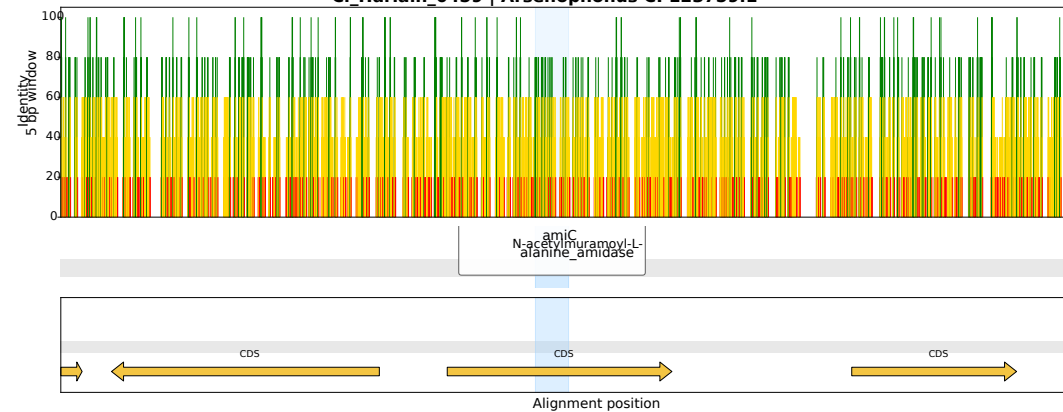

CI\_Harlain\_0320 | *Arsenophonus* CP123499.1

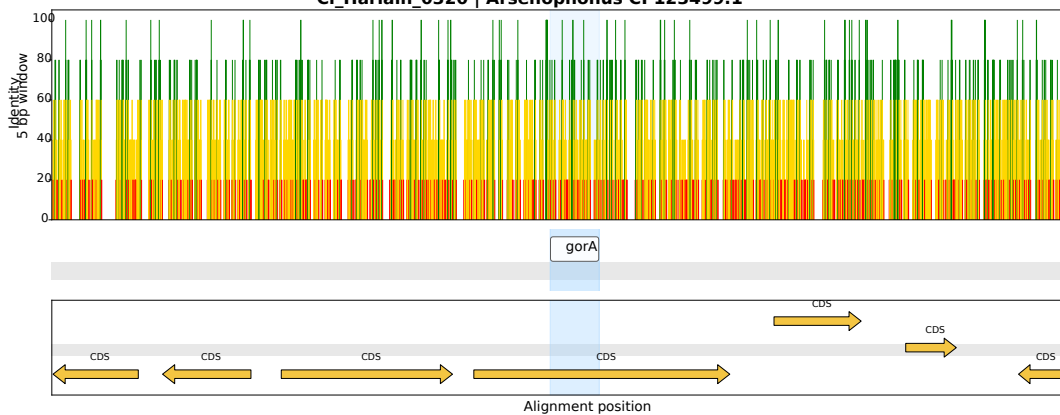

CI\_Harlain\_0293 | *Arsenophonus* OZ026540.1

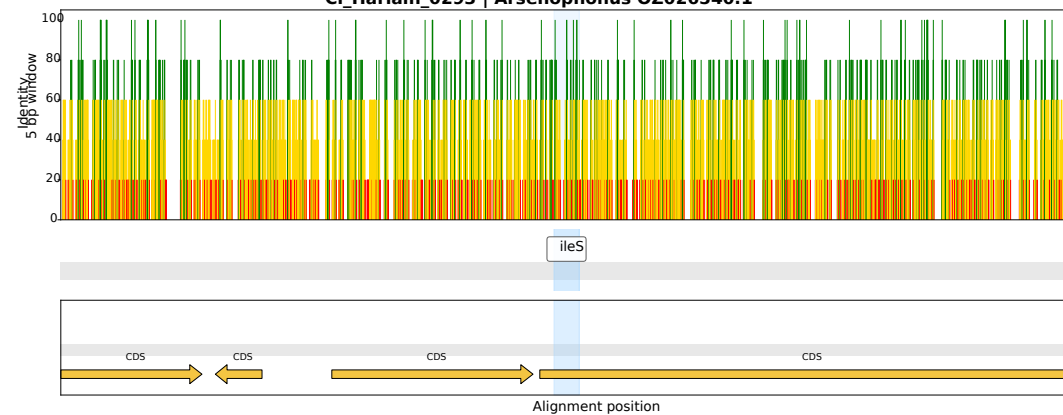

CI\_Harlain\_0384 | *Arsenophonus* CP038613.1

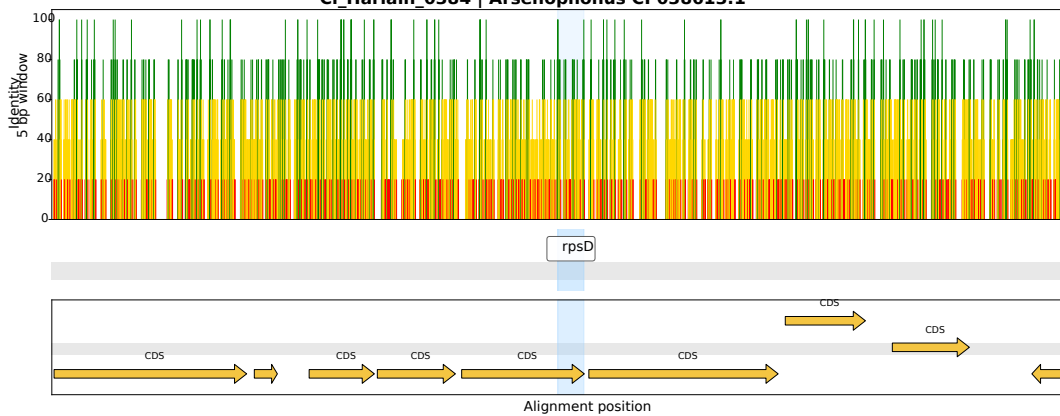

CI\_Harlain\_0054 | *Arsenophonus* CP038613.1

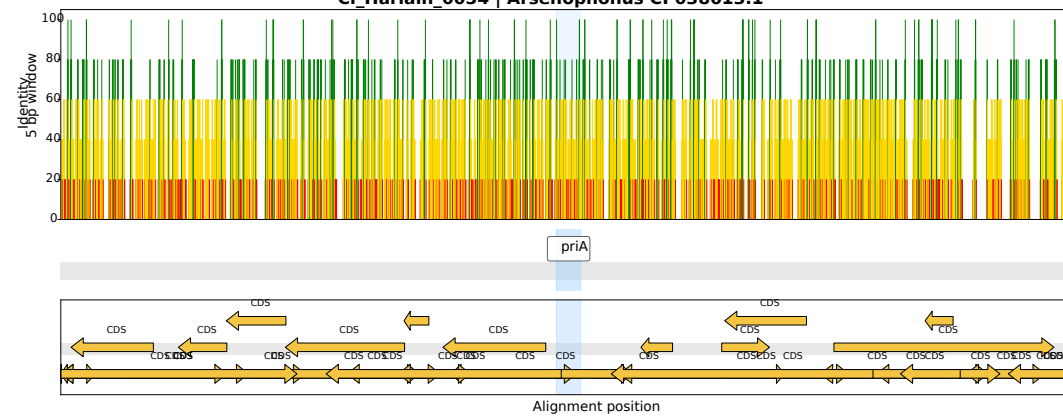

CI\_Harlain\_0009 | *Arsenophonus* CP038613.1

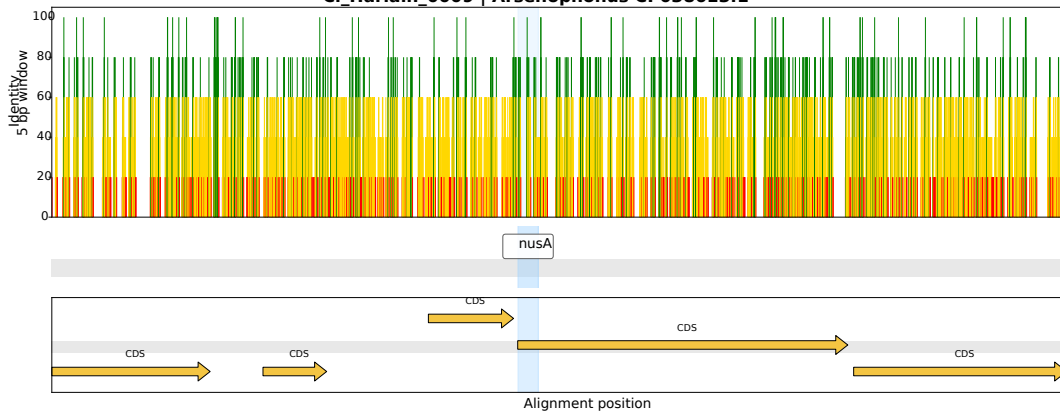

CI\_Harlain\_0134 | *Arsenophonus* OZ026540.1

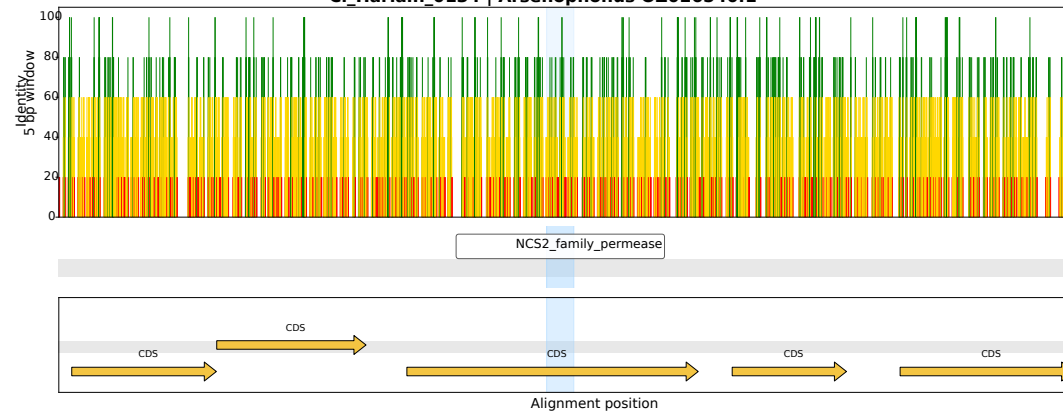

CI\_Harlain\_0220 | *Arsenophonus* CP038613.1

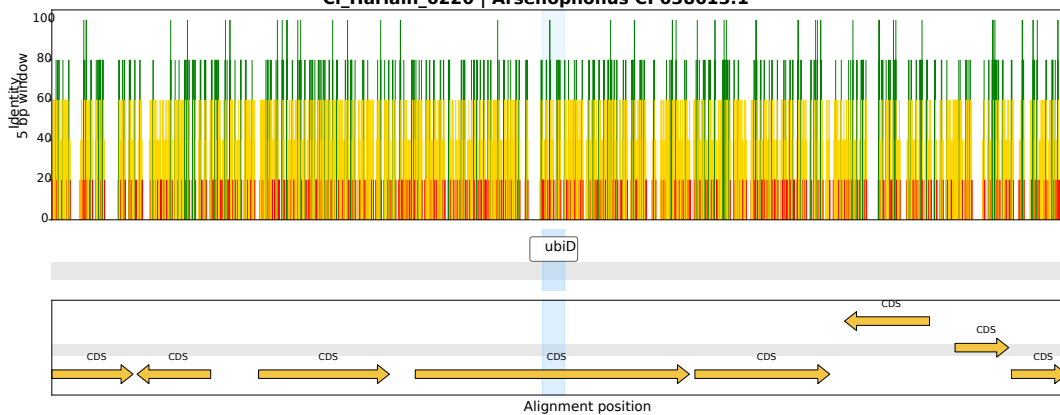

CI\_Harlain\_0429 | *Arsenophonus* CP038613.1

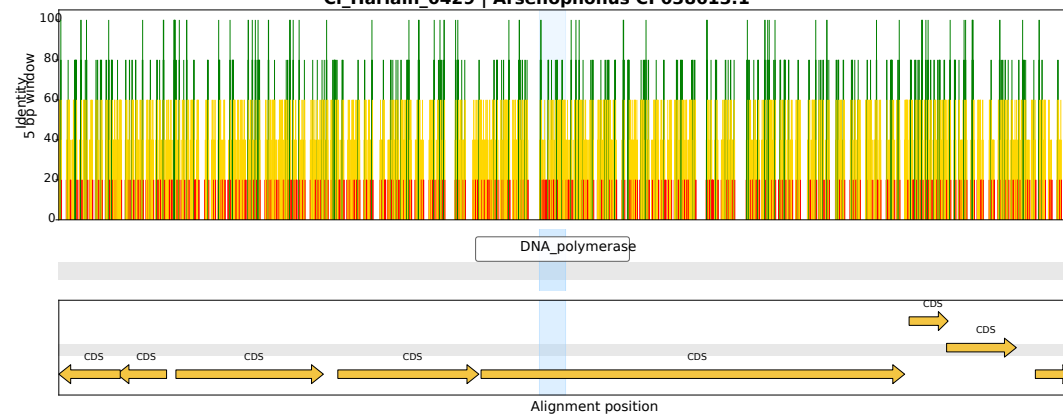

CI\_Harlain\_0202 | *Arsenophonus* CP038613.1

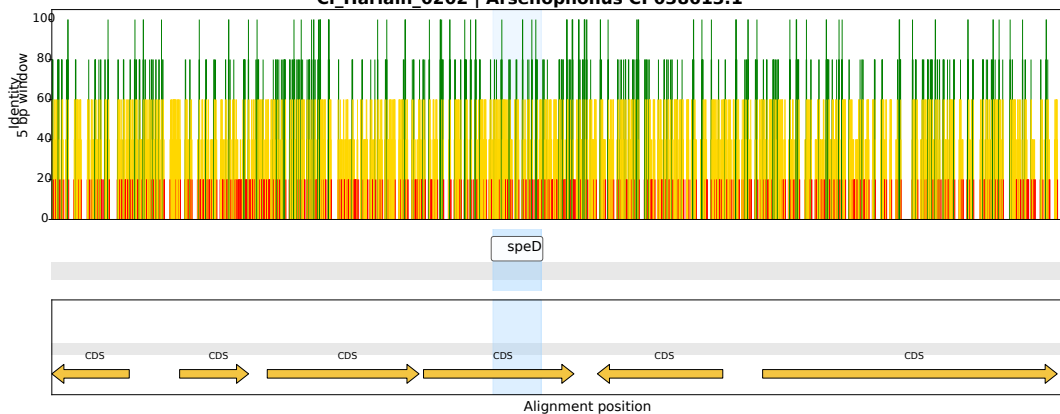

CI\_Harlain\_0280 | *Arsenophonus* OZ026540.1

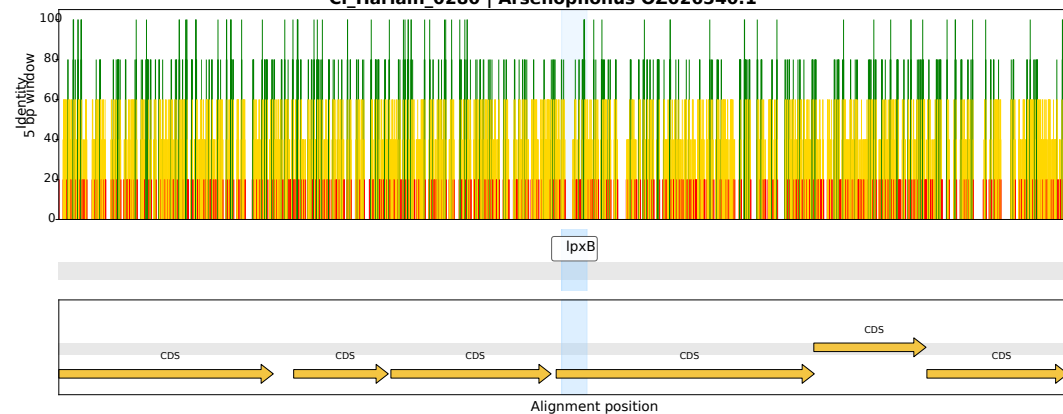

CI\_Harlain\_0304 | *Arsenophonus* CP123759.1

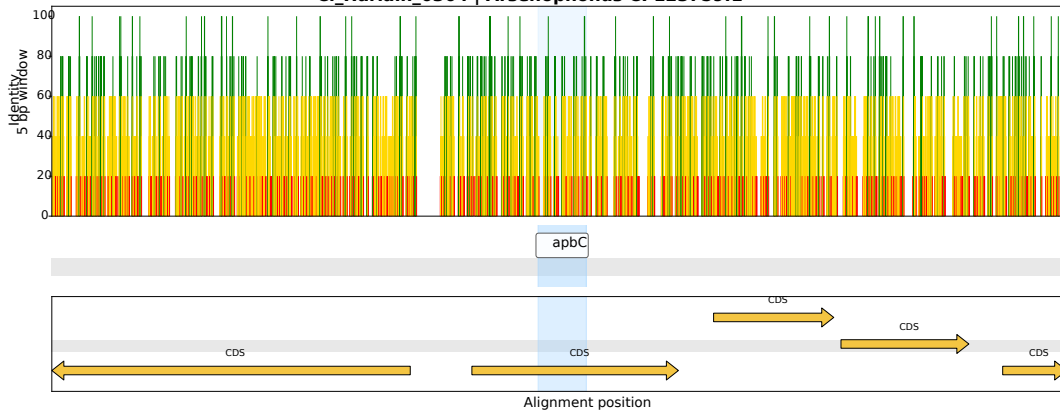

CI\_Harlain\_0147 | *Arsenophonus* CP084222.1

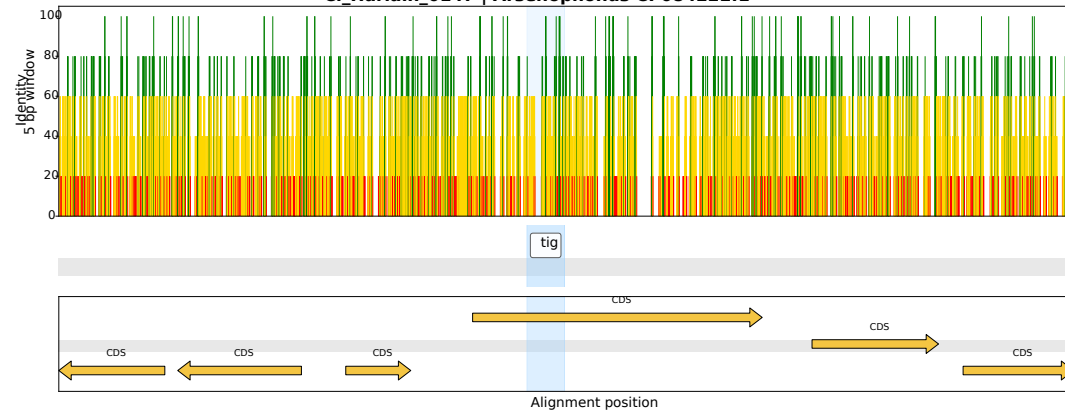

CI\_Harlain\_0398 | *Arsenophonus* CP123499.1

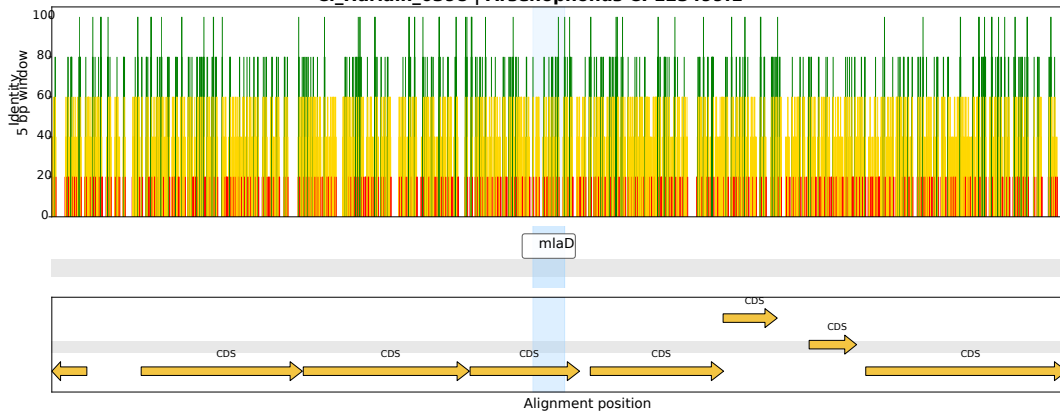

CI\_Harlain\_0098 | *Arsenophonus* OZ026540.1

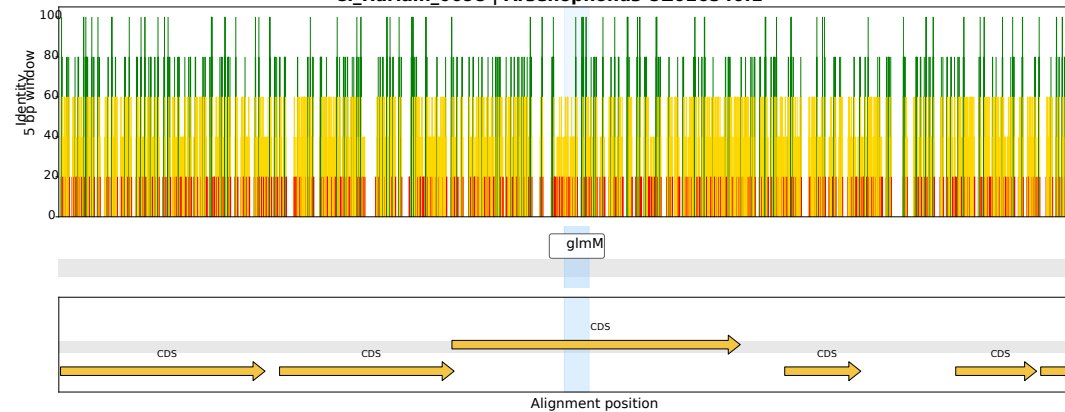

CI\_Harlain\_0179 | *Arsenophonus* CP132903.1

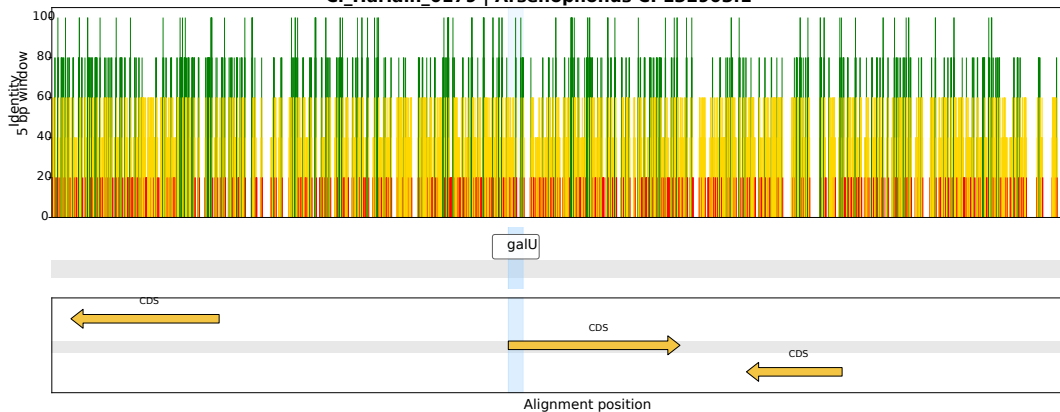

CI\_Harlain\_0007 | *Arsenophonus* CP123759.1

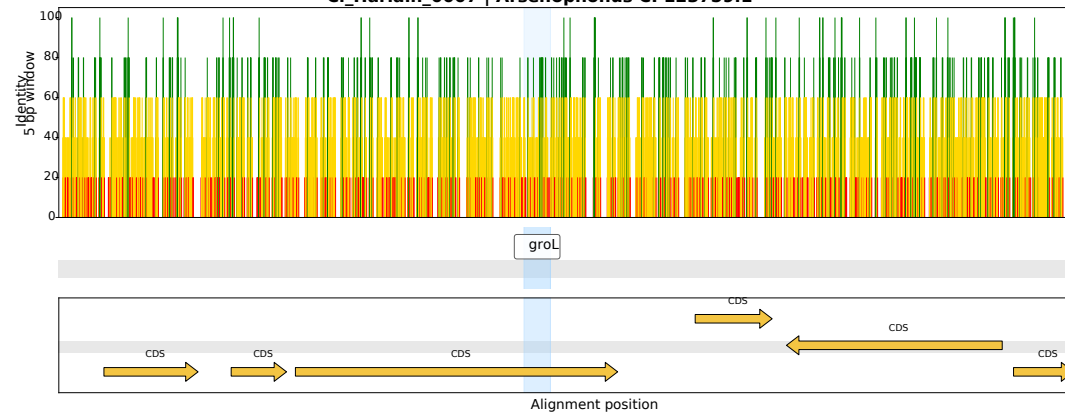

CI\_Harlain\_0215 | *Arsenophonus* CP123499.1

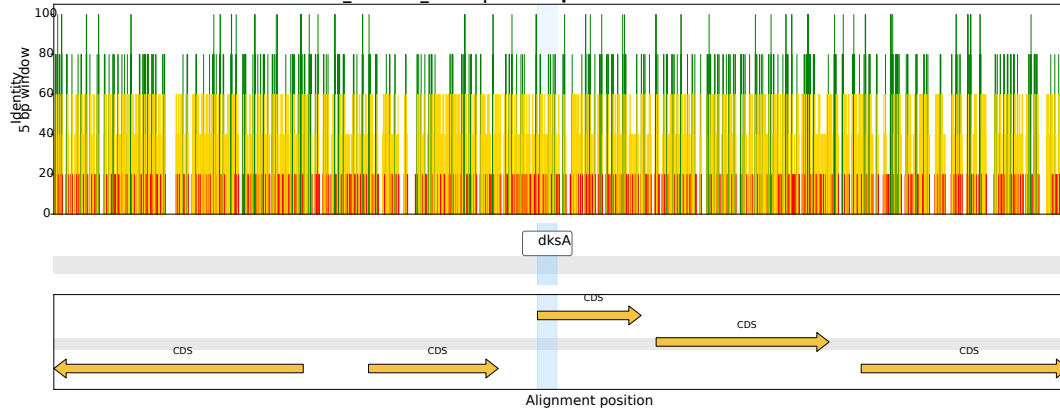

CI\_Harlain\_0117 | *Arsenophonus* OZ026540.1

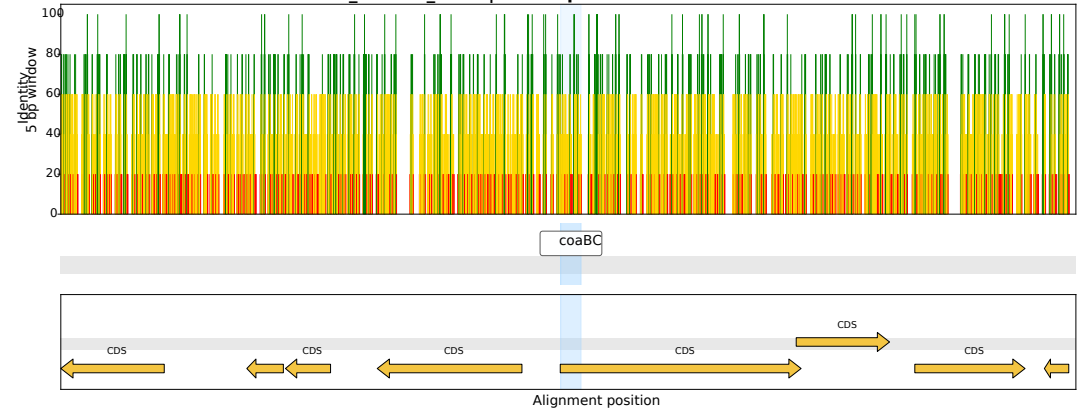

CI\_Harlain\_0323 | *Arsenophonus* CP123759.1

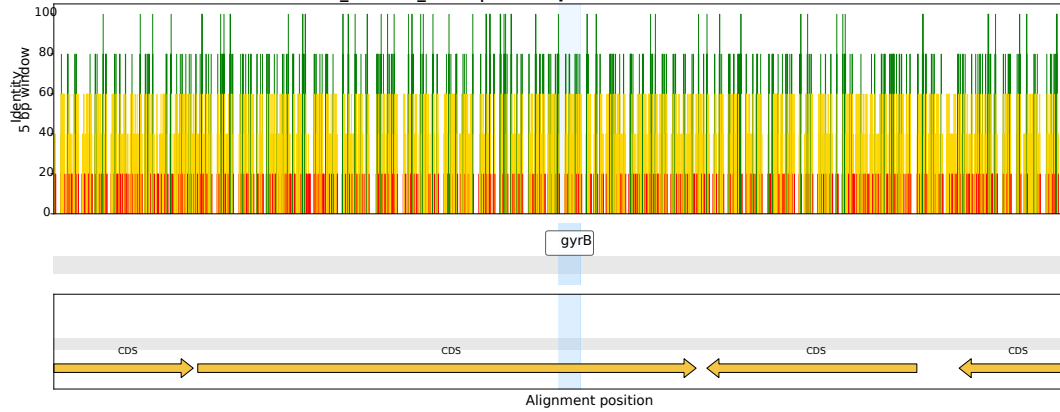

CI\_Harlain\_0386 | *Arsenophonus* OZ026540.1

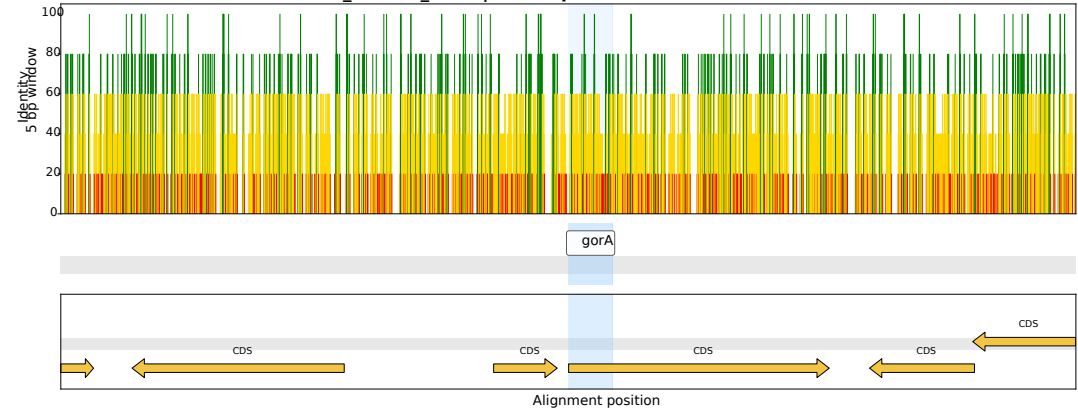

CI\_Harlain\_0059 | *Arsenophonus* OZ026540.1

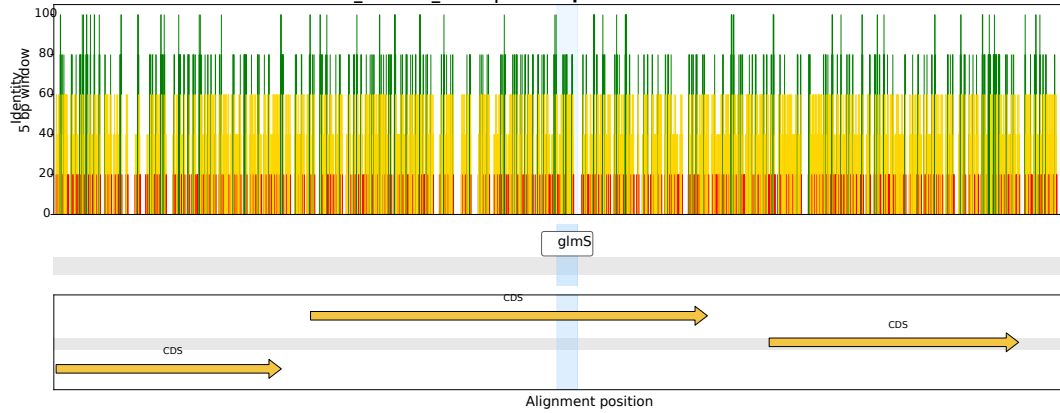

CI\_Harlain\_0258 | *Arsenophonus* LR025108.1

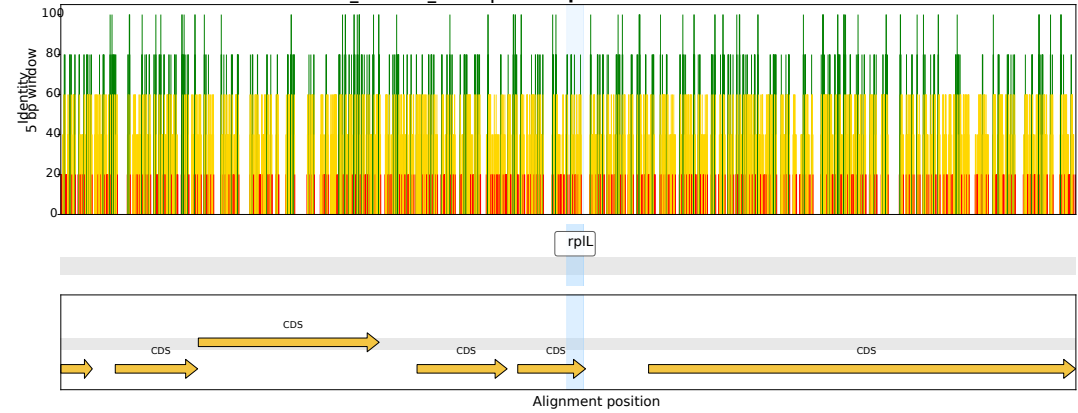

CI\_Harlain\_0050 | *Arsenophonus* CP123499.1

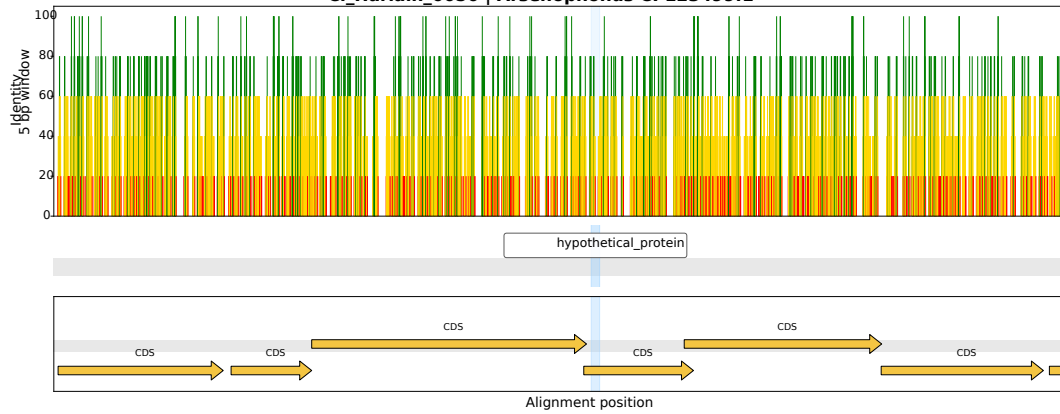

CI\_Harlain\_0303 | *Arsenophonus* LR025108.1

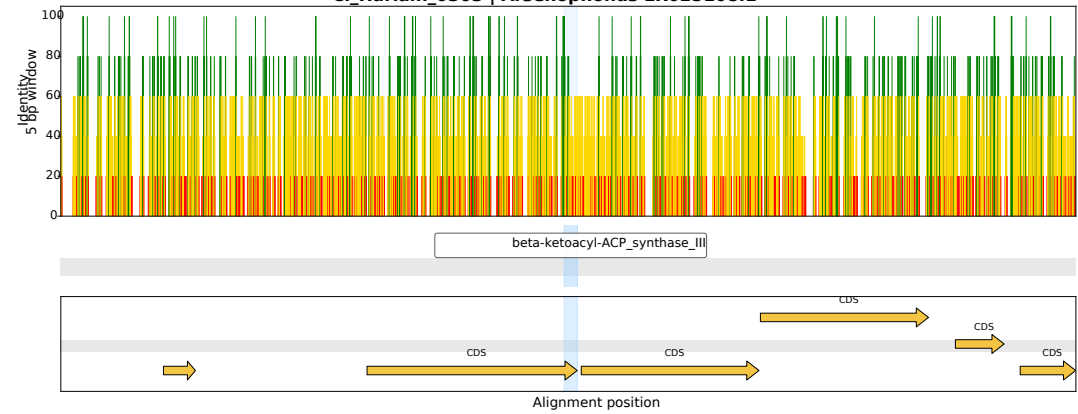

CI\_Harlain\_0182 | *Arsenophonus* CP038613.1

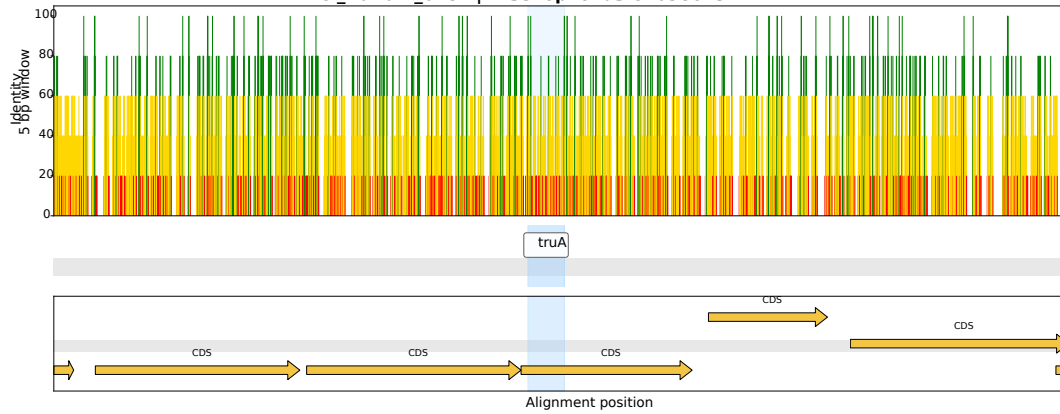

CI\_Harlain\_0092 | *Arsenophonus* OZ026540.1

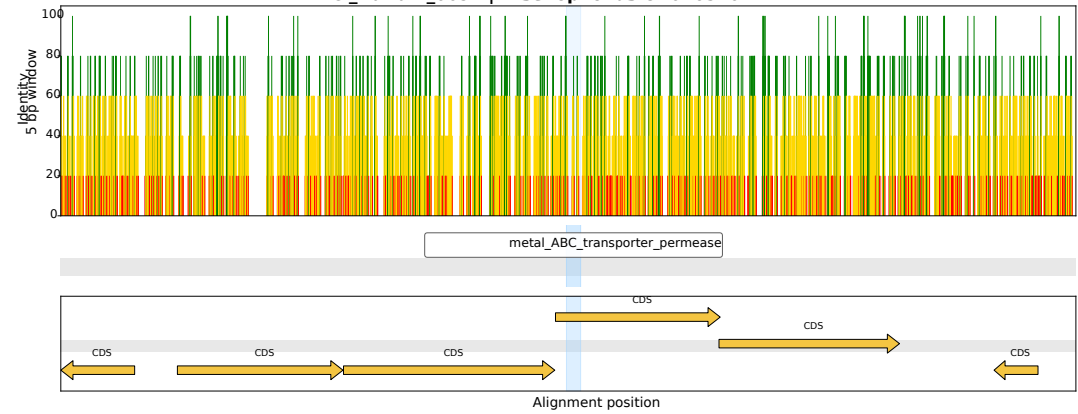

CI\_Harlain\_0456 | *Arsenophonus* CP038155.1

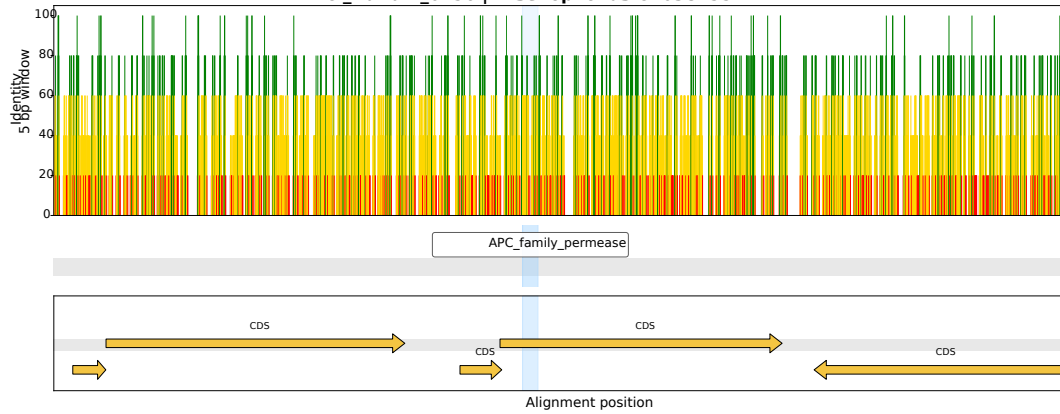

CI\_Harlain\_0356 | *Arsenophonus* CP038614.1

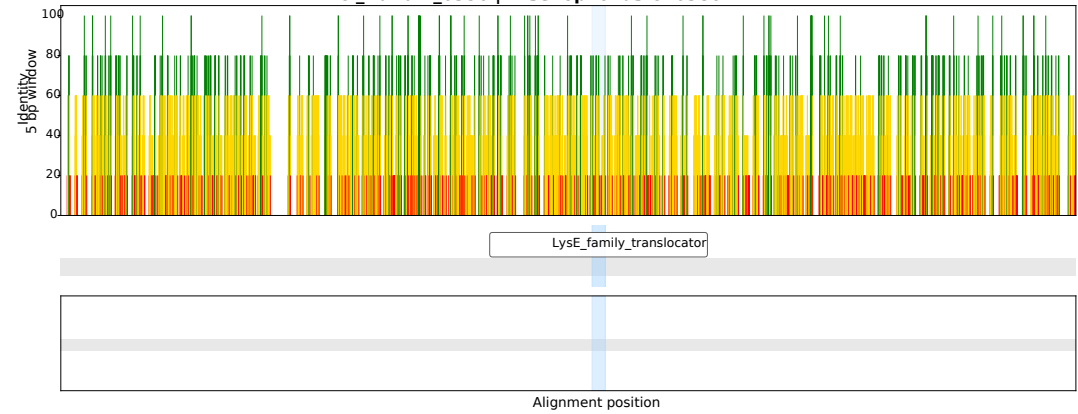

CI\_Harlain\_0444 | *Arsenophonus* CP038155.1

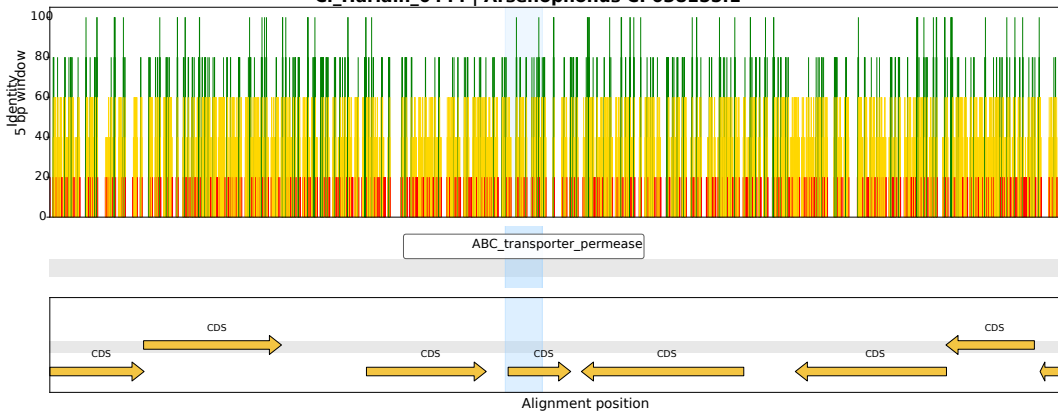

CI\_Harlain\_0401 | *Arsenophonus* CP123492.1

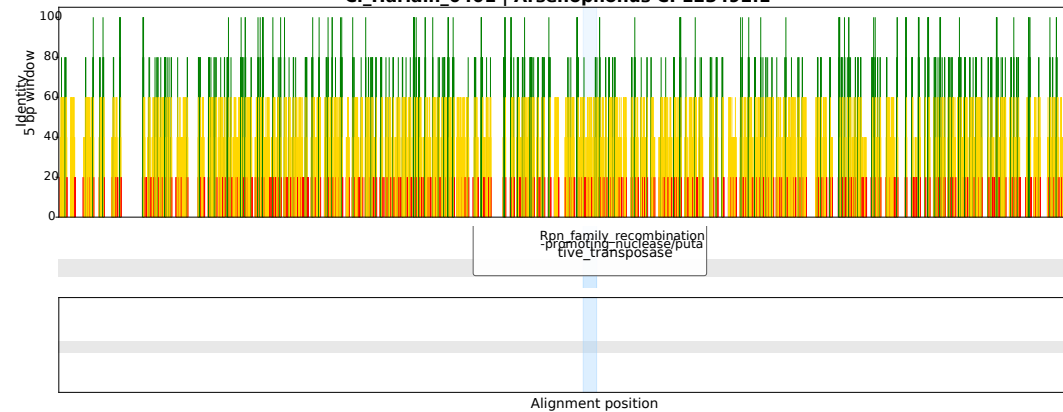

CI\_Harlain\_0345 | *Arsenophonus* LR025108.1

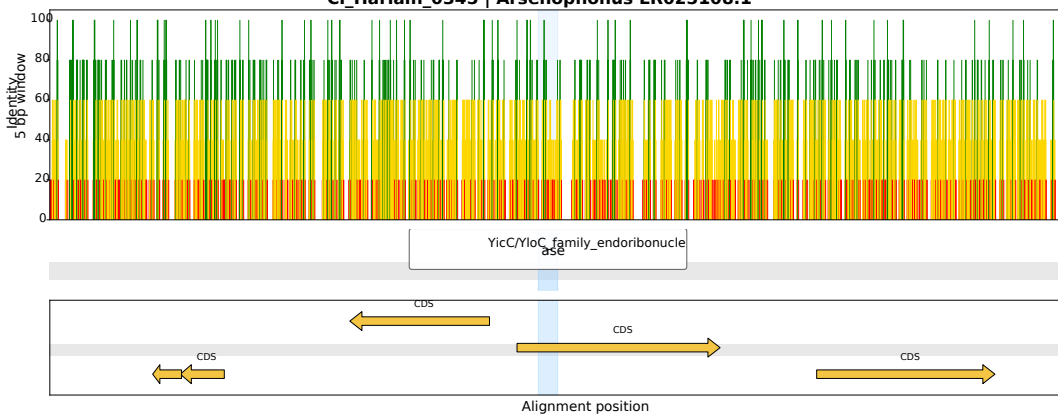

CI\_Harlain\_0211 | *Arsenophonus* OZ026540.1

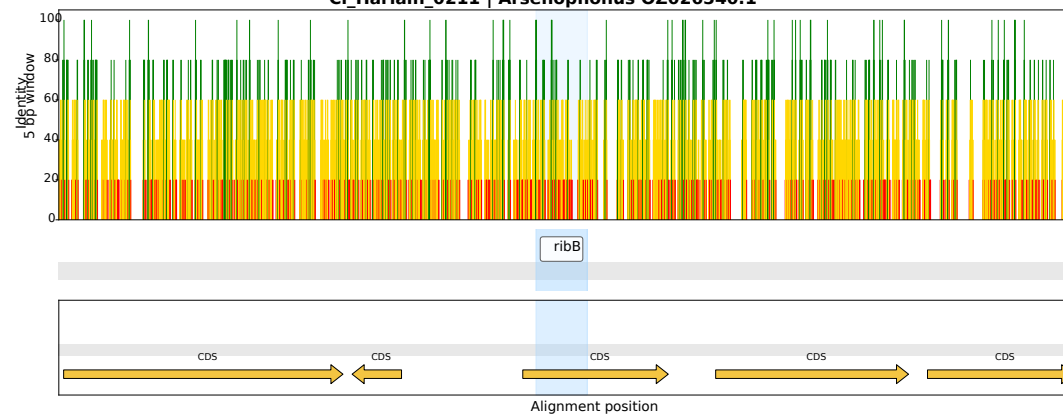

CI\_Harlain\_0326 | *Arsenophonus* OZ026540.1

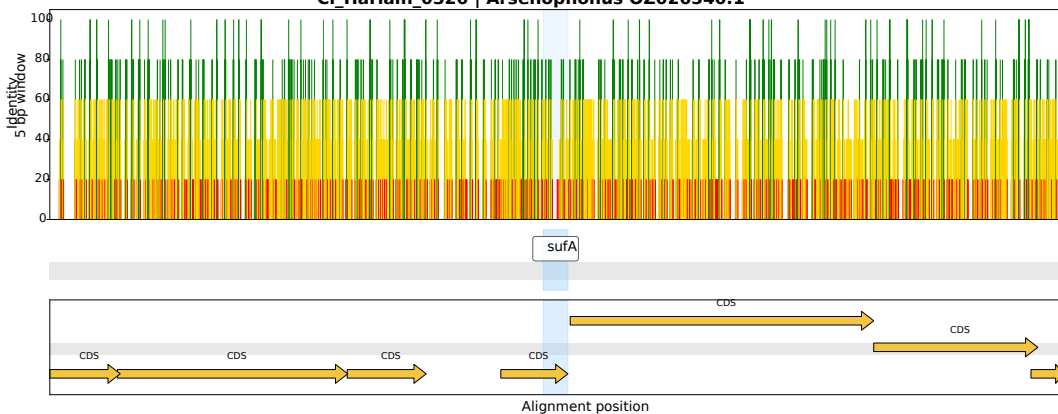

CI\_Harlain\_0388 | *Arsenophonus* CP038155.1

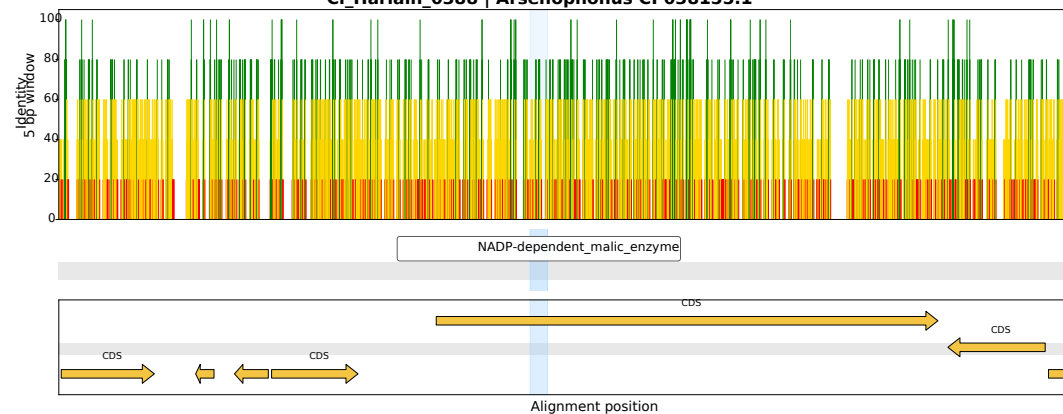

CI\_Harlain\_0313 | *Arsenophonus* CP123759.1

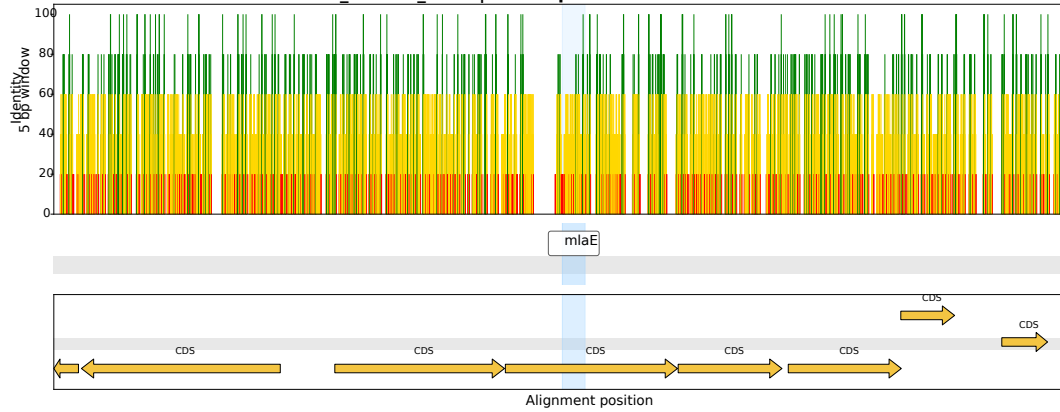

CI\_Harlain\_0088 | *Arsenophonus* CP123523.1

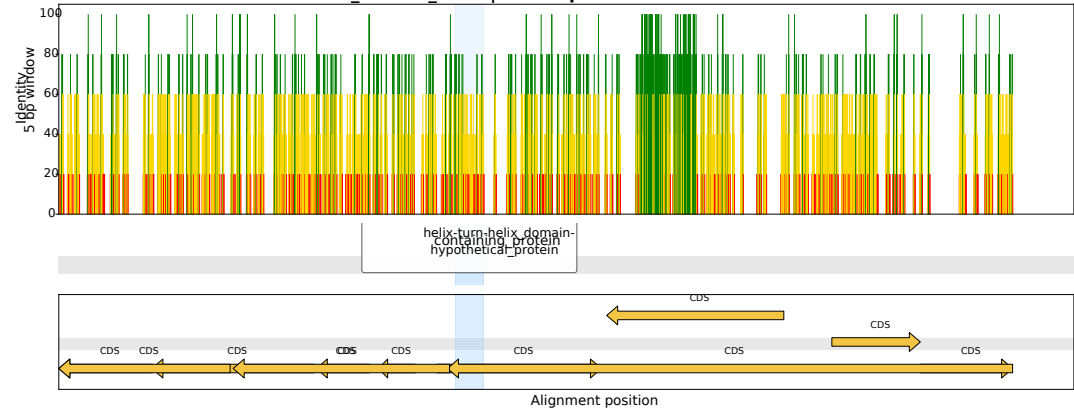

CI\_Harlain\_0468 | *Arsenophonus* CP123499.1

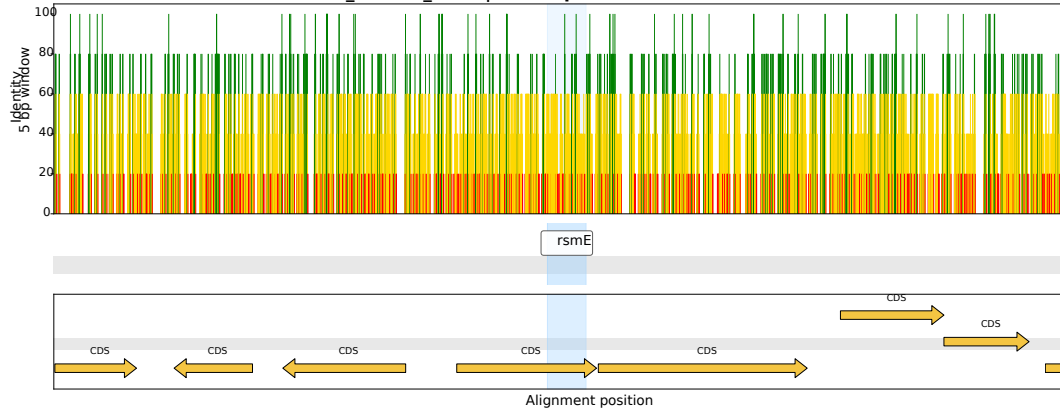

CI\_Harlain\_0281 | *Arsenophonus* CP084222.1

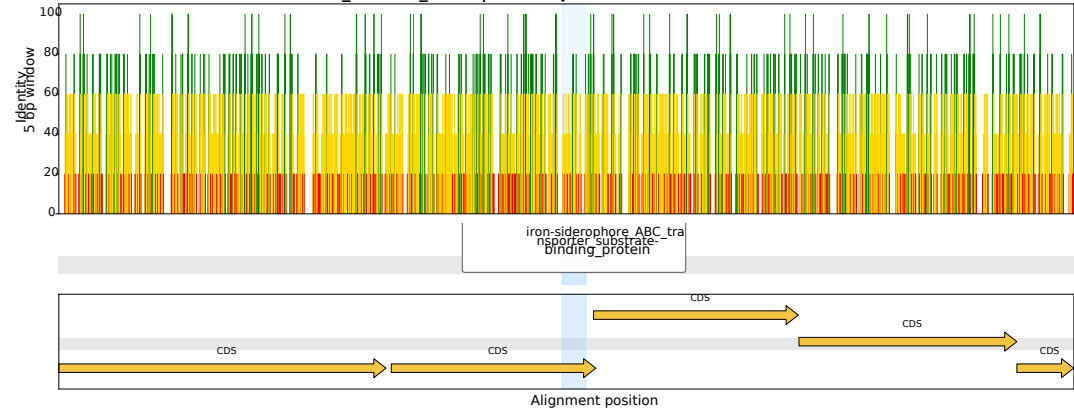

CI\_Harlain\_0458 | *Arsenophonus* CP038613.1

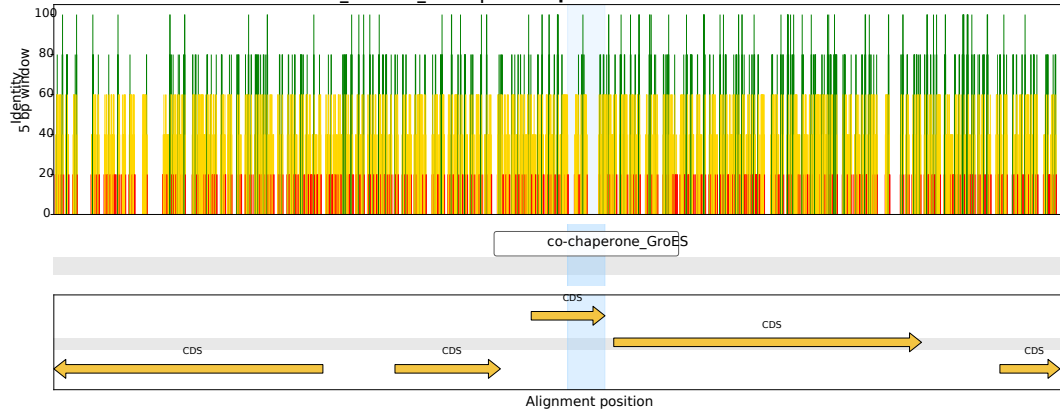

CI\_Harlain\_0174 | *Arsenophonus* CP123499.1

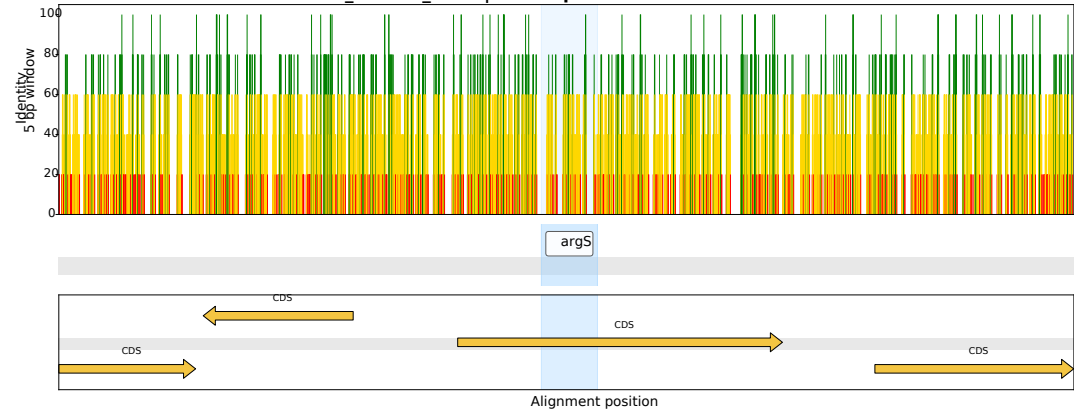

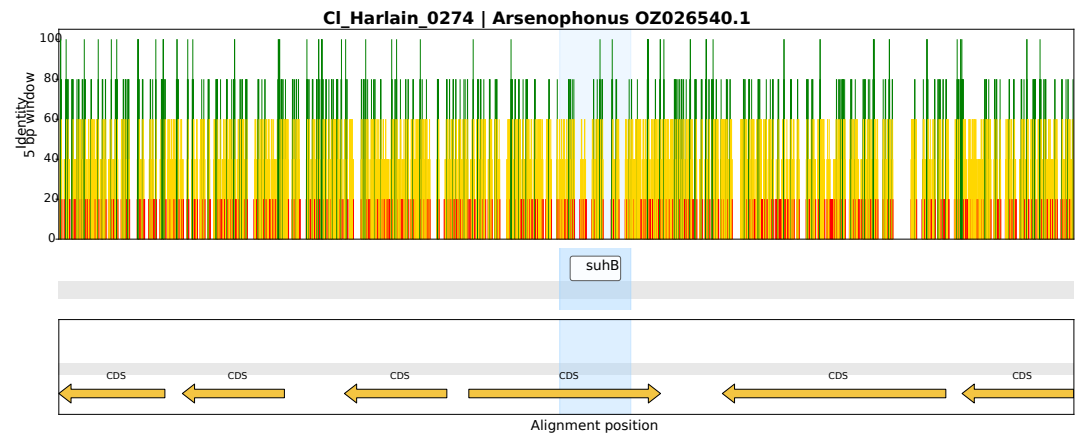
