## Supplementary figure S2 for "*Arsenophonus* ghosts in bed bug genomes: multiple origins and 50 million years of persistence"

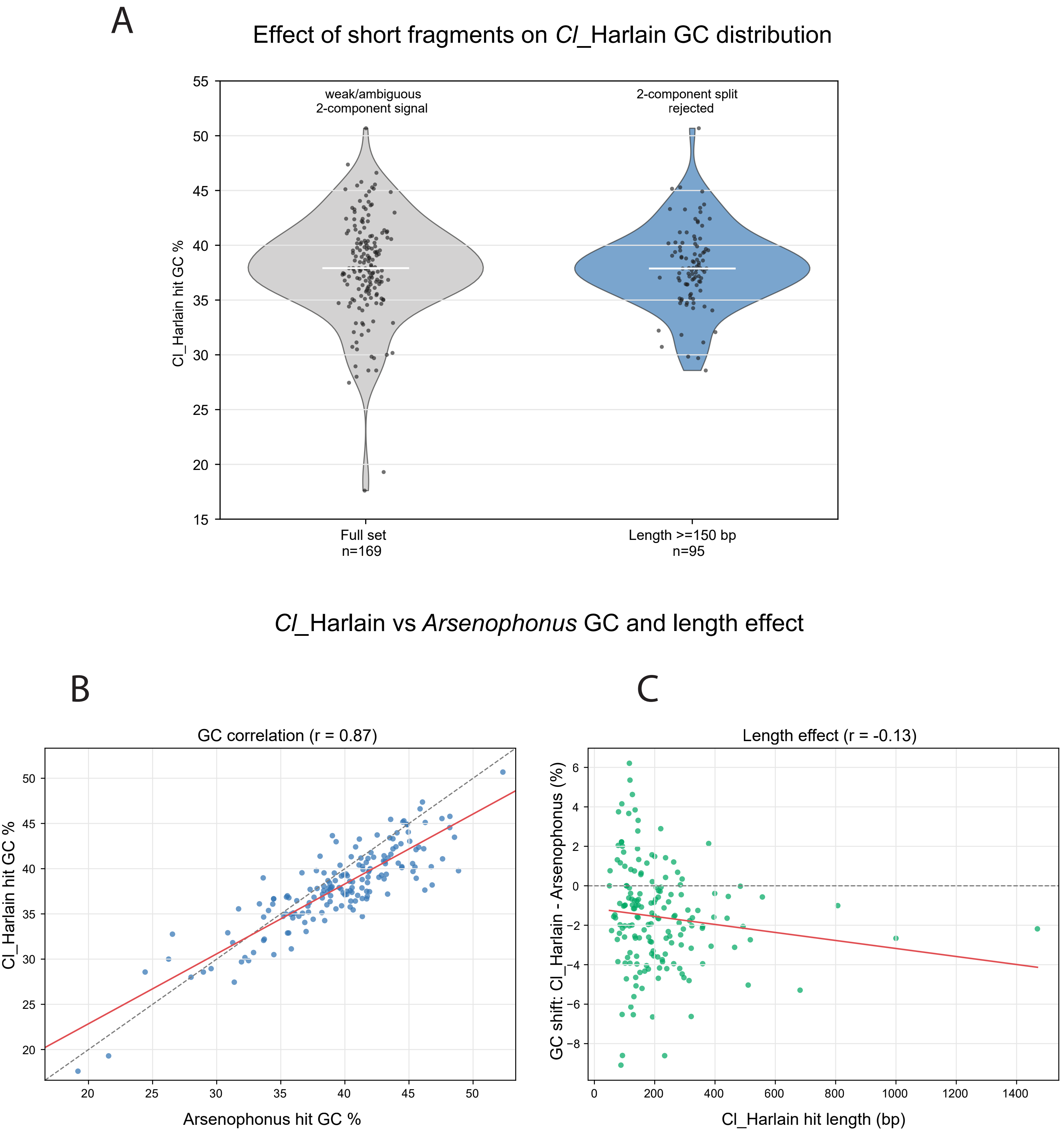


**Supplementary Figure 2**. GC composition of *Cl*_Harlain-associated *Arsenophonus*-derived HGT fragments. (A) Effect of short fragments on the GC distribution of the *Cl*_Harlain-associated HGT set. The complete 169-locus set is centered at approximately 38–39% GC but includes several short low-GC outliers, producing a weak/ambiguous two-component signal. After excluding loci shorter than 150 bp, the remaining 95 loci form a tighter distribution and the two-component split is rejected, indicating that the apparent multimodality is driven mainly by short, compositionally unstable fragments. (B) Relationship between GC content of *Cl*_Harlain HGT fragments and their matched *Arsenophonus* homologous regions. Each point represents one locus. GC values are strongly correlated between the two datasets (Pearson's r = 0.87), indicating retention of the bacterial compositional signal. (C) Relationship between fragment length and GC shift, calculated as *Cl*_Harlain GC minus matched *Arsenophonus* GC. *Cl*_Harlain fragments are on average slightly GC-depleted relative to their *Arsenophonus* matches, but this shift is only weakly associated with fragment length (r = −0.13).
