## Supplementary figure S3 for "*Arsenophonus* ghosts in bed bug genomes: multiple origins and 50 million years of persistence"

A

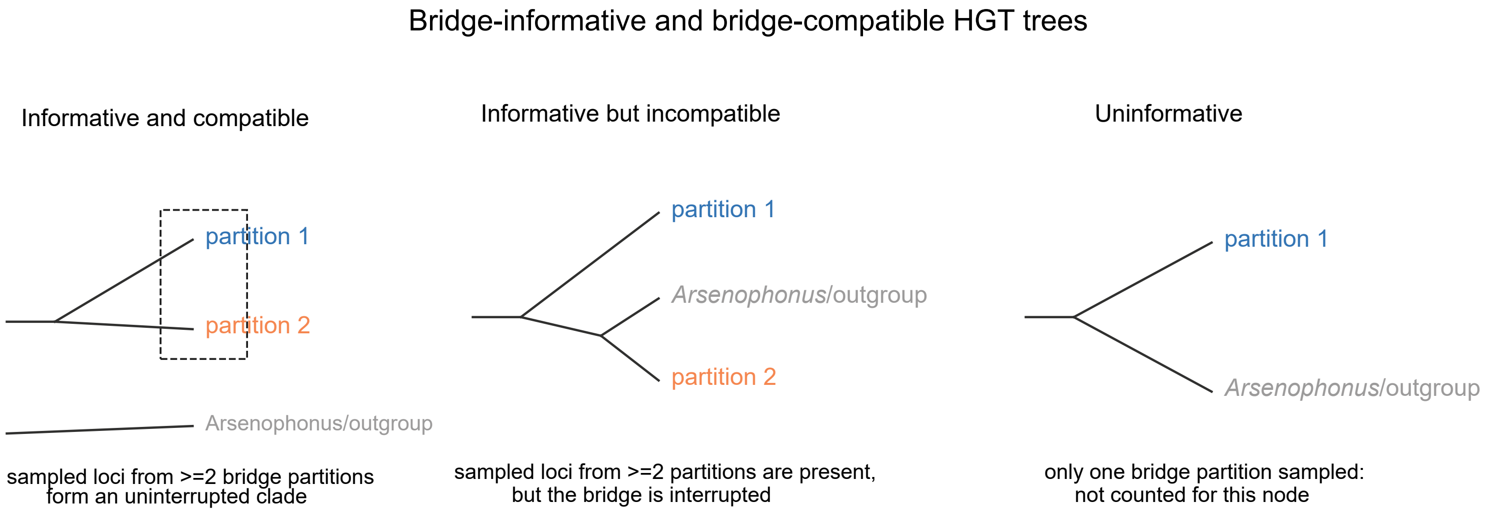


B

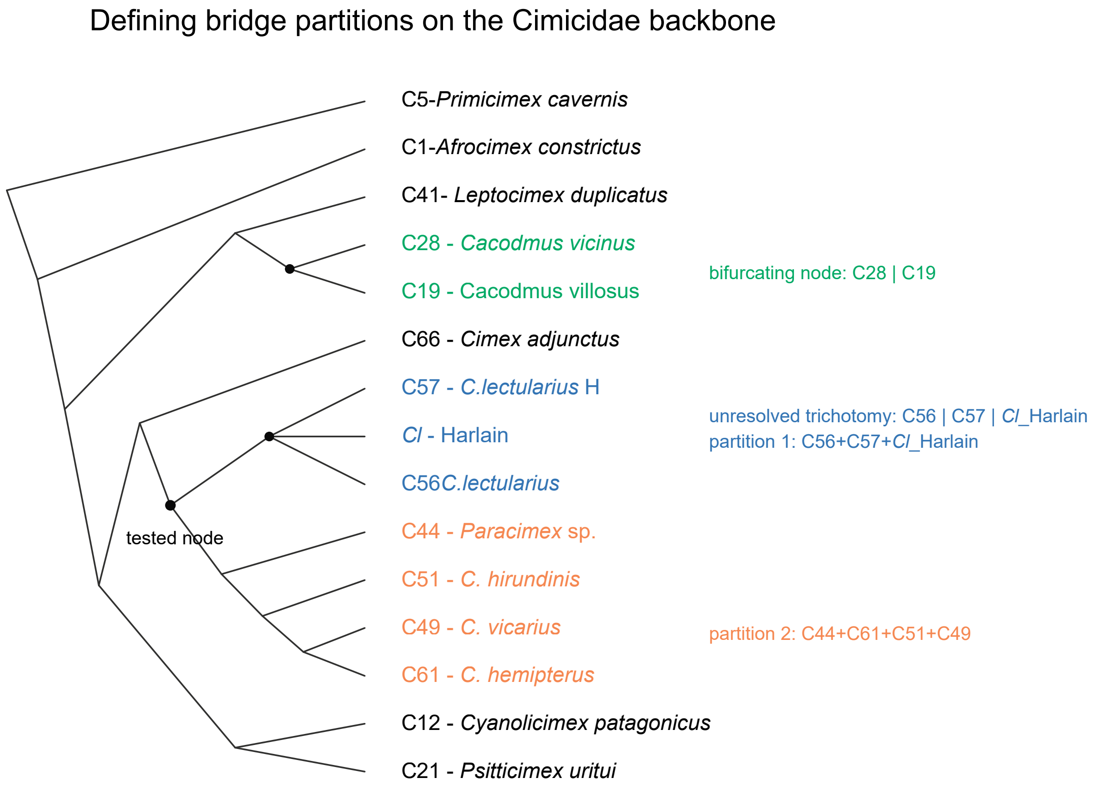


**Supplementary Figure 3**. Bridge-compatibility scoring used to compare individual HGT-locus trees with the Cimicidae backbone phylogeny. (A) Schematic examples of bridge scoring in individual HGT trees. A tree was considered informative for a tested backbone node only when HGT loci from at least two bridge partitions of that node were present. An informative tree was scored as bridge-compatible when the sampled bridge partitions formed an uninterrupted clade. Trees in which *Arsenophonus* or outgroup sequences interrupted the sampled bridge partitions were scored as incompatible, and trees containing loci from only one bridge partition were treated as uninformative for that node. (B) Definition of bridge partitions on the Cimicidae reference backbone. For each tested node, the taxa descending from its immediate child branches defined the bridge partitions. Because the backbone is bifurcating, most tested nodes have two bridge partitions, illustrated by the C28 | C19 node. The unresolved C56+C57+*Cl*_Harlain trichotomy was treated as three partitions. For the core Cimicinae test, bridge partitions corresponded to C56+C57+*Cl*_Harlain and C44+C61+C51+C49.
